# Mathematical modeling of SUMOylation reveals tissue-specific and dynamic responses to abiotic stress

**DOI:** 10.64898/2026.10.07.756741

**Authors:** Kamil Drynda, Jason Banda, Shraboni Ghosh, Lisa Davies, Malcolm J Bennett, Ari Sadanandom, Etienne Farcot, Leah R Band

## Abstract

SUMOylation is an essential and evolutionarily conserved post-translational modification that enables rapid cellular adaptation to environmental stress. Despite its importance, how the SUMOylation machinery controls its dynamics remains poorly understood. Here, we integrated a mathematical model of the plant SUMO cycle with quantitative imaging and biochemical data to decode the regulatory logic of SUMOylation dynamics. The model reveals a fundamental principle: steady-state SUMOylation levels are determined not by the full complement of SUMO-cycle components, but by the abundance of SUMO modifier proteins and E1 enzymes, together with SUMO protease abundance and activity. Strikingly, when we integrated fluorescence measurements from reporters for SUMO-cycle components in *Arabidopsis* roots after three hours of abiotic stress, the model predicted decreased SUMOylation levels across several tissues, contrary to prevailing evidence. Motivated by this unexpected prediction, we performed time-resolved imaging to better characterize the dynamics of key SUMO-cycle components after salt and osmotic stress. These data revealed highly dynamic component levels within the first three hours of stress that were not captured in the previous measurements. Incorporating this temporal data into the model uncovered a twin-peaked SUMOylation response to salt stress: an early surge underpinned by protease dynamics, followed by a decline and a second peak driven by elevated SUMO modifier and E2 levels. Western blot analysis revealed a corresponding twin-peaked pattern of SUMOylation. Together, our findings demonstrate that the plant SUMO cycle is a highly dynamic system that achieves tissue- and stress-specific control of SUMOylation through the coordinated interplay of multiple SUMO-cycle components.

---

Plants have evolved many adaptive mechanisms to perceive and respond to environmental stresses (1–3). Among these adaptive response mechanisms, post-translational modifications (PTMs) are crucial for translating stress stimuli into rapid changes in signaling pathways. PTMs enable plants to rapidly and reversibly respond to changing conditions by modulating protein activity, subcellular localization, protein-protein interactions and stability (4). SUMOylation is an essential PTM that occurs across eukaryotes and plays a significant role in adaptive stress responses (5–9). During SUMOylation, Small Ubiquitin-like Modifier (SUMO) proteins are covalently attached to target proteins via isopeptide bonds, thereby modifying target-protein function (5).

SUMOylation occurs through a series of enzymatic reactions, known as the SUMO cycle (Fig. 1A) (5, 10). The SUMO cycle is similar to the ubiquitination system; both are dynamic and reversible PTM systems that regulate thousands of proteins and involve enzymes with similar structures (11–13). Unraveling the mechanisms regulating PTMs can be challenging due to the large number of components involved; for instance, there are thousands of proteins involved in ubiquitination. In contrast, the SUMO cycle has only 32 components, making SUMOylation much more tractable to study than other PTMs (14).

**Fig 1.**
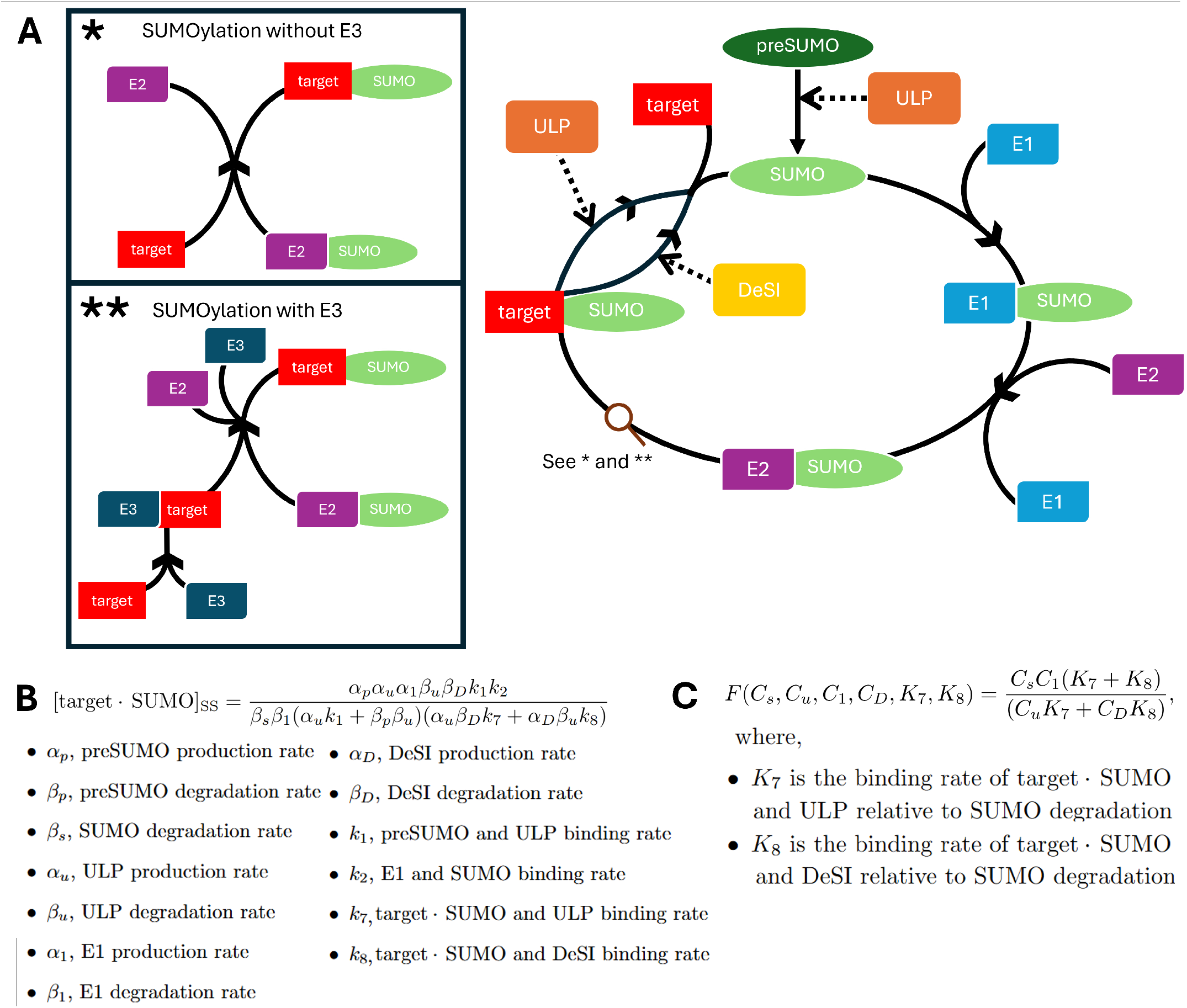
Increased SUMOylation of target proteins after stress arises from coordinated changes in the SUMO cycle. (**A**) Schematic of the SUMO cycle in *Arabidopsis*, highlighting the two routes by which SUMOylated target can form (* and **). E4 is omitted, as polySUMOylation is not included in the mathematical model. (**B**) Steady-state expression for the level of SUMOylated target (see SI Appendix, Section 1.2). (**C**) Closed-form expression for the steady-state fold change of SUMOylated target in response to stress, *F*, as a function of the fold changes in SUMO (*C*^*s*^), ULP (*C*^*u*^), E1 (*C*^1^), and DeSI (*C*^*D*^), together with the parameters *K*^7^ and *K*^8^ (see SI Appendix, Section 2.1 for the model derivation).

The SUMO cycle (Fig. 1A) starts through maturation of preSUMO through ULP-type proteases, which cleave the C terminus of the SUMO protein, generating mature SUMO (5). Mature SUMO is activated by the E1 enzyme, a heterodimer consisting of SAE1a/b and SAE2, forming the E1 *·* SUMO complex (5). The activated SUMO is then transferred to the E2 conjugating enzyme (SCE1) to form the E2 · SUMO complex (5). Target proteins can be SUMOylated in two ways: either by the E2 · SUMO complex binding directly to the target protein, or through binding of the target protein to an E3 ligase, which catalyzes its interaction with the E2 · SUMO complex (15). An additional step, catalyzed by the E4 ligases, forms poly-SUMO chains (5). Finally, ULP and DeSI-type proteases remove SUMO from SUMOylated target proteins, in a process termed deSUMOylation. This process releases the target protein and replenishes the pool of free SUMO, for reuse in future SUMOylation cycles. DeSUMOylation is the sole function of the DeSI-type protease (16), whereas the ULP-type protease has a dual function in facilitating both deSUMOylation and SUMO maturation (5).

To investigate how the components of the SUMO cycle respond to different environmental stresses and thereby modify the SUMOylation of target proteins, Banda et al. developed the SUMO Cell Atlas (14). As many components of the SUMO cycle are encoded by multigene families, this resource characterized isoform-specific abundance and tissue-specific expression patterns in *Arabidopsis thaliana* roots under salt stress, osmotic stress and treatment with the biotic elicitor flagellin.

The SUMO Cell Atlas combines measurements of transcript abundance, protein abundance, and subcellular localization for all 32 known SUMO-cycle components across key root tissues (epidermis, cortex, endodermis, and stele) and developmental zones (meristem and elongation zone). It revealed that the SUMO cycle components are present across these major root tissues and developmental zones, but their abundance and localization vary between tissues. These measurements indicate that the SUMOylation machinery is not uniformly distributed throughout the root and that specific components may contribute to the regulation of SUMOylation in different tissues.

In this study, we focus on the two abiotic stresses examined in the SUMO Cell Atlas: salt and osmotic stress, which induced distinct, tissue-specific changes in the abundance of SUMO pathway components. In particular, salt-stress responses of the SUMO cycle appeared to activate epidermal responses, whereas osmotic stress predominantly affected SUMO-cycle components in the inner root tissues. Furthermore, the Cell Atlas revealed that SCE1 is the only SUMO component that increased significantly under both stress conditions, suggesting that it may play a central role in driving stress-induced SUMOylation (reviewed (17)) and distinct responses among SUMO proteases to the different stresses.

The Cell Atlas demonstrates that stress-induced regulation of the SUMO cycle is spatially heterogeneous and involves coordinated changes across multiple components of the pathway (14). However, experimental observations alone cannot determine how these changes combine to mediate dynamic changes in cellular SUMOylation levels. Here, we developed a mathematical model to understand how the abiotic-stress-induced changes in SUMO-cycle components combine to regulate SUMOylation levels. The modeling revealed that only a subset of the components determines the steady state. Incorporating the SUMO Cell Atlas data, the model predicted that the steady-state SUMOylation response varied across tissues and in several cases contradicted the expected stress-induced increase in SUMOylation seen in previous studies (5, 16, 18, 19).

Given the steady-state predictions were inconsistent with our expectations, we generated image time courses of key SUMO-cycle components during salt and osmotic stress responses. These data revealed that the SUMO-cycle components exhibit highly dynamic responses to stress. Integrating these dynamic measurements, the model predicted stress- and tissue-specific SUMOylation dynamics governed by the balance between SUMOylation, driven by free SUMO and E2, and deSUMOylation, mediated by DeSI and ULP. The model further revealed that E2 drives rapid SUMOylation responses without changing the steady-state level. Early changes in deSUMOylation are shown to rapidly alter the abundance of SUMOylated target whereas increases in SUMO and E2 must propagate through the cycle before influencing SUMOylation. Together, these temporally coordinated effects were predicted to generate transient dynamics, including a twin-peaked response to salt stress in the stele. Western blot analysis subsequently detected a corresponding twin-peaked pattern of SUMO-conjugate accumulation in salt-treated roots, consistent with the predictions. These results imply that the stress-dependent drivers of SUMOylation differ between tissues and reveal how SUMOylation dynamics arise from the coordinated interplay of multiple SUMO-cycle components rather than any single rate-limiting step.

## Results

### Mathematical modeling of the SUMO cycle in *Arabidopsis thaliana*

To model SUMOylation, we represented the SUMO cycle as a series of chemical reactions describing the interactions between families of SUMO-cycle components. Applying the law of mass action yielded a system of ordinary differential equations (ODEs) governing the dynamics (SI Appendix, Section 1.1). The model predicted the concentrations of the following molecular species: SUMO isoforms in their immature (preSUMO) and mature (SUMO) forms; the functional E1 complex; the E2, E3, ULP, and DeSI enzymatic families; free target protein; and the complexes E1 *·* SUMO, E2 *·* SUMO, E3 *·* target, and target *·* SUMO. PolySUMOylation by E4 enzymes was excluded from our model due to the combinatorial complexity of SUMO chain formation and their diverse molecular consequences (5). To maintain tractability, we further assumed that none of the enzymatic steps are saturated. The target protein was modeled as a generic protein, rather than a specific SUMO target, to understand general SUMOylation dynamics.

Assuming that stress-induced changes in preSUMO, ULP, E1, E2, E3, and DeSI arise through changes in their production rates, we nondimensionalized the system to obtain a system of ODEs describing the fold change of each modeled species (see SI Appendix, Section 1.3). The resulting model depends on 25 parameter groupings: 6 correspond to ratios of stressed to unstressed production rates; 15 represent ratios of degradation, binding, and dissociation rates relative to the rate of SUMO degradation; and the remaining 4 describe additional ratios between the concentrations of modeled species prior to the stress application. We note that only the production-rate ratios are directly informed by the experimental data, whereas all other parameters are unknown.

Key features of the model include its cyclical structure, the dual role of ULP, and the two distinct routes for SUMOylated target formation (as shown in the inserts in Fig. 1A). Our model takes inspiration from previous SUMO-cycle modeling studies (20–22); although we note that these previous works focused on human systems, whereas our model describes the SUMO cycle in *Arabidopsis thaliana*, incorporating both the E3 ligases and the DeSI protease, which were not represented in previous models.

### Model predictions reveal key drivers of steady-state SUMOylation levels

To incorporate the SUMO Cell Atlas data (14) into the model and predict how SUMOylation of target protein changes under stress, we had to consider how the measurements were reported. The data consist of measurements taken before and after three hours of stress. As SUMOylation occurs rapidly upon stress onset (23, 24), we assumed that the system is at steady state before and after stress.

Furthermore, the SUMO Cell Atlas (14) provided isoform-level fold changes for SUMO, ULP, E1, E2, E3, and DeSI. To incorporate these data into our model, we aggregated isoform-level measurements into a single fold change for each modeled component. For ULP, E2, E3, and DeSI, this was achieved using fluorescence-derived proportions measured in the root meristem (see SI Appendix, Section 2.2, Table S3).

In contrast, as the E1 enzyme is a dimer, its fold change was calculated accounting for the fold changes in each component of the dimer (see SI Appendix, Section 2.2.3).

Since SUMO2 is the only SUMO family member present in all root-tip tissues (14), we prescribed SUMO changes using the observed fold change of SUMO2. However, the fluorescent protein reporter (14) does not distinguish between preSUMO and mature SUMO. Thus, free SUMO was assumed to exist predominantly in its mature form, consistent with observations in mammalian cells (25) and yeast (26), where the proteases responsible for SUMO maturation are constitutively active (27). As AtSUMO1/2 is a functional ortholog of mammalian SUMO2/3 (5), we expected this to hold in plants. This assumption means that although ULP contributes both to preSUMO maturation and deSUMOylation, in the steady-state model changes in ULP only influenced deSUMOylation.

Under these assumptions, the model enabled us to predict how SUMOylation changes after stress, specifically providing an explicit expression for the steady-state concentration of target protein bound to SUMO, [target SUMO]SS. (Fig. 1B). This expression revealed that the steady-state level of SUMOylated target protein is determined by a subset of the model parameter values, namely the production, degradation, and binding rates of preSUMO, SUMO, E1, ULP, and DeSI, while being independent of the E2- and E3-associated rate parameters.

From this expression, we derived a closed-form relation linking the steady-state fold change in SUMOylated target protein following stress, F, to the fold changes in SUMO, ULP, E1, and DeSI,(*Cs, Cu, C*1, *CD*) (Fig. 1C; derivation in SI Appendix, Section 2.1). This relation showed that in addition to the fold changes in these SUMO-cycle components, the steady-state fold change of SUMOylated target protein also depends on two rate constants, *K*7 and *K*8, which denote the relative rates of ULP- and DeSI-mediated deSUMOylation respectively.

Since the deSUMOylation rates *K*7 and *K*8 are unknown, we explored the range of steady-state responses predicted by the model over a broad range of *K*7 and *K*8 values. We found that changes in SUMO or E1 directly and proportionally affect the steady-state fold change of SUMOylated target (Fig. 1C). In contrast, increasing either DeSI or ULP (with the other components remaining fixed) reduces the steady-state fold change, although the magnitude of this reduction depends on the deSUMOylation rates(SI Appendix, Fig. S1). Differences in the fold changes *Cu* and *CD* alter the relative impact of ULP- and DeSI-mediated deSUMOylation on the stress-induced change in SUMOylation, with the extent of this effect determined by the deSUMOylation rates. Thus, increasing the absolute difference between them (*Cu CD*) leads to a wider range of possible steady-state SUMOylation responses across the deSUMOylation-rate range examined (SI Appendix, Section 2.4, Fig. S2). Moreover, a combined increase in SUMO and E1 amplifies this variability, whereas a combined decrease reduces it (SI Appendix, Section 2.4, Fig. S3).

Taken together, these results provide a quantitative relationship linking experimentally observed changes in SUMO-cycle components to predicted changes in SUMOylated target-protein levels after stress. The model shows that steady-state SUMOylation is governed by the balance between SUMOylation and deSUMOylation, with changes in SUMO and E1 directly scaling the response, while changes in ULP and DeSI introduce parameter-dependent variability through the unknown deSUMOylation rates. Consequently, the model enabled us to predict tissue-specific and stress-specific responses using the SUMO Cell Atlas data.

### Modeling predicts varied SUMOylation responses across root tissues and developmental zones after abiotic stress

Using the SUMO Cell Atlas data, we calculated fold changes for each modeled SUMO-cycle component, revealing differences between salt and osmotic stress and across root tissues (Fig. 2A, C, SI Appendix, Tables S1, S2). Under salt stress, both free SUMO and DeSI levels decreased or changed minimally across all root tissues examined. In contrast, under osmotic stress (Fig. 2C) both free SUMO and DeSI levels increased across all root tissues examined. Under both stresses, E1 and ULP-type proteases displayed tissue- and zone-specific responses, with both increases and decreases observed (Fig. 2B, C). These observations present a complex picture, with distinct tissue-specific and stress-specific responses. Furthermore, in every tissue, the observed changes would act in opposing directions on SUMOylation, making their net effect on SUMOylated target-protein levels difficult to anticipate and highlighting the need for a mathematical model that integrates their combined effects.

**Fig 2.**
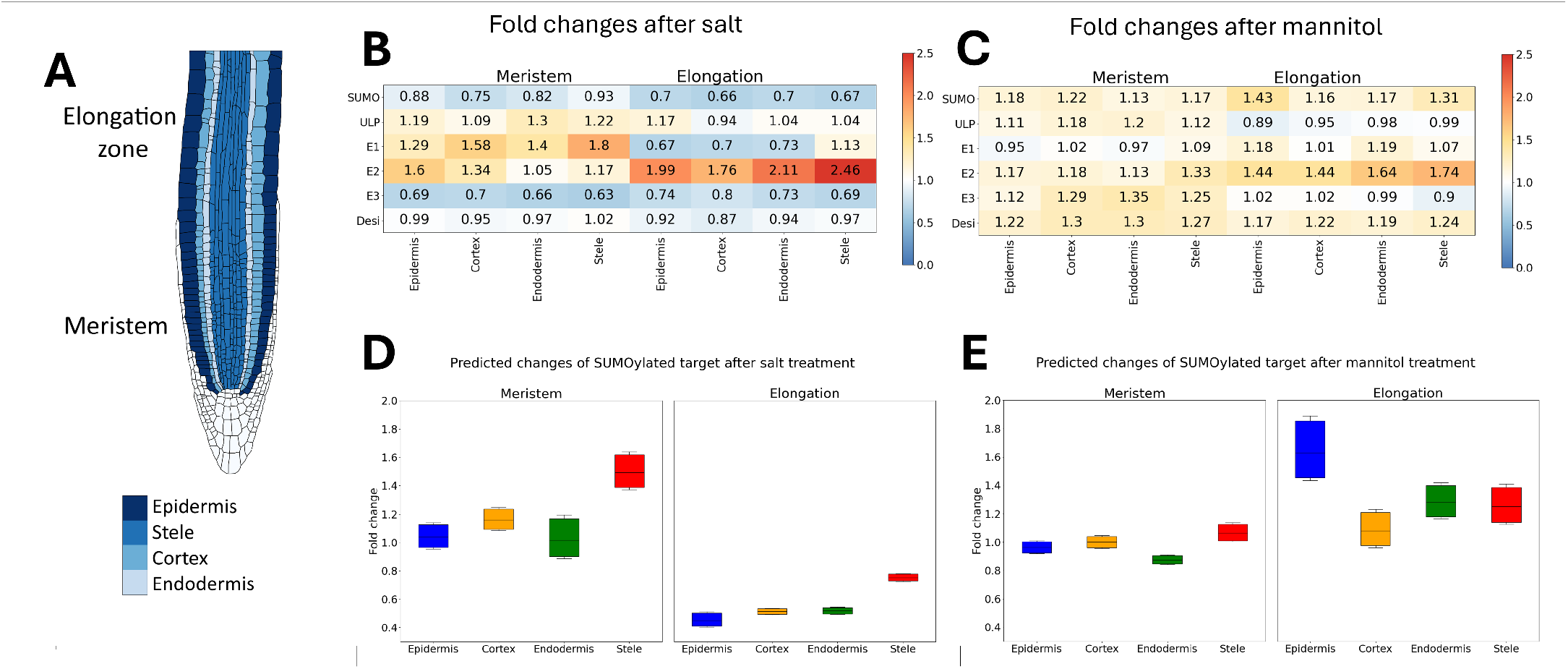
Model predictions reveal how SUMOylated target levels change across root tissues after abiotic stress. (**A**) Schematic of the *Arabidopsis* root tip showing the meristem and elongation zones and the epidermis, cortex, endodermis, and stele cell types. (**B**,**C**) Amalgamated fold changes of SUMO-cycle-component families after salt and osmotic stress, respectively, computed using fluorescence-derived proportions (SI Appendix, Table S3) and the SUMO Cell Atlas data ((14), SI Appendix, Tables S1 and S2). (**D**,**E**) Predicted steady-state fold changes of SUMOylated target after salt (**D**) and mannitol (**E**) treatment, computed using the steady-state fold change of SUMOylated target in Fig. 1C and the fold-change data in panels (**B**) and (**C**). In all simulations, *K*^7^ and *K*^8^ were varied over the range [1*/*100, 100] using 10,000 parameter sets sampled from log space as described in SI Appendix, Section 2.3. These results agree with the analytical bounds discussed in SI Appendix, Section 2.4.

To determine how these observed changes combine to affect SUMOylation, we incorporated the fold changes derived from the SUMO Cell Atlas data (Fig 2B,C; SI Appendix, Tables S1,S2) into the steady-state expression for SUMOylated target protein (Fig. 1C). Previous studies suggest that SUMOylation increases following stress (5, 16, 18, 19); however, our model predicted distinct changes in SUMOylation across *Arabidopsis* root cell types and developmental zones following salt and osmotic stress (Fig. 2D,E). The model predicted elevated SUMOylated target-protein levels after salt stress in the meristematic cortex and stele (Fig. 2D), with the strongest increase in the stele, where E1 upregulation outweighs the opposing effect of increased ULP-mediated deSUMOylation (SI Appendix, Fig. S4). In contrast, predicted responses in the meristematic epidermis and endodermis depended on the deSUMOylation rates, with both increases and decreases in SUMOylated target-protein levels predicted across the parameter range examined. In the elongation zone, however, concurrent reductions in SUMO and E1 resulted in a consistent decrease in SUMOylated target-protein levels across all tissues (Fig. 2B,D).

Under osmotic stress, the model predicted minimal changes in SUMOylated target-protein levels in the meristematic epidermis and cortex, where increased SUMO largely offsets the opposing effects of elevated ULP and DeSI (Fig. 2E; SI Appendix, Fig. S5). In the endodermis, the increase in SUMO was insufficient to compensate for increased deSUMOylation, resulting in a consistent decrease across all deSUMOylation rates examined. Conversely, simultaneous increases in SUMO and E1 in the stele produced a modest but consistent increase in SUMOylation. In the elongation zone under osmotic stress, the combined increases in SUMO, E1, and DeSI resulted in more variable predictions (Fig. 2C,E). SUMOylated target-protein levels increased in the epidermis, endodermis, and stele across the range of deSUMOylation rates examined, although the magnitude of the response varied with these rates. In the cortex, however, the increase in SUMO and decrease in ULP were insufficient to consistently offset the increase in DeSI, resulting in both increases and decreases in SUMOylation depending on the deSUMOylation rates (Fig. 2C,E).

Taken together, these results demonstrate that steady-state fold changes in SUMOylated target-protein level are stress- and tissue-specific. Although previous studies indicate that SUMOylation increases following abiotic stress, the model predicted that this response is not uniform across the *Arabidopsis* root zones and tissues. Instead, this response depends on the combined effect of changes in multiple SUMO-cycle components resulting in SUMOylated target-protein levels that may increase, decrease, or remain largely unchanged. While increases in SUMO and E1 promote SUMOylation, the opposing effects of increased ULP and DeSI introduce substantial deSUMOylation rate-dependent variability into the predicted response, which combine nonlinearly with changes in SUMO and E1. Predicted decreases in the elongation zone after salt stress and minimal changes in the meristem after osmotic stress suggest that steady-state changes alone may not fully capture the stress response. Dynamic responses to stress may occur at earlier times, motivating further investigation of the transient dynamics of SUMOylation following stress.

### Experimental data reveal rapid dynamic changes in key SUMO-cycle components in response to abiotic stress

The steady-state model, given the SUMO Cell Atlas data (14), predicts reduced SUMOylation following salt stress in the elongation zone and minimal changes in the meristem following osmotic stress. These predictions contrast with previous evidence that stress promotes SUMOylation (5, 16, 18, 19). This apparent discrepancy could partly reflect differences in spatial resolution, as previous measurements of the SUMOylome were made at the whole-root or whole-plant level, whereas our model predicts tissue-specific responses. We also hypothesized that the single measurement at 3 h after stress application did not capture the transient dynamics of the SUMOylation response. Given that SUMOylation is known to be rapidly induced by stress, an early increase may no longer be evident at 3 h. To test this hypothesis, we quantified the temporal dynamics of key SUMO-cycle components following salt and osmotic stress over time courses of 3 h and 5 h, respectively (Fig. 3).

**Fig 3.**
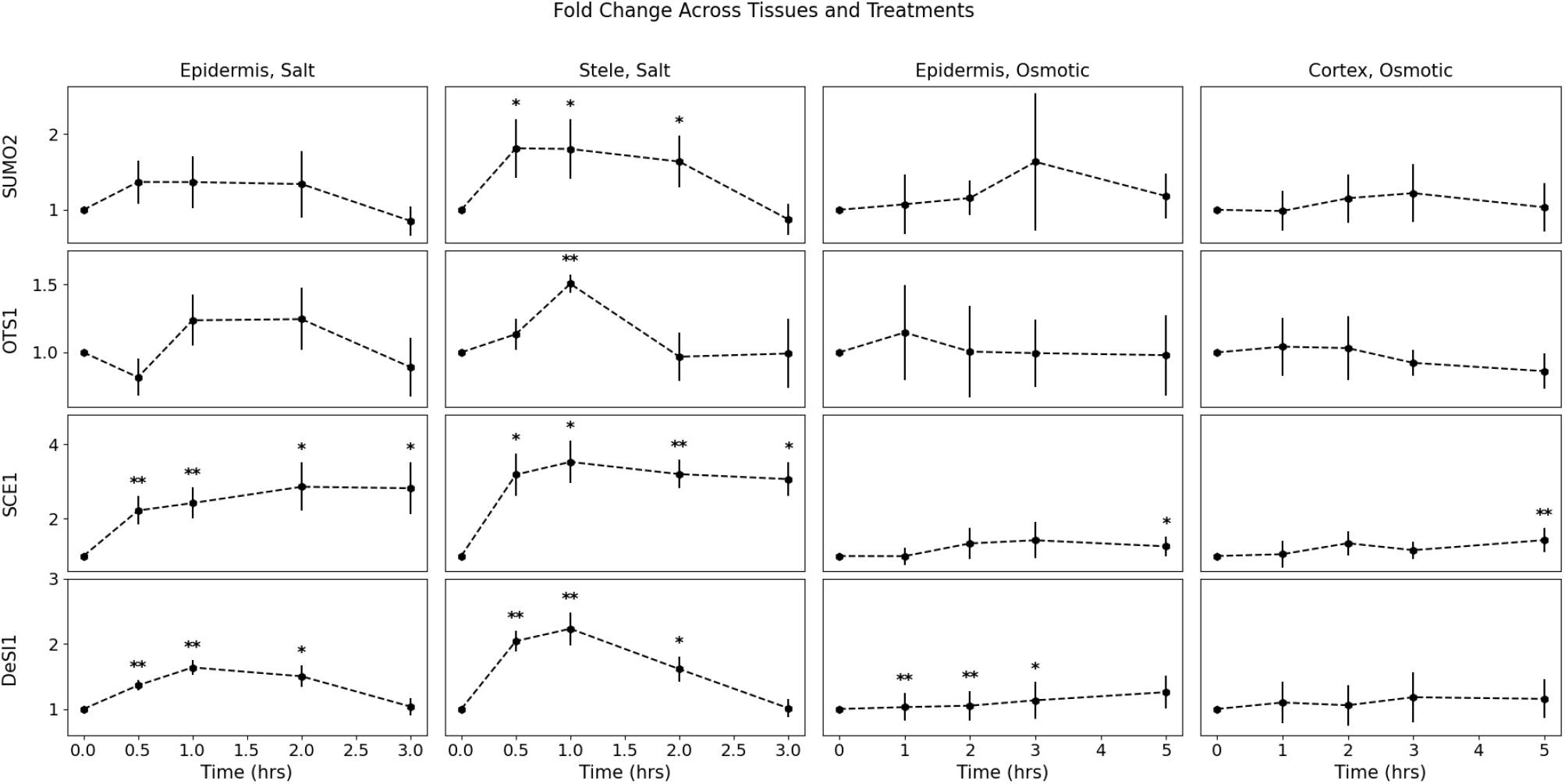
Experimental measurements of the dynamic changes in SUMO-cycle components after abiotic stress. Fold changes in the abundance of SUMO, ULP, E2, and DeSI isoforms (SUMO2, OTS1, SCE1, and DeSI1, respectively) in specified root tissues following stress. Columns show: (1) elongation epidermis under salt stress, (2) elongation stele under salt stress, (3) elongation epidermis under osmotic stress, and (4) elongation cortex under osmotic stress. Fold anges for salt stress were computed relative to the 0 h time point, whereas fold changes for osmotic stress were computed relative to the corresponding untreated baseline experiment. Points show mean fold change, and the error bars show standard deviations (see SI Appendix, Section 4.1). * indicate significant changes (P<0.05), while ** indicate highly significant changes (P<0.01). For the salt treatment experiment, n=3, except for SUMO2 at 2 h, for which n=4. For the mannitol-treatment experiment, n = 6 for SUMO2, except for the 5 h control, for which n = 5; n = 4 for OTS1; and n = 6 for SCE1 and DeSI1. The dynamic model uses the linearly interpolated fold-change trajectories shown here as time-dependent inputs (see SI Appendix, Section 4).

For detailed analysis, we focused on representative proteins from four key protein families: SUMO modifiers, E2 conjugation enzymes, ULP-type and DeSI-type proteases. These proteins are central regulators of SUMOylation and display substantial responses to both salt and osmotic stress. SUMO2 was selected as the primary SUMO modifier because of its ubiquitous expression across root tissues, whereas SCE1, the E2 enzyme, was selected because it is the only SUMO-cycle component significantly upregulated under all stress conditions in the SUMO Cell Atlas (14). OTS1 and DeSI1 were selected as representatives of the ULP and DeSI families, respectively, because both are highly abundant. Additionally, OTS1 has also been shown to have a critical role in salt-stress response (16).

We quantified changes in the four components in the elongation zone in two tissues for each stress condition: the epidermis and stele for salt stress, and the epidermis and cortex for osmotic stress. The epidermis was sampled under both stress conditions to enable direct comparison of the early responses to the two stresses: the steady-state model predicted that the 3 h epidermal measurements would lead to a reduction in SUMOylated target protein after salt stress and an increase in SUMOylated target protein after osmotic stress (Fig. 2D, E). The stele was selected for analysis of the salt-stress response as many SUMO-cycle components exhibited a strong salt-induced regulation in inner root tissues in the SUMO Cell Atlas (14) (Fig. 2B), whereas the cortex was chosen to observe osmotic-stress responses based on its established role in hydrotropic responses (28).

The time-course measurements (Fig. 3) reveal striking differences between salt and osmotic responses. Salt stress induces rapid and pronounced changes in all four imaged components, with SUMO2, DeSI1, and OTS1 all exhibiting strong induction within 1 h, followed by a return to baseline by 3 h, consistent with previously reported salt-induced degradation dynamics of ULP proteases (16). In contrast, SCE1, the sole E2 enzyme, exhibits a significant and sustained increase, highlighting its role as a major driver of rapid stress-induced SUMOylation (14). Notably, this increase does not influence the steady-state model, as E2 does not change the steady-state fold change in SUMOylated target protein (Fig. 1B). This distinction highlights that in these measurements, the components contributing to steady-state behavior return to baseline, whereas components that do not influence steady-state outcomes, such as E2, can remain persistently elevated.

In contrast, osmotic stress produced a more muted response (Fig. 3). Across the epidermis and cortex, changes in SUMO2, OTS1, SCE1, and DeSI1 were generally smaller in magnitude than those observed under salt stress. In summary, these observations suggest that salt stress elicits coordinated and highly dynamic regulation of the SUMO conjugation and deconjugation machinery while the changes after osmotic stress are more muted. Nevertheless, the observed changes in SCE1 and DeSI1 suggest that osmotic stress can still modulate specific components of the SUMO cycle.

### E2 regulates the dynamics of SUMOylated target protein in response to stress

The experimental data revealed dynamic changes in each of the four selected components. Before integrating these data into the model, we sought to isolate the individual contributions of each component by analyzing the predicted response to prescribed step changes in their levels.

Step changes in preSUMO, ULP, E2, and DeSI were imposed through the parameters *Cp, Cu, C*2, and *CD*, representing the ratio between the production rate after stress and the corresponding rate before stress. Increasing each parameter by 50%, alongside perturbations of relevant production parameters, corresponds to a 50% increase in the fold change of the corresponding component. This choice reflects the initial increase observed in all four components in the time-course data (Fig. 3; see SI Appendix, Section 3). We applied a step change to preSUMO rather than SUMO, as SUMO does not have a production parameter.

To better compare the step-change simulations with the later simulations using the dynamic data, we redimensionalized time by introducing the parameter *βs*. This parameter defines the intrinsic timescale of the system: increasing *βs* rescales all the reaction rates equally, so that all processes happen more rapidly. We varied this parameter to examine how timescale influences the transient dynamics. We considered *βs*=1, 2, 3, corresponding to SUMO degradation rates of approximately 1 hour, 30 minutes, and 20 minutes, respectively.

Simulation of step changes required us to specify the 19 remaining unknown parameter values. To define a baseline parameter set, all parameters except *Rs,T, βs*, and those governing the specified step change, were set to unity, corresponding to the assumption that all processes within the SUMO cycle operate on similar timescales. We set *Rs,T* =2.5 to ensure the existence of a finite unstressed steady state, corresponding to a 2.5-fold excess of SUMO relative to target protein under unstressed conditions. Varying the model parameters individually from this baseline produced no qualitative changes in the predicted dynamics (SI Appendix, Section 3, Figs. S7, S8, S12, S14, S18).

The model predicts that a step change in preSUMO produces a proportional increase in the steady-state level of SUMOylated target protein, consistent with the steady-state analysis (Fig. 1B, C). Simulations also reveal transient overshoots before the SUMOylated target-protein level reaches steady state (Fig. 4A), with slower systems exhibiting larger and more prolonged transient overshoots. In contrast, a step change in ULP generates a response driven by competing effects (Fig. 4B): SUMOylated target-protein levels initially decrease due to enhanced deSUMOylation before recovering as increased preSUMO maturation elevates the SUMO pool. A step change in DeSI causes a monotonic reduction in SUMOylated target-protein levels (Fig. 4C) whereas a step change in E2 enhances SUMOylation only transiently, consistent with E2 not affecting the steady state of SUMOylated target-protein level (Fig. 4D). Across all step changes, faster intrinsic timescales lead to more rapid convergence to steady state, whereas slower systems exhibit larger and more prolonged transient peaks and troughs.

**Fig 4.**
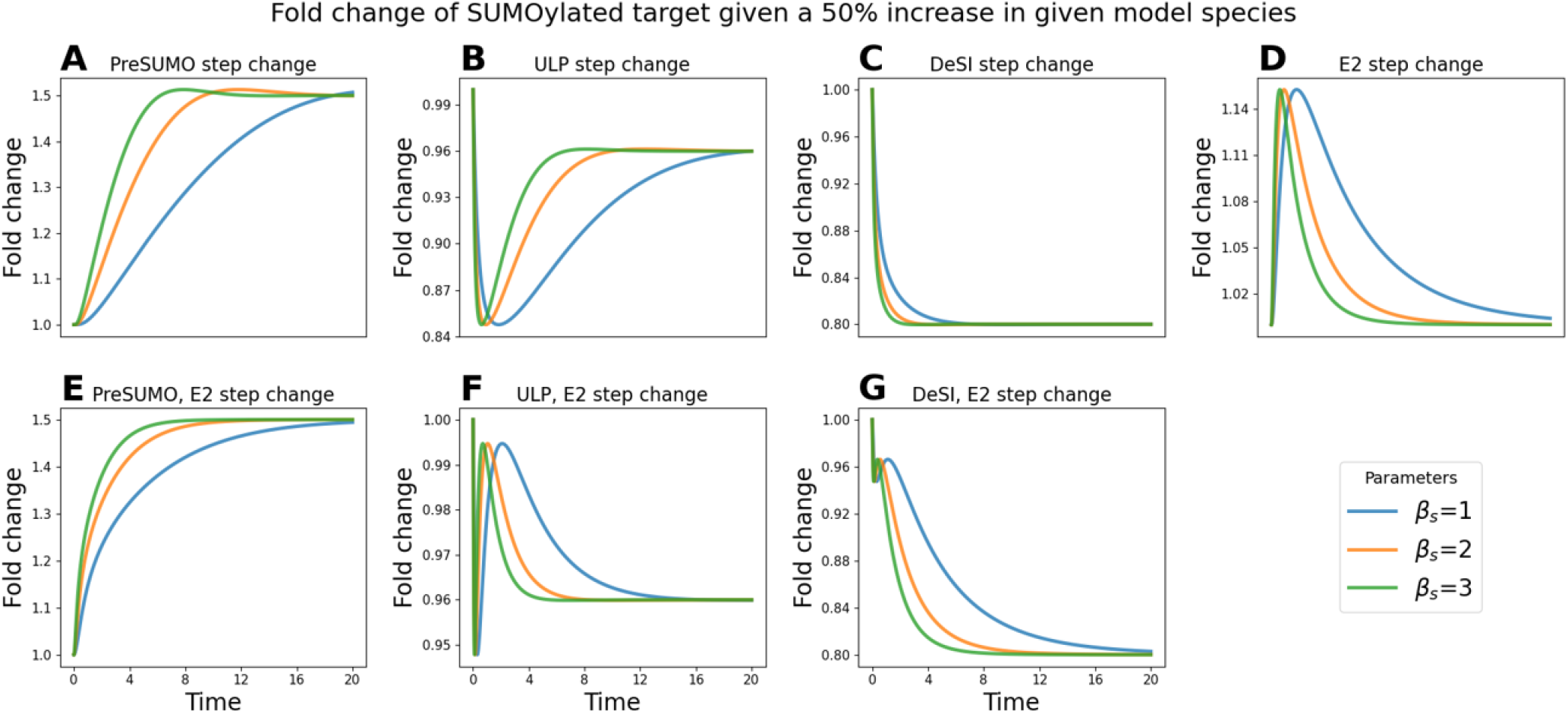
Predicted dynamics of the fold change in SUMOylated target protein following step changes in model components. Time courses of the predicted fold change in SUMOylated target following a 50% step change in preSUMO (A), ULP (B), DeSI (C), E2 (D), preSUMO and E2 simultaneously (E), ULP and E2 simultaneously (F), DeSI and E2 simultaneously (G). Simulations are shown for three values of *β*^*s*^, which sets the intrinsic timescale of the model, with larger values corresponding to a faster SUMO cycle.

As E2 is thought to be a major driver of SUMOylation (14), we also analyzed E2 step changes in combination with the other prescribed perturbations. A simultaneous step change of preSUMO and E2 suppresses the transient overshoot observed following a preSUMO increase (Fig. 4A, E). Combining ULP and E2 step changes reduces both the magnitude and duration of the transient decrease caused by increased ULP (Fig. 4F). Simultaneous increases in E2 and DeSI produce a distinctly non-monotonic response and prolong the time required to reach steady state compared with a DeSI increase alone (Fig. 4G), reflecting the competing effects of enhanced SUMOylation and enhanced deSUMOylation. Together, these results suggest that E2 primarily regulates the speed of SUMOylation and mitigates transient reductions in SUMOylated target-protein levels.

Overall, these simulations demonstrate how the dynamics of SUMOylated target levels are governed by the balance between SUMOylation, driven by levels of SUMO and E2, and deSUMOylation, driven by ULP and DeSI. After onset of stress, deSUMOylation dominates the response by directly affecting SUMOylated target levels. Over longer timescales, however, increased SUMO abundance drives the system towards an elevated steady state, as increases in SUMO must first propagate through the cycle before affecting the level of SUMOylated target protein. Increased levels of E2 accelerate the rise in SUMOylation and reduce the magnitude and duration of transient deSUMOylation-driven decreases in SUMOylation.

### Model predicts tissue- and stress-specific SUMOylation dynamics in response to abiotic stress

We used the time-series data (Fig. 3) as inputs to the dynamic model to investigate how time-dependent changes in SUMO-cycle components propagate to shape the temporal dynamics of the level of SUMOylated target protein. This extends the steady-state analysis (Fig. 2D, E) by capturing transient responses that cannot be represented by a single fold change. To incorporate the experimentally observed dynamics of SUMO-cycle components, we replaced the ODEs governing SUMO, ULP, E2, and DeSI with the linearly interpolated fold-change trajectories shown in Fig. 3. All remaining components evolve according to their original ODE-defined dynamics (SI Appendix, Section 4).

As in the previous analysis, all parameters were set to unity except *Rs,T* which was fixed at 2.5, and *βs*. We considered *βs* = 1 and *βs* = 3 corresponding to intrinsic timescales of approximately 1 h and 20 min, respectively. This allowed us to examine how the intrinsic timescale influences the predicted dynamics. In contrast to the step-change analysis, mature SUMO levels are prescribed directly from the experimental data rather than through preSUMO production. Consequently, ULP acts solely through deSUMOylation in these simulations.

To account for experimental uncertainty, the fold change at each time point was represented by a normal distribution with a mean and standard deviation obtained from the corresponding experimental data point. We drew 1,000 samples from these distributions; non-positive samples were resampled until a positive value was obtained. The resulting samples were propagated through the dynamic model (Fig. 5A–H). This enabled us to quantify the uncertainty in the predicted SUMOylated target dynamics arising from variability in the experimental measurements.

**Fig 5.**
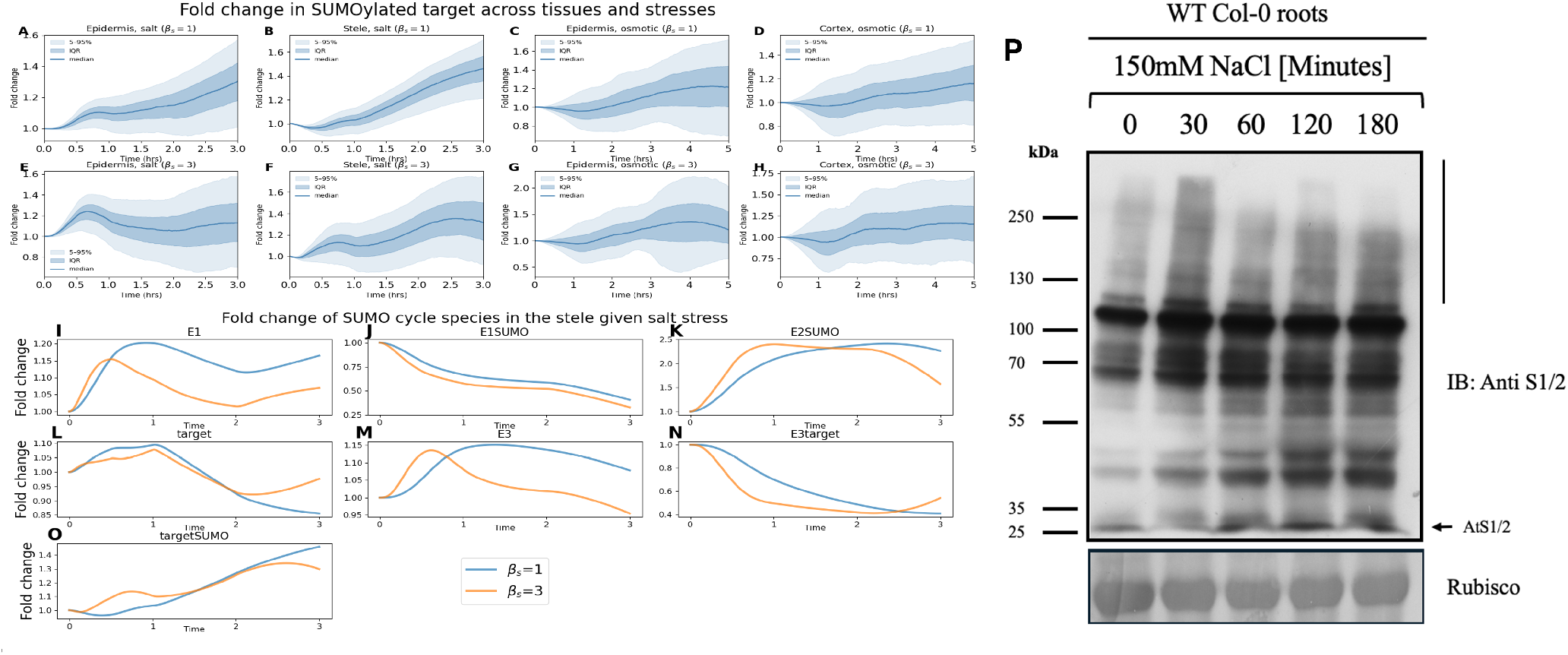
Predicted effect of observed dynamic changes on downstream SUMO-cycle components. (**A–H**)Predicted fold change in SUMOylated target protein across tissues, stresses, and intrinsic timescales using the experimental data in Fig. 3 as model inputs. At each time point, the measured fold change was integrated by sampling from a normal distribution with mean and standard deviation given by the experimental data. One thousand samples were drawn from these distributions, with non-positive values resampled until a positive value was obtained. The resulting samples were then propagated through the model. Solid lines show the median prediction, dark shading the interquartile range, and light shading the 5-95% prediction interval. Simulations are shown for two values of *β*^*s*^(*β*^*s*^ = 1 and *β*^*s*^ = 3), which sets the intrinsic timescale of the SUMO cycle, with larger values corresponding to faster dynamics. **I–O**) Predicted fold changes in E1 (I), E1 *·* SUMO (J), E2 *·* SUMO (K), target (L), E3 (M), E3 *·* target, and target *·* SUMO (O) for the stele under salt stress, for *β*_*s*_ = 1 and *β*_*s*_ = 3 (**P**) Western blot analysis of total SUMOylated protein derived from 5-day-old wild-type Col-0 *Arabidopsis thaliana* roots treated with 150mM NaCl. The vertical bar indicates the laddering effect produced during the increased accumulation of SUMO1/2 conjugates and the arrowhead indicates nonconjugated free SUMO1/2. The bottom panel displays Ponceau staining of ribulose-1,5-bis-phosphate carboxylase/oxygenase (Rubisco) small subunit (RbscS) to serve as a loading control. This experiment was repeated 3 times.

In the epidermis following salt stress, the predicted SUMOylated target response depends strongly on the intrinsic timescale. For a slow timescale (corresponding to *βs* = 1), trajectories within the interquartile range indicate a predominantly monotonic increase in SUMOylated target-protein, with only a subtle transient rise at approximately 0.7 h before continuing upward, with levels increasing beyond 2 h across the 5-95% trajectory range (Fig. 5A). For a faster timescale (*βs* = 3), across trajectories within the interquartile range the inflection at 0.7 h sharpens into a pronounced early peak followed by a decline between 1 h and 3 h (Fig. 5E). The predicted peak–decline structure is driven by the dynamic ULP response (Fig. 3): an initial decrease in ULP promotes SUMOylation, whereas the subsequent increase enhances deSUMOylation, with variability in the sampled data driving variability in the magnitude of this response. Consistent with the step change analysis (Fig. 4), the dynamics depend strongly on the intrinsic timescale, determining whether processes occur as temporally separated events or blend into a single response: slower dynamics sustain the rise in target by delaying SUMO propagation through the cycle, whereas faster dynamics amplify both the peak and decline through faster propagation and stronger E2-mediated acceleration.

The predicted response in the stele following salt stress is qualitatively similar to that in the epidermis for the slower timescale (Fig. 5B). However, as ULP and DeSI peak earlier in the stele, at approximately 1 h (Fig. 3), the model predicts a brief early dip in SUMOylated target before the sustained increase; this initial effect is consistent across the range of sampled inputs for the first ~0.25 h after stress application. For faster timescales, where processes occur as distinct, sequential events, the model predicts a double peak across the interquartile range of predicted responses, with peaks near 0.5 h and 2.5 h (Fig. 5F); the intermediate decrease coincides with transient increases in ULP and DeSI, while the peaks emerge as elevated SUMO and E2 drive renewed SUMOylation through the faster dynamics.

Following osmotic stress, the model predicts an initial decrease in median SUMOylated target levels in the epidermis before SUMOylation subsequently increases (Fig. 5C). As in the step-change simulations (Fig. 4), the early decrease reflects enhanced deSUMOylation via increased ULP, while the later increase reflects elevated SUMO and E2 levels; predicted responses nonetheless vary considerably across the full sampled range, reflecting the large standard deviation in the SUMO2 reporter data at 3 h (Fig. 3). Faster timescales shift the median peak earlier, to around 3.5 h (Fig. 5G), consistent with E2 accelerating SUMOylation propagation; the subsequent decline coincides with increased DeSI and reduced SUMO.

A similar median response is predicted in the cortex for slower timescales, where an initial reduction in SUMOylated target reflects elevated ULP and DeSI activity alongside decreased SUMO abundance, though predictions vary less than in the epidermis across the sampled data (Fig. 5D). For faster timescales, the model predicts a wider range of responses and a higher median peak; within this wider range of predicted behaviors, oscillations emerge (Fig. 5H) whose peaks coincide with peaks in SUMO and E2 and whose troughs coincide with peaks in ULP and DeSI (Fig. 3).

To gain a better understanding of how the dynamics of SUMOylation arise, we examined the predicted dynamics of remaining SUMO-cycle components considering the response to salt in the stele (Fig. 5I–O). Taking the mean fold changes as inputs (Fig. 3), the model predicts that a rapid increase in E2 (Fig. 3) drives formation of E2 · SUMO (Fig. 5K), which in turn depletes E1 · SUMO(Fig. 5J). Consequently, E1 · SUMO decreases across all times, as elevated E2 shifts flux toward E2-SUMO formation, rendering E1 · SUMO a limiting intermediate. Release of E1 from the declining E1 · SUMO pool produces a transient increase in free E1, although this is tempered by its consumption in E1 · SUMO ; faster systems sustain a higher transient peak (Fig. 5I).

The dynamics of E2· SUMO are determined by the interplay between SUMO and E2 availability. In slower systems, the level of E2 SUMOclosely follows the dynamics of E2 (Fig. 3), whereas in faster systems it more closely tracks SUMO availability (Fig. 3, 5K). Free target protein initially increases due to deSUMOylation of SUMOylated target but later decreases as rising E2 · SUMO drives renewed SUMOylation and depletion of free target protein (Fig. 5L). In faster systems, target levels begin to recover at approximately 2 hours, consistent with increased DeSI and ULP activity. Correspondingly, E3 initially increases as E3 · target is consumed in SUMOylated target formation but subsequently declines as elevated ULP and DeSI levels promote deSUMOylation and shift the system back towards E3 · target formation by increasing free target protein availability (Fig. 5M, N).

Taken together, these simulations predict distinct SUMOylation dynamics across tissues and stress conditions, with qualitatively different behaviors emerging as the intrinsic timescale increases: faster timescales produce distinct, sequential events and greater variability in the predicted response, while slower timescales allow the regulatory processes to overlap into a more gradual, combined response. Consistent with the step-change analysis, the predicted responses are governed by the dynamics of SUMOylation and deSUMOylation, with deSUMOylation dominating shortly after stress and transient SUMOylation peaks emerging in faster systems. Across all tissues examined, E2 emerges as a key regulator of the timing and magnitude of the transient SUMOylation responses, with its dynamics determining how rapidly SUMOylated-target-protein levels change and the magnitude of the resulting peaks.

### SUMO conjugates exhibit a twin-peaked response to salt stress, consistent with the model predictions

To test whether the dynamic response predicted by the model could be detected experimentally, we measured SUMO1/2 conjugate abundance in salt-treated *Arabidopsis thaliana* roots. Western blot analysis revealed an increase in SUMO conjugates within 30 minutes of salt exposure. The conjugate levels decreased at 60 minutes and increased again at 120 to 180 minutes (Fig. 5P).

These whole-root measurements displayed SUMOylation dynamics that are consistent with the predicted twin-peaked dynamics in SUMOylated target protein levels in the stele after salt stress (Fig.5F). Importantly, the intrinsic timescale of the system is predicted to strongly shape this response, affecting both peak timing and amplitude: faster systems produce sharper and earlier peaks, whereas slower systems exhibit broader and delayed responses (Fig.5B, F).

Although the precise dynamics remain parameter dependent, these simulations suggest that key processes in the SUMO cycle could operate on comparable intrinsic timescales. The observed dynamics in the Western blot data are more consistent with the simulations with *βs* = 3, corresponding to a SUMO-cycle timescale of approximately 20 minutes.

We conclude that these experimental data support the prediction that SUMOylation is a highly dynamic process, enabled through temporal and spatial regulation of multiple enzymes and ligases within the SUMO cycle.

## Discussion

PTMs such as SUMOylation provide cells with a rapid and reversible means of altering protein function in response to environmental change, without requiring de novo transcription or translation. SUMOylation is a cyclical process, involving multiple families of enzymes, ligases and proteases. This complexity, combined with a lack of knowledge of the associated kinetic parameters, makes intuition alone insufficient to determine how changes in individual SUMO-cycle components affect the proportion of SUMOylated target protein. Here, we used mathematical modeling to analyze published tissue-specific reporter data that quantified changes in SUMO-cycle components in response to osmotic and salt stress (14). The model revealed an important distinction between steady-state and dynamic control of the SUMO cycle. Under the assumptions of the model, free SUMO, E1, ULP and DeSI determine the steady-state level of SUMOylated target protein, whereas the E2 conjugating enzyme substantially alters the speed and transient magnitude of the response without changing its steady-state level.

Osmotic and salt stress can induce rapid accumulation of SUMO conjugates in plants within minutes of stress onset, allowing the cell proteome to reset and fine-tune its stress-specific response (16, 29–32). The increased SUMOylation response during salt stress has been shown to be partially driven through rapid degradation of OTS1 inhibiting deSUMOylation of targets (16, 33). However, incorporating the 3-hour SUMO Cell Atlas data into the steady-state model did not predict an increase in the level of SUMOylated target protein across all root tissues. Although the model predicted increased SUMOylated-target-protein levels in the meristem after salt stress and in the elongation zone after osmotic stress, the model also predicted no substantial change in the levels in the meristem after osmotic stress and surprisingly predicted a decrease in the elongation zone after salt stress. The discrepancy could reflect a localized versus systemic SUMOylation response, as most SUMO blots are performed on whole plant tissue, while the changes predicted here are in specific tissues of the growth zones near the root tip (a section approximately 5mm in length), each of which have distinct abundances of SUMO-cycle components. These zones have varying functions in root development, which could underlie changes in SUMOylation (34). Furthermore, the root meristem is covered by the lateral root cap, which protects the meristematic cells from external stresses and could explain limited changes in SUMOylation in response to osmotic stress (35–37). However, we hypothesized that a key limitation in this steady-state analysis is that the data, collected 3 hours after stress onset, did not capture the early dynamics of SUMO-cycle components (Fig 5P) (5, 16, 18, 19).

Motivated by this hypothesis, we generated imaging time courses for key SUMO-cycle components during salt and osmotic stress. Using these data, the model predicted that SUMOylation is highly dynamic with variations between stresses and tissues that could not be inferred from the previous 3 h timepoint data (14). The model predicted how SUMOylation dynamics arise from the balance between SUMO conjugation, driven by free SUMO and E2, and deSUMOylation, mediated by DeSI and ULP. The predictions suggest that since changes in the proteases act directly on SUMOylated target protein, they lead to rapid changes at earlier time points, whereas changes in free SUMO must propagate through the cycle (via the E1, E2 and E3 enzymes) before affecting the SUMOylated-target-protein levels. Furthermore, the model suggests that E2 accelerates this propagation, driving the system to steady-state when paired with other changes, and strongly affects the transient dynamics despite not influencing the steady-state level of SUMOylated target protein. The model showed how the intrinsic timescale of the SUMO cycle determines whether the effects of successive component changes overlap to produce a gradual SUMOylation response or are temporally separated and mediate distinct peaks and troughs. These model predictions demonstrate why single-timepoint measurements are unlikely to explain the behavior of a dynamic and cyclical PTM system.

Under salt stress, the model predicted twin-peaked SUMOylation dynamics in the stele in response to salt, particularly under the faster-dynamics parameterization, due to the misalignment of peaks in the levels of the conjugating and deconjugating enzymes. Consistent with this model prediction, subsequent Western blot analysis of whole roots detected two periods of increased SUMO-conjugate accumulation separated by an intervening trough. We hypothesize that this double peak may arise because salt induces both an ionic response and an osmotic response, which potentially occur on different time scales. Whilst these data provide support for the model predictions, these are different outputs: the model predicts the abundance of a generic SUMOylated target in a specified tissue, whereas the Western blot reports aggregate SUMO1/2-conjugate abundance across many endogenous targets and the whole root. Developing molecular techniques to quantify SUMOylation in individual tissues would enable better validation of the model predictions and greatly enhance our understanding of SUMOylation responses.

While the Western blot data support our model predictions, these are based on time-courses of a limited number of representative SUMO-cycle components. It is likely that other isoforms in the ULP and DeSI families may exhibit different dynamic responses (14); incorporating other members of these families could therefore modify the predicted SUMOylation dynamics and provide a more complete picture of tissue-specific regulation. Furthermore, the model necessarily includes several simplifying assumptions: it represents a generic SUMO target, excludes polySUMOylation, and assumes enzymatic reactions operate below saturation. Additionally, the free mature SUMO is prescribed from the measured SUMO2 reporter abundance and so the role of ULP in SUMO maturation is not explicitly simulated.

Our model builds upon previous mathematical modeling of SUMOylation (20–22) by providing, to our knowledge, the first mechanistic model of the SUMO cycle in plants, incorporating the plant-specific DeSI protease and explicitly representing E3-mediated target SUMOylation. The inclusion of DeSI has important consequences for both steady-state and dynamic behavior: differences between the fold changes of DeSI and ULP increase the range of predicted steady-state SUMOylation responses across deSUMOylation rates, while dynamic changes in DeSI are a major driver of changes in SUMOylated target levels. Furthermore, our model directly incorporates experimentally measured changes in SUMO-cycle components and uses these data to predict the response of the SUMO cycle to abiotic stress. This provides a framework for linking experimentally observed regulation of individual SUMO components to emergent changes in SUMOylation.

The model also generates experimentally testable predictions. The dynamic data (Fig. 3) show a sustained increase in SCE1 abundance after salt treatment. These elevated E2 levels may result in the SUMOylation machinery responding differently to a subsequent stress, as there may be less scope for a further stress-induced increase in E2 abundance. This observation suggests a potential mechanism for the previously reported refractory period (24), during which the SUMOylation response to a subsequent stress is attenuated. The timing of the second stress may therefore be critical, as the secondary response could depend on the extent to which SCE1 has returned towards its unstressed level. This could be investigated by applying a second stress at different intervals after the initial salt treatment and determining how sustained SCE1 abundance affects the magnitude and timing of the subsequent SUMOylation response. Additionally, partial loss-of-function lines targeting individual SUMO components, such as *sumo2, sce1* or *ots1*, could test the predicted contribution of individual components to the magnitude and dynamics of the stress-induced SUMOylation response.

Our conclusions focus on the qualitative features of the predicted dynamics and we acknowledge that the precise timing and amplitude of the predicted peaks should be interpreted cautiously. Nevertheless, the predictions support the broader conclusion that PTM systems are highly dynamic, precisely regulated systems that achieve tissue- and stress-specific control through the coordinated interplay of multiple SUMO-cycle components. The model predicts that perturbing E2 primarily alters the timing and transient magnitude of SUMOylation, whereas perturbing SUMO, E1, ULP or DeSI alter the balance between SUMOylation and deSUMOylation, and thus control both the dynamic and steady-state behaviors.

## Materials and Methods

### Mathematical modeling

Full details of the derivation of the model equations are provided in the SI text sections 1-4. Simulations were performed using Python. All code and data used in this paper will be deposited in GitLab (https://gitlab.com/leahband/SUMO_pathway_model/).

### Plant material and growth conditions

Reporter lines from the SUMO Cell Atlas (14) were used in this study. Seeds were surface sterilized for 4 min in 30% (v/v) bleach containing 0.005% Triton X-100, followed by five washes with sterile water and were then stratified at 4 °C for 48 h in the dark. Seeds were germinated on media containing half-strength Murashige and Skoog (MS) media (Sigma), 0.4% sucrose and 1% Bacto-agar at pH 5.7. Seedlings were grown vertically for 6 days under day temperature of 23 °C and night temperature of 18 °C with a 16-h photoperiod (150 *µ*mol *m*^*−*^2*s*^*−*^1).

### Confocal imaging and treatments

6-day-old plants were treated with 150mM NaCl (salt stress) or 300mM mannitol (osmotic stress) as in (14) and imaged at multiple time points using a Leica SP8 confocal microscope (Leica Microsystems) using a 20x objective. The same confocal settings and tissue-specific data analysis were used as previously described in (14).

### Western blot

5-day-old seedlings were transferred to a set-up containing either ½ MS liquid media or ^1^ MS liquid media supplemented with 150mM NaCl. The seedling roots were exposed for the required time points – 0 minutes, 30 minutes, 60 minutes, 120 minutes and 180 minutes. After the required treatment time, 50 roots were removed using a scalpel blade and flash frozen in liquid nitrogen. Total protein was extracted using a SUMO extraction buffer containing 25mM Tris-HCL (pH 8), 150mM NaCl, 1mM EDTA, 2% glycerol, 0.1% Triton x-100, 0.7% NP-40, 70mM NEM, 50mM KCL, 20mM MgCl2, 0.25% sodium deoxycholate and a protease inhibitor cocktail tablet at a working concentration of 1 tablet per 10ml buffer. After protein extraction, the samples were loaded onto an 8% SDS-polyacrylamide gel and detected following the protocol outlined in Banda et al., 2025. The membranes were probed with anti-SUMO1/2 (anti AtS1/2) antibody, followed by anti-Rabbit HRP (Abcam) antibody.

### Use of AI

Microsoft 365 Copilot and GPT-4o (ChatGPT) were used during the final preparation of the manuscript for proofreading and language editing. Claude 5 Sonnet and Codex were used for assistance with debugging and refining the Python code. All AI-assisted code was reviewed and tested by the authors. The authors take full responsibility for the final manuscript, code and results.

## ACKNOWLEDGMENTS

We thank UKRI BBSRC for the strategic longer larger Grant, SUMO code (BB/v003534/1), to A.S., M.J.B and L.R.B.

