## Supplementary Information for "Mathematical modeling of SUMOylation reveals tissue-specific and dynamic responses to abiotic stress"

**Drynda et al.**

#### This PDF file includes:

Supporting text
Figs. S1 to S29
Tables S1 to S3
SI References

### Supporting Information Text

In this Supplementary Information, we provide further details on the mathematical modeling. The aim of the model is to predict the level of SUMOylated target in response to stress, first using the SUMO Cell Atlas data (1) and then the newly generated dynamic data. In section 1, we derive a nondimensional model of the SUMO cycle that describes the dynamics of SUMO, the functional E1 complex, the enzymatic families, target protein and their complexes. In section 2, we use the model to derive a function that predicts the fold change of SUMOylated target given the measured fold changes of SUMO, ULP, E1, and DeSI. Furthermore, we adapt the SUMO Cell Atlas data (1) so it can be used in the model. In section 3, we move on from steady state analysis and introduce step changes into the model to analyze how the dynamics of the model respond to them. Finally, in section 4, we process the dynamic data, construct the dynamic model and describe how the dynamic data is input into it.

### 1. Base SUMOylation Model

In this section, we derive the chemical reactions underlying the SUMO cycle and use them to formulate a system of ODEs describing the time evolution of the cycle species concentrations (section 1.1). We then solve this system for steady state (section 1.2) and use these solutions along with stress-induced production changes to nondimensionalize the model (section 1.3).

**1.1. Dimensional model ODEs.** The SUMO cycle in *Arabidopsis* (Fig. 1A) can be represented by the following set of chemical reactions, defined in terms of the SUMO isoform pool, the functional E1 complex, the E2, E3, DeSI, and ULP enzyme families, the target protein, and the resulting complexes:

$$\text{preSUMO} + \text{ULP} \xrightarrow{k_1} \text{SUMO} + \text{ULP} \tag{1a}$$

$$\text{E1} + \text{SUMO} \xrightarrow{k_2} \text{E1} \cdot \text{SUMO} \tag{1b}$$

$$\text{E2} + \text{E1} \cdot \text{SUMO} \xrightarrow{k_3} \text{E2} \cdot \text{SUMO} + \text{E1} \tag{1c}$$

$$\text{target} + \text{E2} \cdot \text{SUMO} \xrightarrow{k_4} \text{target} \cdot \text{SUMO} + \text{E2} \tag{1d}$$

$$\text{E3} + \text{target} \xrightarrow{k_5} \text{E3} \cdot \text{target} \tag{1e}$$

$$\text{E3} \cdot \text{target} + \text{E2} \cdot \text{SUMO} \xrightarrow{k_6} \text{target} \cdot \text{SUMO} + \text{E2} + \text{E3} \tag{1f}$$

$$\text{target} \cdot \text{SUMO} + \text{ULP} \xrightarrow{k_7} \text{SUMO} + \text{target} + \text{ULP} \tag{1g}$$

$$\text{target} \cdot \text{SUMO} + \text{DeSI} \xrightarrow{k_8} \text{SUMO} + \text{target} + \text{DeSI} \tag{1h}$$

$$\emptyset \xrightarrow{\alpha_p} \text{preSUMO} \xrightarrow{\beta_p} \emptyset \tag{1i}$$

$$\emptyset \xrightarrow{\alpha_u} \text{ULP} \xrightarrow{\beta_u} \emptyset \tag{1j}$$

$$\emptyset \xrightarrow{\alpha_1} \text{E1} \xrightarrow{\beta_1} \emptyset \tag{1k}$$

$$\emptyset \xrightarrow{\alpha_2} \text{E2} \xrightarrow{\beta_2} \emptyset \tag{1l}$$

$$\emptyset \xrightarrow{\alpha_3} \text{E3} \xrightarrow{\beta_3} \emptyset \tag{1m}$$

$$\emptyset \xrightarrow{\alpha_T} \text{target} \xrightarrow{\beta_T} \emptyset \tag{1n}$$

$$\emptyset \xrightarrow{\alpha_D} \text{DeSI} \xrightarrow{\beta_D} \emptyset \tag{1o}$$

$$\text{SUMO} \xrightarrow{\beta_s} \emptyset \tag{1p}$$

Here, $\alpha_i$ and $\beta_i$ ($i \in \{p, u, 1, 2, 3, T, D\}$) denote production and degradation rates of the corresponding species. The free SUMO degradation rate $\beta_s$ represents SUMO turnover and is required to balance SUMO maturation and obtain a finite steady state. The parameters $k_j$ ($j \in \{1, 2, ..., 8\}$) denote the binding rates of the different steps in the SUMO cycle. We note that ULP and DeSI are treated as a catalytic species whose concentrations are regulated only by production and degradation.

Let

$$X = \{\text{preSUMO}, \text{ULP}, \text{SUMO}, \text{E1}, \text{E1}\cdot\text{SUMO}, \text{E2}, \text{E2}\cdot\text{SUMO}, \text{target}, \text{E3}, \text{E3}\cdot\text{target}, \text{target}\cdot\text{SUMO}, \text{DeSI}\}$$

be the set of all the species in the chemical reactions listed above. For each $x \in X$, we let the variable $[x]$ denote its concentration.

Applying the law of mass action to Eq. (1) yields the following system of ordinary differential equations (ODEs):

$$\frac{\mathrm{d}}{\mathrm{d}t}[\text{preSUMO}] = \alpha_p - \beta_p[\text{preSUMO}] - k_1[\text{preSUMO}][\text{ULP}], \quad [2a]$$

$$\frac{\mathrm{d}}{\mathrm{d}t}[\text{ULP}] = \alpha_u - \beta_u[\text{ULP}], \quad [2b]$$

$$\frac{\mathrm{d}}{\mathrm{d}t}[\text{SUMO}] = k_1[\text{preSUMO}][\text{ULP}] - k_2[\text{E1}][\text{SUMO}] + k_7[\text{target}\cdot\text{SUMO}][\text{ULP}] + k_8[\text{target}\cdot\text{SUMO}][\text{DeSI}] - \beta_s[\text{SUMO}], \quad [2c]$$

$$\frac{\mathrm{d}}{\mathrm{d}t}[\text{E1}] = \alpha_1 - \beta_1[\text{E1}] - k_2[\text{E1}][\text{SUMO}] + k_3[\text{E2}][\text{E1}\cdot\text{SUMO}], \quad [2d]$$

$$\frac{\mathrm{d}}{\mathrm{d}t}[\text{E1}\cdot\text{SUMO}] = k_2[\text{E1}][\text{SUMO}] - k_3[\text{E2}][\text{E1}\cdot\text{SUMO}], \quad [2e]$$

$$\frac{\mathrm{d}}{\mathrm{d}t}[\text{E2}] = \alpha_2 - \beta_2[\text{E2}] - k_3[\text{E2}][\text{E1}\cdot\text{SUMO}] + k_4[\text{target}][\text{E2}\cdot\text{SUMO}] + k_6[\text{E2}\cdot\text{SUMO}][\text{E3}\cdot\text{target}], \quad [2f]$$

$$\frac{\mathrm{d}}{\mathrm{d}t}[\text{E2}\cdot\text{SUMO}] = k_3[\text{E2}][\text{E1}\cdot\text{SUMO}] - k_4[\text{target}][\text{E2}\cdot\text{SUMO}] - k_6[\text{E2}\cdot\text{SUMO}][\text{E3}\cdot\text{target}], \quad [2g]$$

$$\frac{\mathrm{d}}{\mathrm{d}t}[\text{target}] = \alpha_T - \beta_T[\text{target}] - k_4[\text{target}][\text{E2}\cdot\text{SUMO}] - k_5[\text{E3}][\text{target}] + k_7[\text{target}\cdot\text{SUMO}][\text{ULP}] + k_8[\text{target}\cdot\text{SUMO}][\text{DeSI}], \quad [2h]$$

$$\frac{\mathrm{d}}{\mathrm{d}t}[\text{E3}] = \alpha_3 - \beta_3[\text{E3}] - k_5[\text{E3}][\text{target}] + k_6[\text{E2}\cdot\text{SUMO}][\text{E3}\cdot\text{target}], \quad [2i]$$

$$\frac{\mathrm{d}}{\mathrm{d}t}[\text{E3}\cdot\text{target}] = k_5[\text{E3}][\text{target}] - k_6[\text{E2}\cdot\text{SUMO}][\text{E3}\cdot\text{target}], \quad [2j]$$

$$\frac{\mathrm{d}}{\mathrm{d}t}[\text{target}\cdot\text{SUMO}] = k_4[\text{target}][\text{E2}\cdot\text{SUMO}] + k_6[\text{E2}\cdot\text{SUMO}][\text{E3}\cdot\text{target}] - k_7[\text{target}\cdot\text{SUMO}][\text{ULP}] - k_8[\text{target}\cdot\text{SUMO}][\text{DeSI}], \quad [2k]$$

$$\frac{\mathrm{d}}{\mathrm{d}t}[\text{DeSI}] = \alpha_D - \beta_D[\text{DeSI}]. \quad [2l]$$

**1.2. Dimensional model steady-state solutions.** For all species $x \in X$, we let $[x]_{\text{SS}}$ denote the concentration of $x$ at steady state. Solving Eq. (2) for steady state we obtain

$$[\text{preSUMO}]_{\text{SS}} = \frac{\alpha_p \beta_u}{\alpha_u k_1 + \beta_p \beta_u}, \quad [3a]$$

$$[\text{ULP}]_{\text{SS}} = \frac{\alpha_u}{\beta_u}, \quad [3b]$$

$$[\text{SUMO}]_{\text{SS}} = \frac{\alpha_p \alpha_u k_1}{\beta_s(\alpha_u k_1 + \beta_p \beta_u)}, \quad [3c]$$

$$[\text{E1}]_{\text{SS}} = \frac{\alpha_1}{\beta_1}, \quad [3d]$$

$$[\text{E1} \cdot \text{SUMO}]_{\text{SS}} = \frac{\alpha_p \alpha_u \alpha_1 \beta_2 k_1 k_2}{\alpha_2 \beta_s \beta_1 k_3 (\alpha_u k_1 + \beta_p \beta_u)}, \quad [3e]$$

$$[\text{E2}]_{\text{SS}} = \frac{\alpha_2}{\beta_2}, \quad [3f]$$

$$[\text{E2} \cdot \text{SUMO}]_{\text{SS}} = \frac{\alpha_p \alpha_u \alpha_1 \beta_T \beta_3 k_1 k_2 - \alpha_u \alpha_T \alpha_3 \beta_s \beta_1 k_1 k_5 - \alpha_T \alpha_3 \beta_p \beta_u \beta_s \beta_1 k_5}{\alpha_T \beta_s \beta_1 \beta_3 k_4 (\alpha_u k_1 + \beta_p \beta_u)}, \quad [3g]$$

$$[\text{target}]_{\text{SS}} = \frac{\alpha_T}{\beta_T}, \quad [3h]$$

$$[\text{E3}]_{\text{SS}} = \frac{\alpha_3}{\beta_3}, \quad [3i]$$

$$[\text{E3} \cdot \text{target}]_{\text{SS}} = \frac{\alpha_T^2 \alpha_3 \beta_s \beta_1 k_4 k_5 (\alpha_u k_1 + \beta_p \beta_u)}{\beta_T k_6 (\alpha_p \alpha_u \alpha_1 \beta_3 \beta_T k_1 k_2 - \alpha_u \alpha_T \alpha_3 \beta_s \beta_1 k_1 k_5 - \alpha_T \alpha_3 \beta_p \beta_u \beta_s \beta_1 k_5)}, \quad [3j]$$

$$[\text{target} \cdot \text{SUMO}]_{\text{SS}} = \frac{\alpha_p \alpha_u \alpha_1 \beta_u \beta_D k_1 k_2}{\beta_s \beta_1 (\alpha_u k_1 + \beta_p \beta_u)(\alpha_u \beta_D k_7 + \alpha_D \beta_u k_8)}, \quad [3k]$$

$$[\text{DeSI}]_{\text{SS}} = \frac{\alpha_D}{\beta_D}. \quad [3l]$$

From the expression for $[\text{E3} \cdot \text{target}]_{\text{SS}}$, Eq. (3j), the following inequality must hold for the steady-state concentrations to be both positive and finite:

$$\alpha_p \alpha_u \alpha_1 \beta_3 \beta_T k_1 k_2 - \alpha_u \alpha_T \alpha_3 \beta_s \beta_1 k_1 k_5 - \alpha_T \alpha_3 \beta_p \beta_u \beta_s \beta_1 k_5 > 0. \quad [4]$$

**1.3. Nondimensionalization.** In order to nondimensionalize the model we introduce stress-induced production changes of preSUMO, E1, E2, E3, ULP, and DeSI. Specifically, for their corresponding subscripts $k \in \{p, u, 1, 2, 3, D\}$, we suppose the production rate $\alpha_k$ takes the form

$$\alpha_k = \begin{cases} \alpha_{k,\text{ns}}, & \text{without stress,} \\ \alpha_{k,\text{s}}, & \text{with stress.} \end{cases} \quad [5]$$

We note that we assume that the production rate of the target protein, $\alpha_T$, is unaffected by stress.

We nondimensionalize Eq. (2) using unstressed steady-state solutions (Eq. (3), Eq. (5)):

$$[x] = [x]_{\text{SS,ns}} \, [x]^*, \quad [6]$$

where asterisks are used to denote the nondimensionalized variables. We consider time relative to the timescale of SUMO degradation, via

$$\tau = \beta_s t. \quad [7]$$

This rescaling yields the following nondimensional parameter groups:

$$\begin{aligned} &\gamma_i = \frac{\beta_i}{\beta_s}, \quad C_k = \frac{\alpha_{k,\text{s}}}{\alpha_{k,\text{ns}}}, \\ &K_1 = \frac{k_1[\text{ULP}]_{\text{SS,ns}}}{\beta_s}, \quad K_2 = \frac{k_2[\text{E1}]_{\text{SS,ns}}}{\beta_s}, \quad K_3 = \frac{k_3[\text{E2}]_{\text{SS,ns}}}{\beta_s}, \quad K_4 = \frac{k_4[\text{target}]_{\text{SS,ns}}}{\beta_s}, \\ &K_5 = \frac{k_5[\text{E3}]_{\text{SS,ns}}}{\beta_s}, \quad K_6 = \frac{k_6[\text{target}]_{\text{SS,ns}}}{\beta_s}, \quad K_7 = \frac{k_7[\text{ULP}]_{\text{SS,ns}}}{\beta_s}, \quad K_8 = \frac{k_8[\text{DeSI}]_{\text{SS,ns}}}{\beta_s}, \\ &R_{s,1} = \frac{[\text{SUMO}]_{\text{SS,ns}}}{[\text{E1}]_{\text{SS,ns}}}, \quad R_{s,T} = \frac{[\text{SUMO}]_{\text{SS,ns}}}{[\text{target}]_{\text{SS,ns}}}, \quad R_{T,2} = \frac{[\text{target}]_{\text{SS,ns}}}{[\text{E2}]_{\text{SS,ns}}}, \quad R_{T,3} = \frac{[\text{target}]_{\text{SS,ns}}}{[\text{E3}]_{\text{SS,ns}}}, \end{aligned} \quad [8]$$

for $i \in \{p, u, 1, 2, 3, T, D\}$ and $k \in \{p, u, 1, 2, 3, D\}$.

The parameters $\gamma_i$ represent ratios between the degradation rates, $\beta_i$, to the SUMO degradation rate, $\beta_s$. The parameters $C_k$ represent the fold changes in production rates of the relevant species. The parameters $K_j$ ($j \in \{1, 2, ..., 8\}$), are nondimensional binding rates obtained by scaling the dimensional rates $k_j$ by the appropriate unstressed dimensional steady-state concentration and the SUMO degradation rate $\beta_s$. Therefore, each $K_j$ quantifies the relative strength of each reaction compared to SUMO degradation. Finally, the parameters $R_{v,w}$ for $(v, w) \in \{(s, 1), (s, T), (T, 2), (T, 3)\}$ denote the ratios between the unstressed steady-state concentrations of key pathway components.

Using Eq. (6)-Eq. (8), the nondimensional system of ODEs (Eq. (2)) under stress yields:

$$\frac{\mathrm{d}}{\mathrm{d}\tau}[\text{preSUMO}]^* = \gamma_p(C_p - [\text{preSUMO}]^*) + K_1(C_p - [\text{preSUMO}]^*[\text{ULP}]^*), \tag{9a}$$

$$\frac{\mathrm{d}}{\mathrm{d}\tau}[\text{ULP}]^* = \gamma_u(C_u - [\text{ULP}]^*), \tag{9b}$$

$$\begin{aligned}\frac{\mathrm{d}}{\mathrm{d}\tau}[\text{SUMO}]^* &= [\text{preSUMO}]^*[\text{ULP}]^* - K_2[\text{E1}]^*[\text{SUMO}]^* + \frac{K_2K_7}{K_7+K_8}[\text{target}\cdot\text{SUMO}]^*[\text{ULP}]^* \\ &+ \frac{K_2K_8}{K_7+K_8}[\text{target}\cdot\text{SUMO}]^*[\text{DeSI}]^* - [\text{SUMO}]^*,\end{aligned} \tag{9c}$$

$$\frac{\mathrm{d}}{\mathrm{d}\tau}[\text{E1}]^* = \gamma_1(C_1 - [\text{E1}]^*) + K_2R_{s,1}([\text{E2}]^*[\text{E1}\cdot\text{SUMO}]^* - [\text{E1}]^*[\text{SUMO}]^*), \tag{9d}$$

$$\frac{\mathrm{d}}{\mathrm{d}\tau}[\text{E1}\cdot\text{SUMO}]^* = K_3([\text{E1}]^*[\text{SUMO}]^* - [\text{E2}]^*[\text{E1}\cdot\text{SUMO}]^*), \tag{9e}$$

$$\begin{aligned}\frac{\mathrm{d}}{\mathrm{d}\tau}[\text{E2}]^* &= \gamma_2(C_2 - [\text{E2}]^*) + K_2R_{s,T}R_{T,2}([\text{target}]^*[\text{E2}\cdot\text{SUMO}]^* - [\text{E2}]^*[\text{E1}\cdot\text{SUMO}]^*) \\ &+ K_5R_{T,2}([\text{E2}\cdot\text{SUMO}]^*[\text{E3}\cdot\text{target}]^* - [\text{target}]^*[\text{E2}\cdot\text{SUMO}]^*),\end{aligned} \tag{9f}$$

$$\begin{aligned}\frac{\mathrm{d}}{\mathrm{d}\tau}[\text{E2}\cdot\text{SUMO}]^* &= \frac{K_2K_4R_{s,T}}{K_2R_{s,T}-K_5}([\text{E2}]^*[\text{E1}\cdot\text{SUMO}]^* - [\text{target}]^*[\text{E2}\cdot\text{SUMO}]^*) \\ &+ \frac{K_4K_5}{K_2R_{s,T}-K_5}([\text{target}]^*[\text{E2}\cdot\text{SUMO}]^* - [\text{E2}\cdot\text{SUMO}]^*[\text{E3}\cdot\text{target}]^*),\end{aligned} \tag{9g}$$

$$\begin{aligned}\frac{\mathrm{d}}{\mathrm{d}\tau}[\text{target}]^* &= \gamma_T(1 - [\text{target}]^*) - K_2R_{s,T}[\text{target}]^*[\text{E2}\cdot\text{SUMO}]^* \\ &+ K_5([\text{target}]^*[\text{E2}\cdot\text{SUMO}]^* - [\text{E3}]^*[\text{target}]^*) + \frac{K_2K_7}{K_7+K_8}[\text{target}\cdot\text{SUMO}]^*[\text{ULP}]^* \\ &+ \frac{K_2K_8}{K_7+K_8}[\text{target}\cdot\text{SUMO}]^*[\text{DeSI}]^*,\end{aligned} \tag{9h}$$

$$\frac{\mathrm{d}}{\mathrm{d}\tau}[\text{E3}]^* = \gamma_3(C_3 - [\text{E3}]^*) + K_5R_{T,3}([\text{E2}\cdot\text{SUMO}]^*[\text{E3}\cdot\text{target}]^* - [\text{E3}]^*[\text{target}]^*), \tag{9i}$$

$$\frac{\mathrm{d}}{\mathrm{d}\tau}[\text{E3}\cdot\text{target}]^* = \frac{K_6(K_2R_{s,T}-K_5)}{K_4}([\text{E3}]^*[\text{target}]^* - [\text{E2}\cdot\text{SUMO}]^*[\text{E3}\cdot\text{target}]^*), \tag{9j}$$

$$\begin{aligned}\frac{\mathrm{d}}{\mathrm{d}\tau}[\text{target}\cdot\text{SUMO}]^* &= \frac{(K_2R_{s,T}-K_5)(K_7+K_8)}{K_2R_{s,T}}[\text{target}]^*[\text{E2}\cdot\text{SUMO}]^* \\ &+ \frac{K_5(K_7+K_8)}{K_2R_{s,T}}[\text{E2}\cdot\text{SUMO}]^*[\text{E3}\cdot\text{target}]^* - K_7[\text{target}\cdot\text{SUMO}]^*[\text{ULP}]^* \\ &- K_8[\text{target}\cdot\text{SUMO}]^*[\text{DeSI}]^*,\end{aligned} \tag{9k}$$

$$\frac{\mathrm{d}}{\mathrm{d}\tau}[\text{DeSI}]^* = \gamma_D(C_D - [\text{DeSI}]^*). \tag{9l}$$

**Summary.** The system in Eq. (9) describes the dynamics of the nondimensional species concentrations given a stress-induced change in production for preSUMO, E1, E2, E3, ULP, and DeSI. In section 2, we use

Eq. (9) to derive a function that predicts the fold change of SUMOylated target using SUMO Cell Atlas data (1). In section 3, we extend this framework to investigate the effects of step changes in production on the dynamics of the fold change of SUMOylated target. Finally, in section 4, we use Eq. (9) together with time-dependent experimental data to predict the dynamic fold change of the SUMOylated target.

### 2. Computing the steady-state fold change of SUMOylated target

Increased SUMOylation of target proteins is a key component of plant stress response. We therefore investigate how stress-induced changes in species within the SUMO cycle affect the concentration of SUMOylated target, [target · SUMO], as indicated by previous experimental studies showing increased SUMOylation following stress (2–5).

To quantify this response, we use data from the SUMO Cell Atlas (1), which provide fold changes in SUMO, ULP, E1, E2, E3, and DeSI isoforms before and 3 hours after stress across different regions of the *Arabidopsis* root. Our aim is to integrate these data into a mathematical framework to predict the resulting fold change in SUMOylated target.

In section 2.1, we derive a function for the steady-state fold change of SUMOylated target in terms of the fold changes of SUMO, ULP, E1, and DeSI, using the nondimensional ODE model (Eq. (9)). In section 2.2, we convert the experimentally measured isoform-level fold changes into species-level quantities that can be used as inputs to the model. Since the parameters in Eq. (8) are unknown, section 2.3 outlines our parameter sampling approach. Finally, in section 2.4, we analyze how prescribed changes in SUMO, ULP, E1, and DeSI influence the steady-state fold change of SUMOylated target, and assess the role of parameter uncertainty in shaping these predictions.

**2.1. Deriving the SUMOylated target steady-state fold change.** In this subsection, we derive an expression for the steady-state fold change of SUMOylated target in terms of the experimentally measured fold changes in SUMO, ULP, E1, and DeSI.

To do this, we assume that stress-induced changes in species levels arise from changes in their production rates. We solve the ODEs in Eq. (9) for steady state (section 2.1.1) and use the resulting solution to obtain a steady-state fold change function for SUMOylated target (section 2.1.2).

***2.1.1. Nondimensional model steady-state solutions.*** Let $[x]^*_{\mathrm{SS,ns}}$ and $[x]^*_{\mathrm{SS,s}}$ denote the steady states of Eq. (9) without and with stress, respectively.

By the choice of nondimensionalisation in Eq. (6), we have

$$[x]^*_{\mathrm{SS,ns}} = 1, \quad \text{for all } x \in X.$$

Thus, each stressed steady-state value $[x]^*_{\mathrm{SS,s}}$ can be interpreted directly as the fold change in $[x]$ following stress. We use this property in section 2.1.2 to derive the fold-change function for SUMOylated target.

To find the stressed steady state, we solve Eq. (9) for steady state and obtain:

$$[\mathrm{preSUMO}]^*_{\mathrm{SS,s}} = \frac{C_p(K_1 + \gamma_p)}{C_u K_1 + \gamma_p}, \tag{10a}$$

$$[\mathrm{ULP}]^*_{\mathrm{SS,s}} = C_u, \tag{10b}$$

$$[\mathrm{SUMO}]^*_{\mathrm{SS,s}} = \frac{C_p C_u (K_1 + \gamma_p)}{C_u K_1 + \gamma_p}, \tag{10c}$$

$$[\mathrm{E1}]^*_{\mathrm{SS,s}} = C_1, \tag{10d}$$

$$[\mathrm{E1} \cdot \mathrm{SUMO}]^*_{\mathrm{SS,s}} = \frac{C_p C_u C_1 (K_1 + \gamma_p)}{C_2 (C_u K_1 + \gamma_p))}, \tag{10e}$$

$$[\mathrm{E2}]^*_{\mathrm{SS,s}} = C_2, \tag{10f}$$

$$[\text{E2} \cdot \text{SUMO}]^*_{\text{SS,s}} = \frac{C_u(C_pC_1K_2R_{s,T}\gamma_p + K_1(C_pC_1K_2R_{s,T} - C_3K_5)) - C_3K_5\gamma_p}{(C_uK_1 + \gamma_p)(K_2R_{s,T} - K_5)}, \quad [10g]$$

$$[\text{target}]^*_{\text{SS,s}} = 1, \quad [10h]$$

$$[\text{E3}]^*_{\text{SS,s}} = C_3, \quad [10i]$$

$$[\text{E3} \cdot \text{target}]^*_{\text{SS,s}} = \frac{C_3(C_uK_1 + \gamma_p)(K_2R_{s,T} - K_5)}{C_u(C_pC_1K_2R_{s,T}\gamma_p + K_1(C_pC_1K_2R_{s,T} - C_3K_5)) - C_3K_5\gamma_p}, \quad [10j]$$

$$[\text{target} \cdot \text{SUMO}]^*_{\text{SS,s}} = \frac{C_pC_uC_1(K_1 + \gamma_p)(K_7 + K_8)}{(C_uK_1 + \gamma_p)(C_uK_7 + C_DK_8)}, \quad [10k]$$

$$[\text{DeSI}]^*_{\text{SS,s}} = C_D. \quad [10l]$$

To ensure that the stressed steady state is positive and finite, the conditions

$$K_2R_{s,T} - K_5 > 0, \quad [11a]$$

and

$$C_u(C_pC_1K_2R_{s,T}\gamma_p + K_1(C_pC_1K_2R_{s,T} - C_3K_5)) - C_3K_5\gamma_p > 0, \quad [11b]$$

must hold.

From Eq. (10), the steady-state concentration of free target protein remains unchanged under stress. The stressed steady-state levels of ULP, E1, E2, E3, and DeSI scale directly with their respective production fold changes ($C_u, C_1, C_2, C_3, C_D$), while other species depend on both production changes and the nondimensional parameter groups defined in Eq. (8).

***2.1.2. Fold change derivation.*** From Eq. (10k), the fold change of SUMOylated target depends on the production fold changes $C_p$, $C_u$, $C_1$, and $C_D$. The quantities $C_u$, $C_1$, and $C_D$ correspond to the fold changes in ULP, E1, and DeSI and can be directly computed using the SUMO Cell Atlas data (1). In contrast, the production fold change in preSUMO, $C_p$, is not directly measured; instead, the data are assumed to provide the fold change in mature SUMO isoforms. Thus, we denote the fold change of total SUMO by $C_s$.

From Eq. (10c), the steady-state fold change of SUMO satisfies

$$C_s = [\text{SUMO}]^*_{\text{SS,s}} = \frac{C_pC_u(K_1 + \gamma_p)}{C_uK_1 + \gamma_p}. \quad [12]$$

This expression provides a relationship between the unknown production fold change $C_p$ and the observable quantity $C_s$. Rearranging Eq. (12) gives

$$C_p = \frac{C_s(C_uK_1 + \gamma_p)}{C_u(K_1 + \gamma_p)}. \quad [13]$$

We can therefore eliminate the unknown quantity $C_p$ by substituting Eq. (13) into the non-dimensional steady-state expression for SUMOylated target (Eq. (10k)). This yields a fold change function that depends on experimentally measurable fold changes:

$$F(C_s, C_u, C_1, C_D, K_7, K_8) = \frac{C_sC_1(K_7 + K_8)}{C_uK_7 + C_DK_8}. \quad [14]$$

The parameters $K_7$ and $K_8$ represent the relative contributions of ULP- and DeSI-mediated deSUMOylation, respectively. As these parameters are unknown, we explore their effects using parameter sampling (see section 2.3 for details).

Eq. (14) therefore provides a direct mapping from experimentally measured fold changes to the predicted fold change of SUMOylated target. Thus, although $[\text{target} \cdot \text{SUMO}]^*_{\text{SS,s}}$ depends on the unknown production change $C_p$, the relationship Eq. (12) allows us to rewrite the result entirely in terms of the measured SUMO fold change $C_s$.

Throughout the manuscript, we use Eq. (14) together with experimentally measured fold changes in the SUMO, ULP, E1, and DeSI species, enabling quantitative, tissue-specific predictions of how SUMOylation levels respond to salt and osmotic stress (see Fig. 2 D, E).

**2.2. Calculating the species fold changes.** The SUMO Cell Atlas (1) reports fold changes for SUMO, ULP, E1, E2, E3, and DeSI isoforms 3 hours after stress relative to the unstressed levels. To apply the SUMOylated target steady-state fold change function (Eq. (14)), derived in section 2.1, we assume that the SUMO system is at steady state both before and after stress. Under this assumption, the pre- and post-stress states correspond to the unstressed and stressed steady states of the model, respectively.

To incorporate these data into the model, we must map isoform-level measurements onto the species-level fold changes used in Eq. (14). In this subsection, we derive the required mappings, which depend on whether a species consists of multiple isoforms or is formed through the assembly of a protein complex.

***2.2.1. Using data to estimate the fold changes of SUMO, ULP, E3, and DeSI.*** The species representing SUMO, ULP, E3, and DeSI each consist of multiple isoforms (Table S3). Let $[a] \in \{[\text{SUMO}], [\text{ULP}], [\text{E3}], [\text{DeSI}]\}$, and let $[b_1], \ldots, [b_n]$ denote the contributing isoforms. Then

$$[a] \ = \ \sum_{y=1}^{n} [b_y]. \tag{15}$$

We denote unstressed and stressed steady-state concentrations by $[\,\cdot\,]_{\text{SS,ns}}$ and $[\,\cdot\,]_{\text{SS,s}}$, respectively. The corresponding fold changes are defined by

$$[b_y]^*_{\text{SS,s}} \ = \ \frac{[b_y]_{\text{SS,s}}}{[b_y]_{\text{SS,ns}}}, \qquad [a]^*_{\text{SS,s}} \ = \ \frac{[a]_{\text{SS,s}}}{[a]_{\text{SS,ns}}}. \tag{16}$$

The SUMO Cell Atlas reports $\log_2$ fold changes. If $D_{b_y}$ denotes the reported value for isoform $[b_y]$, then

$$[b_y]^*_{\text{SS,s}} = 2^{D_{b_y}}. \tag{17}$$

Using fluorescence data (Table S3), we estimate the baseline proportion of $[a]$ contributed by $[b_y]$ under unstressed conditions,

$$P_{b_y} \ = \ \frac{[b_y]_{\text{SS,ns}}}{[a]_{\text{SS,ns}}}, \qquad 0 < P_{b_y} \leq 1, \quad \sum_{i=1}^{n} P_{b_y} = 1. \tag{18}$$

Combining Eq. (15)–Eq. (18) yields

$$[a]^*_{\text{SS,s}} = \frac{[a]_{\text{SS,s}}}{[a]_{\text{SS,ns}}} = \frac{\sum_{y=1}^{n} [b_y]_{\text{SS,s}}}{[a]_{\text{SS,ns}}} = \frac{\sum_{y=1}^{n} [b_y]^*_{\text{SS,s}} [b_y]_{\text{SS,ns}}}{[a]_{\text{SS,ns}}} = \sum_{y=1}^{n} P_{b_y} \, [b_y]^*_{\text{SS,s}}, \tag{19}$$

so that the species-level fold change is a weighted sum of isoform-level fold changes.

***2.2.2. Using data to estimate the fold change of E2.*** E2 has a single isoform (SCE1), so its fold change is given directly by the data:

$$[\text{E2}]^*_{\text{SS,s}} = [\text{SCE1}]^*_{\text{SS,s}}, \tag{20}$$

after converting from $\log_2$ values using Eq. (17).

***2.2.3. Using data to estimate the fold change of E1.*** The previous approaches do not apply to E1, which is a heterodimer composed of SAE2 and either SAE1a or SAE1b. The concentration of assembled E1 heterodimer is given by

$$[\text{E1}] = [\text{SAE1a} \cdot \text{SAE2}] + [\text{SAE1b} \cdot \text{SAE2}]. \tag{21}$$

To obtain the fold change of E1 we consider the following reaction scheme:

$$\text{SAE1a} + \text{SAE2} \underset{q_{1r}}{\overset{q_{1f}}{\rightleftharpoons}} \text{SAE1a} \cdot \text{SAE2} \tag{22a}$$

$$\mathrm{SAE1b} + \mathrm{SAE2} \underset{q_{2r}}{\overset{q_{2f}}{\rightleftharpoons}} \mathrm{SAE1b} \cdot \mathrm{SAE2} \qquad [22b]$$

$$\emptyset \xrightarrow{\alpha_{\mathrm{SAE1a}}} \mathrm{SAE1a} \xrightarrow{\beta_{\mathrm{SAE1a}}} \emptyset \qquad [22c]$$

$$\emptyset \xrightarrow{\alpha_{\mathrm{SAE1b}}} \mathrm{SAE1b} \xrightarrow{\beta_{\mathrm{SAE1b}}} \emptyset \qquad [22d]$$

$$\emptyset \xrightarrow{\alpha_{\mathrm{SAE2}}} \mathrm{SAE2} \xrightarrow{\beta_{\mathrm{SAE2}}} \emptyset. \qquad [22e]$$

Applying the law of mass action to Eq. (22) yields a system of ODEs which we solve for steady state to obtain:

$$[\mathrm{SAE1a}]_{\mathrm{SS}} = \alpha_{\mathrm{SAE1a}}/\beta_{\mathrm{SAE1a}}, \qquad [23a]$$

$$[\mathrm{SAE1b}]_{\mathrm{SS}} = \alpha_{\mathrm{SAE1b}}/\beta_{\mathrm{SAE1b}}, \qquad [23b]$$

$$[\mathrm{SAE2}]_{\mathrm{SS}} = \alpha_{\mathrm{SAE2}}/\beta_{\mathrm{SAE2}}, \qquad [23c]$$

$$[\mathrm{SAE1a} \cdot \mathrm{SAE2}]_{\mathrm{SS}} = \frac{q_{1f}\alpha_{\mathrm{SAE1a}}\alpha_{\mathrm{SAE2}}}{q_{1r}\beta_{\mathrm{SAE1a}}\beta_{\mathrm{SAE2}}} = \frac{q_{1f}}{q_{1r}}[\mathrm{SAE1a}]_{\mathrm{SS}}[\mathrm{SAE2}]_{\mathrm{SS}}, \qquad [23d]$$

$$[\mathrm{SAE1b} \cdot \mathrm{SAE2}]_{\mathrm{SS}} = \frac{q_{2f}\alpha_{\mathrm{SAE1b}}\alpha_{\mathrm{SAE2}}}{q_{2r}\beta_{\mathrm{SAE1b}}\beta_{\mathrm{SAE2}}} = \frac{q_{2f}}{q_{2r}}[\mathrm{SAE1b}]_{\mathrm{SS}}[\mathrm{SAE2}]_{\mathrm{SS}}. \qquad [23e]$$

For each complex, we assume that its binding and unbinding rates are equal, such that $q_{1f} = q_{1r}$ for Eq. (22a) and $q_{2f} = q_{2r}$ for Eq. (22b), while allowing the rates to differ between the two complexes. Assuming $q_{1f} = q_{1r}$ and $q_{2f} = q_{2r}$ and using Eq. (21), Eq. (23d), and Eq. (23e), the steady-state concentration of E1 simplifies to

$$[\mathrm{E1}]_{\mathrm{SS}} = [\mathrm{SAE2}]_{\mathrm{SS}}\big([\mathrm{SAE1a}]_{\mathrm{SS}} + [\mathrm{SAE1b}]_{\mathrm{SS}}\big). \qquad [24]$$

Assuming stress-induced production changes as defined in Eq. (5), Eq. (24) implies that the fold change of the E1 family is

$$[\mathrm{E1}]^*_{\mathrm{SS,s}} = \frac{[\mathrm{E1}]_{\mathrm{SS,s}}}{[\mathrm{E1}]_{\mathrm{SS,ns}}} = \frac{[\mathrm{SAE2}]_{\mathrm{SS,s}}\big([\mathrm{SAE1a}]_{\mathrm{SS,s}} + [\mathrm{SAE1b}]_{\mathrm{SS,s}}\big)}{[\mathrm{SAE2}]_{\mathrm{SS,ns}}\big([\mathrm{SAE1a}]_{\mathrm{SS,ns}} + [\mathrm{SAE1b}]_{\mathrm{SS,ns}}\big)}. \qquad [25]$$

Using the fluorescence data (Table S3) and Eq. (18), we estimate the baseline fractions of the two SAE1 isoforms as

$$[\mathrm{SAE1a}]_{\mathrm{SS,ns}} = 0.52\,[\mathrm{SAE1}]_{\mathrm{SS,ns}}, \qquad [26a]$$

$$[\mathrm{SAE1b}]_{\mathrm{SS,ns}} = 0.48\,[\mathrm{SAE1}]_{\mathrm{SS,ns}}, \qquad [26b]$$

where $[\mathrm{SAE1}]_{\mathrm{SS,ns}} = [\mathrm{SAE1a}]_{\mathrm{SS,ns}} + [\mathrm{SAE1b}]_{\mathrm{SS,ns}}$.

Using the isoform-level fold change definition in Eq. (16) and substituting Eq. (26), we obtain

$$\begin{aligned}[\mathrm{E1}]^*_{\mathrm{SS,s}} &= [\mathrm{SAE2}]^*_{\mathrm{SS,s}}\,\frac{0.52\,[\mathrm{SAE1}]_{\mathrm{SS,ns}}[\mathrm{SAE1a}]^*_{\mathrm{SS,s}} + 0.48\,[\mathrm{SAE1}]_{\mathrm{SS,ns}}[\mathrm{SAE1b}]^*_{\mathrm{SS,s}}}{[\mathrm{SAE1}]_{\mathrm{SS,ns}}} \\ &= [\mathrm{SAE2}]^*_{\mathrm{SS,s}}\big(0.52\,[\mathrm{SAE1a}]^*_{\mathrm{SS,s}} + 0.48\,[\mathrm{SAE1b}]^*_{\mathrm{SS,s}}\big). \qquad [27]\end{aligned}$$

Thus, the E1 fold change is a weighted combination of SAE1 isoform fold changes, scaled by SAE2 fold change.

In summary, the experimental data provide isoform-level fold changes (Tables S1 and S2) together with baseline abundance estimates (Table S3). Using Eq. (19), Eq. (20), and Eq. (27), we convert these into species-level fold changes for SUMO, ULP, E1, E2, E3, and DeSI, enabling direct incorporation of the SUMO Cell Atlas data into the model. These expressions are used to generate the heatmaps shown in Fig. 2B,C from the data in Tables S1 and S2.

#### 2.3. Parameter surveys.

Except for the fold-change parameters ($C_p, C_u, C_1, C_2, C_3$, and $C_D$), all model parameters (Eq. (8)) are unknown. To assess how uncertainty in the deSUMOylation rates influences model predictions, we explored the parameter space defined by $K_7, K_8 \in [1/100, 100]$, assuming that all remaining parameters satisfy the conditions in Eq. (11).

To generate parameter sets, the interval $[1/100, 100]$ is sampled at 100 logarithmically spaced values for each parameter. The Cartesian product of these values is used to construct all possible $(K_7, K_8)$ pairs, yielding 10,000 parameter sets in total. Each parameter set is subsequently used to evaluate the steady-state fold change of SUMOylated target (Eq. (14)).

This range enables broad exploration of parameter space without imposing strong prior assumptions on the relative rates of ULP- and DeSI-mediated deSUMOylation. The resulting distributions of model outputs are displayed using boxplots. These simulations are used in section 2.4 and to generate the predictions shown in Fig. 2D,E using the data in Tables S1 and S2.

#### 2.4. Analyzing the steady-state fold change of SUMOylated target.

To investigate how changes in individual species influence the steady-state fold change of SUMOylated target (Eq. (14)) we prescribe fold changes in SUMO, ULP, E1, and DeSI. This allows us to examine how these inputs interact and to determine the resulting range of possible responses. In particular, we identify when the steady-state fold change of SUMOylated target depends on deSUMOylation rates $(K_7, K_8)$, and quantify how uncertainty in these parameters contributes to variability in model predictions.

##### 2.4.1. Effect of SUMO and E1 on the fold change of SUMOylated target.

From Eq. (14), if SUMO and E1 vary while ULP and DeSI remain unchanged, the steady-state fold change of the SUMOylated target is

$$F(C_s, 1, C_1, 1, K_7, K_8) = C_s C_1, \tag{28}$$

with special cases $F(C_s, 1, 1, 1, K_7, K_8) = C_s$ and $F(1, 1, C_1, 1, K_7, K_8) = C_1$. Thus, SUMO and E1 influence the fold change directly and proportionally, with no dependence on $K_7$ and $K_8$.

##### 2.4.2. Effect of ULP and DeSI on the fold change of SUMOylated target.

If ULP varies while SUMO, E1, and DeSI remain unchanged, the predicted steady-state fold change of the SUMOylated target is

$$F(1, C_u, 1, 1, K_7, K_8) = \frac{K_7 + K_8}{(C_u K_7 + K_8)}, \tag{29}$$

which depends explicitly on $K_7$ and $K_8$. The qualitative behavior of Eq. (29) is determined by whether ULP decreases or increases under stress:

$$F(1, C_u, 1, 1, K_7, K_8) \begin{cases} > 1, & \text{if } 0 < C_u < 1, \\ = 1, & \text{if } C_u = 1, \\ < 1, & \text{if } C_u > 1. \end{cases}$$

Thus, increasing ULP reduces the steady-state fold change of the SUMOylated target, while decreasing ULP increases it.

To quantify the effect of parameter uncertainty, we evaluate Eq. (29) over all admissible values of $K_7$ and $K_8$, obtaining

$$\min_{\{K_7, K_8\}} F(1, C_u, 1, 1, K_7, K_8) = \begin{cases} 1, & 0 < C_u < 1, \\ 1/C_u, & C_u \geq 1, \end{cases} \tag{30a}$$

$$\max_{\{K_7, K_8\}} F(1, C_u, 1, 1, K_7, K_8) = \begin{cases} 1/C_u, & 0 < C_u < 1, \\ 1, & C_u \geq 1. \end{cases} \tag{30b}$$

Hence,

$$\text{Range}(1, C_u, 1, 1) = \frac{|C_u - 1|}{C_u}. \qquad [30c]$$

These bounds agree with the parameter sweep in Fig. S1A and show that changes in ULP levels introduce an explicit dependence on deSUMOylation rates in the predicted steady-state fold change of SUMOylated target. From Eq. (30c) we see that the magnitude of $C_u$ determines the extent of this variability across the parameter sets. The same analysis applies to DeSI (Fig. S1B), demonstrating that it similarly drives parameter-dependent variability.

Allowing both ULP and DeSI to vary simultaneously gives

$$F(1, C_u, 1, C_D, K_7, K_8) = \frac{(K_7 + K_8)}{(C_u K_7 + C_D K_8)}. \qquad [31]$$

Evaluating Eq. (31) over $K_7$ and $K_8$ yields

$$\min_{\{K_7, K_8\}} = F(1, C_u, 1, C_D, K_7, K_8) = \frac{1}{\max(C_u, C_D)}, \qquad [32a]$$

$$\max_{\{K_7, K_8\}} F(1, C_u, 1, C_D, K_7, K_8) = \frac{1}{\min(C_u, C_D)}. \qquad [32b]$$

Thus,

$$\text{Range}(1, C_u, 1, C_D) = \frac{|C_D - C_u|}{C_D C_u}. \qquad [32c]$$

These bounds are consistent with the numerical simulations seen in Fig. S2 and show that variability in the predicted steady-state fold change of SUMOylated target throughout the parameter space is governed by the difference between ULP and DeSI fold changes: when $C_u = C_D$, the model does not depend on deSUMOylation rates, while increasing disparity between $C_u$ and $C_D$ increases the range of possible outcomes.

***2.4.3. Combined influence of all SUMO components on the steady-state fold change of SUMOylated target.*** Using Eq. (31), we rewrite Eq. (14) as

$$F(C_s, C_u, C_1, C_D, K_7, K_8) = C_s C_1 \, F(1, C_u, 1, C_D, K_7, K_8), \qquad [33]$$

showing that SUMO and E1 act multiplicatively, while ULP and DeSI modulate the result through the parameter-dependent term $F(1, C_u, 1, C_D, K_7, K_8)$.

Evaluating Eq. (33) over all admissible deSUMOylation rates yields

$$\min_{\{K_7, K_8\}} F(C_s, C_u, C_1, C_D, K_7, K_8) = \frac{C_s C_1}{\max(C_u, C_D)}, \qquad [34a]$$

$$\max_{\{K_7, K_8\}} F(C_s, C_u, C_1, C_D, K_7, K_8) = \frac{C_s C_1}{\min(C_u, C_D)}, \qquad [34b]$$

and therefore

$$\text{Range}(C_s, C_u, C_1, C_D) = \frac{C_s C_1 \, |C_D - C_u|}{C_D C_u}. \qquad [34c]$$

These analytical bounds agree with the parameter sweep results (Figs. S3). They show that SUMO and E1 scale the variability of the steady-state fold change of SUMOylated target introduced by ULP and DeSI: when $C_s C_1 > 1$, parameter-dependent variation throughout the parameter space is amplified, whereas when $C_s C_1 < 1$, it is attenuated.

**Summary.** In this section, we developed a mathematical framework to predict the steady-state fold change of SUMOylated target in response to stress, using fold change data from the SUMO Cell Atlas. Starting from the nondimensional ODE model, we derived an explicit expression for the fold change in SUMOylated target in terms of the fold changes of SUMO, ULP, E1, and DeSI. We then constructed mappings from isoform-level measurements to species-level fold changes, enabling direct incorporation of experimental data into the model.

To account for uncertainty in model parameters, we performed parameter surveys. Our analysis shows that SUMO and E1 contribute multiplicatively and independently of parameter values, while ULP and DeSI introduce parameter-dependent variability. In particular, the steady-state fold change is independent of parameter uncertainty when ULP and DeSI fold changes are equal, and the extent of variability increases with the disparity between their fold changes.

Overall, this framework provides a systematic way to link experimentally observed changes in the SUMO cycle to predicted changes in SUMOylated target, while explicitly characterizing the role of parameter uncertainty.

### 3. Simulation of model dynamics under prescribed step changes

Previous experimental studies indicate that stress induces an increase in SUMOylation (2–5), leading to elevated levels of SUMOylated target. However, when the SUMO Cell Atlas data (Fig. 2B,C) are incorporated into the steady-state fold-change formula (Eq. (14)), the predicted response is not uniformly increasing: SUMOylation decreases in the elongation zone following salt treatment (Fig. 2D) and remains largely unchanged in the meristem following mannitol treatment (Fig. 2E).

These results suggest that steady-state analysis alone may not fully capture the stress response. In particular, SUMOylation may increase transiently following stress before relaxing to the stressed steady state.

To investigate this possibility, we analyze the dynamic response of the model to prescribed step changes in production. The experimental data provide dynamic fold changes for SUMO, ULP, E2, and DeSI (Fig. 3), whereas E1 and E3 exhibit negligible changes following stress. Thus, we examine step changes in preSUMO, ULP, E2, and DeSI. As SUMO has no associated production term in the model, changes in SUMO are implemented through step changes in preSUMO. Simulations are performed in dimensional time to enable direct comparison with the experimental measurements presented in section 4. This analysis allows us to determine whether transient SUMOylation dynamics can explain the apparent discrepancy between experimental observations and steady-state predictions. While the main text focuses on the predicted dynamics of SUMOylated target, this section presents the corresponding dynamics of the remaining model species together with a sensitivity analysis of the SUMOylated target response to perturbations in the model parameters.

#### 3.1. Methodology.

We consider the system Eq. (9) and recover dimensional time by multiplying each equation by $\beta_s$. As $\beta_s$ sets the intrinsic timescale of the system, varying it allows us to investigate how timescale influences the model dynamics. Simulations are performed over the interval (0 h, 20 h), ensuring that the fastest systems reach the stressed steady state within the simulation window.

Simulations are initialized at the unstressed steady state,

$$[x]^* = 1 \quad \text{at} \quad t = 0 \quad \text{for all} \quad x \in X, \tag{35}$$

and solved numerically using `solve_ivp` from `SciPy.integrate` (6).

We consider the parameter set

$$\mu = \{\gamma_p, \gamma_u, \gamma_1, \gamma_2, \gamma_3, \gamma_T, \gamma_D, \beta_s, K_1, K_2, K_3, K_4, K_5, K_6, K_7, K_8, R_{s,1}, R_{s,T}, R_{T,2}, R_{T,3}\},$$

consisting of the nondimensional parameters in Eq. (8) together with $\beta_s$. Since the dimensional parameter values are unknown, we define a baseline parameter set, denoted $\mu_{g,b}$, by setting $R_{s,T} = 2.5$ and $\beta_s = g$, with all remaining parameters equal to 1. This choice satisfies the constraints in Eq. (11), assumes that all processes within the SUMO cycle operate on comparable timescales, and allows the overall timescale of the system to be varied through $\beta_s$.

To isolate the contribution of individual species, we impose production step changes in preSUMO, ULP, E2, and DeSI. For a prescribed fold change $z \neq 1$, we set $C_h = z$ (Eq. (9)) for $h \in \{p, u, 2, D\}$ while fixing all remaining fold changes at 1. To generate simulations in which the specified component very rapidly evolves to its new steady state (approximating a step-change), we additionally set $\gamma_h = 100$. The resulting parameter set is denoted $\mu_{g,h}$.

To aid interpretation of several parameter perturbations, we also redimensionalize system Eq. (9). The dimensional parameters are recovered from $\mu_{g,h}$ using the definitions in Eq. (8), uniquely determining all parameters except $\alpha_u$, $\alpha_T$, and $\alpha_D$, which are fixed at 1.

**3.2. Response to a preSUMO step change.** A 50% increase in preSUMO production under the parameter sets $\mu_{1,p}$, $\mu_{2,p}$, and $\mu_{3,p}$ (see section 3.1 for details) causes preSUMO to rapidly reach its new steady state owing to the large value of $\gamma_p$ (Fig. S6A). This increase propagates through the SUMOylation pathway, increasing SUMO, E1 · SUMO, E2 · SUMO, and SUMOylated target (Fig. S6D,E,G,K), while E1, E2, and E3 · target decrease as they are incorporated into complexes (Fig. S6C,F,J). ULP and DeSI remain unchanged (Fig. S6B,L) because their dynamics are decoupled from the remainder of the system. Across all simulations, faster intrinsic timescales produce more rapid convergence to the stressed steady state, whereas slower timescales generate larger and more persistent transient increases in SUMOylated target.

To assess the robustness of these predictions, each parameter was varied individually about the baseline parameter sets $\mu_{1,p}$ and $\mu_{3,p}$ by doubling and halving its value. For each perturbation, we examine only the predicted dynamics of SUMOylated target. Parameter sensitivities were qualitatively similar across timescales, with the slower-timescale behavior corresponding to the early-time dynamics of the faster system (Fig. S7; Fig. S8). Consequently, parameter variation for the remaining step-change analyses is presented only for $\mu_{3,h}$.

The fold change of SUMOylated target is largely insensitive to parameters governing preSUMO production, $\gamma_p$, $K_1$ (Fig. S8A,I), and the ULP- and DeSI-mediated deSUMOylation, $\gamma_u$, $\gamma_D$, $K_7$, $K_8$ (Fig. S8B,F,O,P), as these do not alter the stressed steady states under a preSUMO perturbation. In contrast, parameters controlling intermediate SUMOylation reactions, particularly $K_2$, $K_3$, $K_4$, $K_5$, and the initial SUMO-to-target ratio $R_{s,T}$, substantially modify the transient SUMOylation response (Fig. S8J,K,L,M,R). Although these parameters do not alter the qualitative behavior, they change both the magnitude of the transient peak and the rate at which the system approaches steady state. The counterintuitive behaviors involving changes in $K_2$ and $K_5$ arise from changes in the dimensional steady-state concentrations encoded within the nondimensional parameters (Fig. S9, Fig. S10).

Overall, these results indicate that the response to increased preSUMO is robust to most parameter changes. The transient dynamics are governed primarily by the kinetics of the intermediate SUMOylation reactions and by the initial balance between SUMO and target protein, with $R_{s,T}$ exerting the strongest influence on the magnitude and duration of the SUMOylation response.

**3.3. Response to ULP step change.** ULP performs two opposing functions within the SUMO cycle: it promotes preSUMO maturation, increasing the available SUMO pool, while also directly deSUMOylating SUMOylated target. For a 50% increase in ULP production under the parameter sets $\mu_{1,u}$, $\mu_{2,u}$, and $\mu_{3,u}$ (see section 3.1 for details), we see ULP rapidly reaches its new steady state (Fig. S11B), owing to the large value of $\gamma_u$. Increased ULP accelerates preSUMO maturation, reducing preSUMO levels (Fig. S11A) and increasing SUMO, E1 · SUMO, and E2 · SUMO (Fig. S11D,E,G), while free E1, E2, and E3 · target decrease as they are incorporated into complexes (Fig. S11C,F,J). In contrast to the preSUMO perturbation, SUMOylated target initially decreases because of enhanced deSUMOylation before recovering as the larger SUMO pool propagates through the pathway. DeSI remains unchanged because its dynamics are decoupled from the remainder of the system (Fig. S11L). Faster intrinsic timescales accelerate convergence to the stressed steady state, whereas slower systems exhibit a larger and more prolonged transient decrease in SUMOylated target.

To assess the robustness of these predictions, each parameter was doubled and halved about the baseline parameter set $\mu_{3,u}$. For each perturbation, we examine only the predicted dynamics of SUMOylated target. Unlike the preSUMO perturbation, the response is sensitive to parameters governing both SUMO maturation and deSUMOylation. In particular, changes in $\gamma_p$, $K_1$, $K_7$, and $K_8$ modify the stressed steady state of SUMOylated target (Fig. S12A,I,O,P), reflecting the dependence of the steady state on preSUMO maturation and deSUMOylation (see Eq. (10k)). Parameters controlling the intermediate SUMOylation reactions, particularly $K_2$, $K_3$, $K_4$, $K_5$, and the initial SUMO-to-target ratio $R_{s,T}$, primarily influence the transient response by altering both the size of the initial decrease and the rate of recovery (Fig. S12J–M,R). Among these, increases in $K_2$ and $R_{s,T}$ uniquely generate pronounced transient peaks in SUMOylated target, highlighting their importance in shaping the transient response. The seemingly counterintuitive behavior associated with $K_2$ arises from changes in the dimensional steady-state concentrations encoded within the nondimensional parameters (Fig. S9).

Overall, the response to increased ULP reflects the competition between its two roles in the SUMO cycle. Immediately after the perturbation, enhanced deSUMOylation dominates, producing a transient reduction in SUMOylated target. At longer times, accelerated preSUMO maturation increases the SUMO pool, eventually outweighing the direct deSUMOylation effect and driving the system towards a new steady state. The magnitude and duration of the transient are governed primarily by the intermediate SUMOylation reactions and the initial SUMO-to-target ratio, whereas the new stressed steady state is determined by the balance between preSUMO maturation and deSUMOylation.

**3.4. Response to E2 step change.** Unlike preSUMO and ULP, changes in E2 do not alter the steady-state level of SUMOylated target (Eq. (10k)); instead, they regulate the transient response. We therefore considered a 50% increase in E2 production under the parameter sets $\mu_{1,2}$, $\mu_{2,2}$, and $\mu_{3,2}$ (see section 3.1 for details). Owing to the large value of $\gamma_2$, E2 rapidly reaches its new steady state (Fig. S13F). As E2 acts downstream of preSUMO, preSUMO levels remain unchanged (Fig. S13A). Elevated E2 increases the formation of E2 · SUMO, producing a transient increase in SUMOylated target (Fig. S13G,K). Consequently, free E1 increases while E1 · SUMO decreases as SUMO is transferred more rapidly through the conjugation pathway (Fig. S13C,E). SUMO and free target exhibit transient decreases as they are consumed during conjugation before recovering through deSUMOylation (Fig. S13D,H). ULP and DeSI remain unchanged because their dynamics are decoupled from the remainder of the system (Fig. S13B,L). As in the previous perturbations, faster intrinsic timescales shorten the transient response, whereas slower systems exhibit broader and more persistent transient increases in SUMOylated target.

To assess the robustness of these predictions, each parameter was doubled and halved about the baseline parameter set $\mu_{3,2}$. For each perturbation, we examine only the predicted dynamics of SUMOylated target. As E2 acts downstream of preSUMO, changes in E2 do not affect the fold change of preSUMO, thus the fold change of SUMOylated target is insensitive to changes in $\gamma_p$, $K_1$ (Fig. S14A,I). Similarly, perturbations in the production parameters of ULP- and DeSI $\gamma_u$, $\gamma_D$, do not affect the fold change of SUMOylated target (Fig.S14O,P). Consistent with the steady-state analysis, parameter variation does not alter the stressed steady state but instead modifies the magnitude and duration of the transient response. Parameters governing the intermediate SUMOylation reactions, particularly $K_2$, $K_3$, $K_4$, $K_5$, together with the initial SUMO-to-target ratio $R_{s,T}$, have the greatest influence on the transient dynamics (Fig. S14J-M,R). In contrast, parameters governing preSUMO production and the production of ULP and DeSI have little influence on the response. Several seemingly counterintuitive behaviours, particularly those associated with $K_2$, $K_3$, $K_5$, $K_7$, and $K_8$, arise because the corresponding nondimensional parameters also encode the dimensional steady-state concentrations (Figs. S9, S10, S15, S16).

Overall, increased E2 acts primarily as a transient accelerator of SUMOylation rather than changing the final level of SUMOylated target. The transient response is governed mainly by the intermediate SUMOylation reactions and the initial SUMO-to-target ratio.

**3.5. Response to DeSI step change.** Unlike ULP, DeSI functions solely as a deSUMOylating enzyme. We considered the model response to a 50% increase in DeSI production under the parameter sets $\mu_{1,D}$, $\mu_{2,D}$, and $\mu_{3,D}$ (see section 3.1 for details). Owing to the large value of $\gamma_D$, DeSI rapidly reaches its new steady state (Fig. S17L). Increased DeSI produces a monotonic reduction in SUMOylated target (Fig. S17K), consistent with its direct role in promoting deSUMOylation. The resulting release of SUMO and target protein generates transient increases in free SUMO and target (Fig. S17D,H), which propagate through the conjugation pathway, producing transient increases in E1 $\cdot$ SUMO, E2 $\cdot$ SUMO, and E3 $\cdot$ target (Fig. S17E,G,J). ULP remains unchanged because its dynamics are decoupled from the remainder of the system (Fig. S17B). As in the previous perturbations, faster intrinsic timescales accelerate convergence to the stressed steady state, whereas slower systems exhibit broader and more persistent transient responses.

To assess the robustness of these predictions, each parameter was doubled and halved about the baseline parameter set $\mu_{3,D}$. For each perturbation, we examine only the predicted dynamics of SUMOylated target. The response is most sensitive to the deSUMOylation parameters $K_7$ and $K_8$, which directly alter the stressed steady state of SUMOylated target under a DeSI step change (Fig. S18O,P). In contrast, variation in most other parameters has little influence on the predicted dynamics, with only $K_2$ and the initial SUMO-to-target ratio $R_{s,T}$ producing significant transient changes (Fig. S18J,R).

Overall, the response to increased DeSI is dominated by enhanced deSUMOylation. The stressed steady state is determined by the deSUMOylation rates, while the transient response is remarkably robust to variation in the remaining parameters. Only changes in $K_2$ and the initial SUMO-to-target ratio substantially modify the transient dynamics.

**Summary.** Overall, the step-change analysis demonstrates that transient SUMOylation dynamics depend strongly on which component of the SUMO cycle is perturbed. Increased preSUMO promotes sustained SUMOylation, elevated ULP produces an initial decrease followed by recovery through enhanced SUMO maturation, increased E2 generates a purely transient increase without altering the stressed steady state, and increased DeSI drives a monotonic reduction in SUMOylation. Across all perturbations, the intrinsic timescale determines whether these responses appear as broad sustained changes or sharp transient peaks, with faster systems exhibiting earlier and more pronounced dynamics.

Parameter variation shows that these qualitative responses are robust despite uncertainty in the model parameters. Most perturbations primarily affect the timing and magnitude of the transient response rather than its overall form. The dynamics are consistently most sensitive to the intermediate SUMOylation reaction rates $K_2$–$K_5$ and the initial SUMO-to-target ratio $R_{s,T}$, identifying these as the principal regulators of transient SUMOylation. Together, these results suggest that transient behavior is an intrinsic feature of the SUMO cycle and provide a mechanistic basis for interpreting the time-resolved experimental data presented in the following section.

### 4. Simulation of SUMOylated target dynamics given experimental data

The steady-state analysis presented in section 2 describes how changes in individual SUMO pathway components influence the long-term level of SUMOylated target protein. However, abiotic stress induces transient and tissue-specific changes in the abundance of SUMO pathway components, which may interact with the intrinsic kinetics of the SUMO cycle to produce dynamic responses that cannot be captured by a steady-state analysis alone. In particular, transient changes in SUMO, ULP, E2 and DeSI may propagate through the conjugation and deconjugation machinery, generating peaks, troughs, or sustained changes in SUMOylated target levels.

To investigate these effects, we incorporated experimentally measured time courses of SUMO, ULP, E2, and DeSI into a reduced model. This allowed us to use the observed dynamics as time-dependent inputs while retaining the mechanistic ODE description of the remaining SUMO pathway components. We first describe how the experimental measurements were converted into fold-change trajectories and how their associated variability was quantified. We then describe the formulation of the dynamic model and use it to investigate how experimentally observed perturbations propagate through the SUMO cycle and influence SUMOylated target dynamics using the findings from section 3.

**4.1. Processing the experimental time-course data.** The raw data consist of fluorescence-per-area measurements for representative isoforms of SUMO, ULP, E2, and DeSI in different root tissues under control conditions and following salt or osmotic stress. Salt-stress measurements were collected at baseline and after 0.5, 1, 2, and 3 h, whereas osmotic-stress measurements were collected at baseline and after 1, 2, 3, and 5 h. For each isoform, tissue, stress condition, and time point, multiple biological replicates were obtained (see Fig. 3).

To compare measurements across experiments and time points, fluorescence measurements were expressed as fold changes relative to the corresponding control measurements. Rather than calculating fold changes from the means of the stressed and control measurements, we propagated the variability in the raw measurements by considering all possible pairings of stressed and control measurements. This provides a distribution of fold-change values from which both the mean response and its associated variability can be estimated.

For the salt stress experiments, a single baseline control dataset was paired with each stressed time-point dataset. For the osmotic stress experiments, each stressed time-point dataset was paired with its corresponding time-matched control dataset. Specifically, for a given isoform, tissue, stress condition, and time point, let $N$ denote the corresponding control measurements and $M$ the stressed measurements. We define the fold-change dataset as

$$J = \left\{ \frac{m}{n} \ : \ m \in M, \ n \in N \right\}. \tag{36}$$

The mean and standard deviation of $J$ were then used to represent the experimentally observed fold change and its associated uncertainty for each isoform, tissue, stress condition, and time point.

**4.2. Formulation of a reduced dynamic model using experimental data.** To incorporate the experimentally observed time courses into the model, we modify system Eq. (9) by prescribing the dynamics of SUMO, ULP, E2, and DeSI. For each species $R \in \{s, u, 2, D\}$, we construct a continuous input function $I_R(t)$ by linearly interpolating the experimentally determined fold changes shown in Fig. 3. Here, $I_R(t)$ represents the fold change of species $R$ relative to its unstressed steady-state level at dimensional time $t$. The experimentally prescribed species are given by

$$[\mathrm{SUMO}]^*(t) = I_s(t), \tag{37a}$$
$$[\mathrm{ULP}]^*(t) = I_u(t), \tag{37b}$$
$$[\mathrm{E2}]^*(t) = I_2(t), \tag{37c}$$
$$[\mathrm{DeSI}]^*(t) = I_D(t). \tag{37d}$$

As the interpolated experimental inputs are functions of dimensional time, we redimensionalize time in the governing non-dimensional equations by rescaling each ODE using $\beta_s$ (see Eq. (7)). Furthermore, as we are no longer prescribing E1 or E3 fold changes we set $C_1 = 1$ and $C_3 = 1$. Prescribing SUMO directly also removes the dependence of the remaining system on the preSUMO equation. Thus, with these modifications, from Eq. (9) we obtain the reduced dynamic system:

$$\frac{\mathrm{d}}{\mathrm{d}t}[\mathrm{E1}]^* = \beta_s \left[ \gamma_1 (C_1 - [\mathrm{E1}]^*) + K_2 R_{s,1} ([\mathrm{E2}]^* [\mathrm{E1} \cdot \mathrm{SUMO}]^* - [\mathrm{E1}]^* [\mathrm{SUMO}]^*) \right], \tag{38a}$$

$$\frac{\mathrm{d}}{\mathrm{d}t}[\mathrm{E1} \cdot \mathrm{SUMO}]^* = \beta_s K_3 ([\mathrm{E1}]^* [\mathrm{SUMO}]^* - [\mathrm{E2}]^* [\mathrm{E1} \cdot \mathrm{SUMO}]^*), \tag{38b}$$

$$\begin{aligned}\frac{\mathrm{d}}{\mathrm{d}t}[\mathrm{E2} \cdot \mathrm{SUMO}]^* = {} & \beta_s \frac{K_2 K_4 R_{s,T}}{K_2 R_{s,T} - K_5} ([\mathrm{E2}]^* [\mathrm{E1} \cdot \mathrm{SUMO}]^* - [\mathrm{target}]^* [\mathrm{E2} \cdot \mathrm{SUMO}]^*) \\ & + \beta_s \frac{K_4 K_5}{K_2 R_{s,T} - K_5} ([\mathrm{target}]^* [\mathrm{E2} \cdot \mathrm{SUMO}]^* - [\mathrm{E2} \cdot \mathrm{SUMO}]^* [\mathrm{E3} \cdot \mathrm{target}]^*),\end{aligned} \tag{38c}$$

$$\frac{\mathrm{d}}{\mathrm{d}t}[\mathrm{target}]^* = \beta_s \Bigg[ \gamma_T(1-[\mathrm{target}]^*) - K_2 R_{s,T}[\mathrm{target}]^*[\mathrm{E2}\cdot\mathrm{SUMO}]^* + K_5([\mathrm{target}]^*[\mathrm{E2}\cdot\mathrm{SUMO}]^* - [\mathrm{E3}]^*[\mathrm{target}]^*) + \frac{K_2K_7}{K_7+K_8}[\mathrm{target}\cdot\mathrm{SUMO}]^*[\mathrm{ULP}]^* + \frac{K_2K_8}{K_7+K_8}[\mathrm{target}\cdot\mathrm{SUMO}]^*[\mathrm{DeSI}]^* \Bigg], \quad [38d]$$

$$\frac{\mathrm{d}}{\mathrm{d}t}[\mathrm{E3}]^* = \beta_s\left[\gamma_3(1-[\mathrm{E3}]^*) + K_5 R_{T,3}([\mathrm{E2}\cdot\mathrm{SUMO}]^*[\mathrm{E3}\cdot\mathrm{target}]^* - [\mathrm{E3}]^*[\mathrm{target}]^*)\right], \quad [38e]$$

$$\frac{\mathrm{d}}{\mathrm{d}t}[\mathrm{E3}\cdot\mathrm{target}]^* = \beta_s\frac{K_6(K_2R_{s,T}-K_5)}{K_4}([\mathrm{E3}]^*[\mathrm{target}]^* - [\mathrm{E2}\cdot\mathrm{SUMO}]^*[\mathrm{E3}\cdot\mathrm{target}]^*), \quad [38f]$$

$$\frac{\mathrm{d}}{\mathrm{d}t}[\mathrm{target}\cdot\mathrm{SUMO}]^* = \beta_s\Bigg[\frac{(K_2R_{s,T}-K_5)(K_7+K_8)}{K_2R_{s,T}}[\mathrm{target}]^*[\mathrm{E2}\cdot\mathrm{SUMO}]^* + \frac{K_5(K_7+K_8)}{K_2R_{s,T}}[\mathrm{E2}\cdot\mathrm{SUMO}]^*[\mathrm{E3}\cdot\mathrm{target}]^* - K_7[\mathrm{target}\cdot\mathrm{SUMO}]^*[\mathrm{ULP}]^* - K_8[\mathrm{target}\cdot\mathrm{SUMO}]^*[\mathrm{DeSI}]^*\Bigg], \quad [38g]$$

where SUMO, ULP, E2, and DeSI are prescribed by Eq. (37). The system is initialized at the unstressed steady state, such that

$$[u]^*(0) = 1 \quad \text{for all} \quad u \in \{\mathrm{E1}, \mathrm{E1}\cdot\mathrm{SUMO}, \mathrm{E2}\cdot\mathrm{SUMO}, \mathrm{target}, \mathrm{E3}, \mathrm{E3}\cdot\mathrm{target}, \mathrm{target}\cdot\mathrm{SUMO}\},$$

with $I_R(0) = 1$ for $R \in \{s, u, 2, D\}$. The system is then solved numerically using `solve_ivp` from `SciPy.integrate` (6).

Prescribing the dynamics of SUMO, ULP, E2, and DeSI removes the dependence of the system on the parameters $\gamma_p$, $\gamma_u$, $\gamma_2$, $\gamma_D$, $K_1$, and $R_{T,2}$. The remaining parameter set is therefore

$$\chi = \{\gamma_1, \gamma_3, \gamma_T, \beta_s, K_2, K_3, K_4, K_5, K_6, K_7, K_8, R_{s,1}, R_{s,T}, R_{T,3}\}.$$

As in section 3, we define the baseline parameter set $\chi_g$ by setting $R_{s,T} = 2.5$, $\beta_s = g$, and all remaining parameters equal to one. We consider $\beta_s = 1$ and $\beta_s = 3$, corresponding to intrinsic timescales of approximately 1 h and 20 min, respectively.

**4.3. Analyzing model dynamics given experimental data.** Having incorporated the experimental time-course data into the reduced model, we next examine how the observed changes in SUMO, ULP, E2, and DeSI propagate through the remainder of the SUMO cycle. We first quantify the effect of experimental uncertainty on the predicted SUMOylated target dynamics, before examining the coordinated response of the remaining pathway components. Finally, we assess how variation in the model parameters influences the predicted temporal response.

***4.3.1. Computing SUMOylated target dynamics given experimental uncertainty.*** Fig. 3 provides the experimentally derived means and standard deviations of fold changes for four tissue–stress combinations: epidermis under salt stress, stele under salt stress, epidermis under osmotic stress, and cortex under osmotic stress. These values were obtained using the procedure described in section 4.1.

To account for the uncertainty in the experimental measurements, the fold change at each time point was represented by a normal distribution with mean and standard deviation given by the corresponding experimental values. One thousand samples were drawn from these distributions; non-positive samples were resampled until a positive value was obtained. The resulting samples were propagated through the dynamic model. Each sampled dataset therefore provides a distinct realization of the experimentally observed time courses, allowing the uncertainty in the experimental measurements to be propagated through the SUMO cycle.

For each sampled time course, the system was solved using `SciPy.integrate`. The resulting trajectories for SUMOylated target were then combined to obtain the median, interquartile range, and 95% interval at each time point. This procedure allows the predicted dynamics to reflect both the temporal structure of the experimental data and its associated variability. The resulting predictions are shown in Fig. 5A–H.

***4.3.2. Predicted dynamics of the remaining SUMO cycle species.*** Using the mean experimentally derived fold-change trajectories for SUMO, ULP, E2, and DeSI (Fig. 3) as inputs, the model predicts coordinated changes throughout the remainder of the SUMO pathway (Figs. 5I–O, S19, S24, S27). Across the tissue–stress conditions considered, E1 · SUMO decreases while E2 · SUMO increases, reflecting the redistribution of SUMO from the E1-bound to the E2-bound state. This is accompanied by an increase in free E1 and a decrease in E3 · target, as the latter is converted to SUMOylated target. In contrast, changes in free target and E3 are comparatively small. The magnitude of these responses broadly follows the magnitude of the prescribed experimental perturbations, with the largest changes predicted for salt stress in the stele and the smallest for osmotic stress in the cortex.

The intrinsic timescale of the SUMO cycle, controlled by $\beta_s$, determines how these experimentally prescribed changes interact as they propagate through the pathway. Increasing $\beta_s$ does not simply accelerate the same response; it can qualitatively alter the resulting dynamics. At the faster intrinsic timescale, perturbations propagate more rapidly through the conjugation pathway, allowing transient features in the experimentally prescribed inputs to generate pronounced peaks and troughs in downstream species. In contrast, the slower intrinsic timescale smooths these interactions, producing broader and more sustained responses.

Together, these results demonstrate that the experimentally observed dynamics of individual SUMO pathway components can generate qualitatively distinct downstream SUMOylation dynamics. The tissue- and stress-specific inputs determine the magnitude and temporal structure of the perturbation, while the intrinsic timescale of the SUMO cycle determines how these signals are integrated and propagated through the pathway.

***4.3.3. Sensitivity of SUMOylated target dynamics.*** To assess the robustness of the predicted SUMOylation dynamics, each remaining model parameter was independently doubled and halved about the baseline parameter sets $\chi_1$ and $\chi_3$. The resulting changes in SUMOylated target dynamics were examined across the four tissue–stress conditions (Figs. S20, S21, S22, S23, S25, S26, S28, S29).

Across all tissue–stress conditions and both baseline parameter sets, variations in $\gamma_3$, $\gamma_T$, and $R_{T,3}$ have little to no effect on the predicted SUMOylated target dynamics. This indicates that, within the parameter ranges considered, variation in E3 production, target production, and E3 · target interaction strength has limited influence on the downstream response.

For the slower intrinsic timescale, most parameter perturbations primarily alter the magnitude of the SUMOylated target response, with $K_2$ and $R_{s,T}$ being notable exceptions. In contrast, at the faster intrinsic timescale, variations in the intermediate conjugation parameters $K_2$–$K_6$ more frequently produce qualitative changes in the predicted dynamics, including changes in the amplitude, timing, and prominence of transient peaks and troughs. The extent and direction of these changes depend on the prescribed experimental inputs and their relative magnitudes. Thus, parameter variation modifies how the tissue- and stress-specific input trajectories are translated into downstream SUMOylation dynamics.

**Summary.** The dynamic model demonstrates how experimentally observed changes in SUMO, ULP, E2, and DeSI can propagate through the SUMO cycle to generate tissue- and stress-specific SUMOylated target dynamics. The experimental trajectories determine the magnitude and temporal structure of the perturbation, while the intrinsic timescale of the pathway determines how these signals interact and propagate. In particular, faster intrinsic dynamics can preserve and amplify transient features of the experimental inputs, resulting in qualitatively distinct peaks and troughs, whereas slower dynamics tend to produce smoother and more sustained responses. Parameter sensitivity analysis further indicates that the predicted dynamics are relatively insensitive to several parameters associated with E3 and target regulation, while parameters governing the intermediate conjugation reactions can strongly influence the qualitative behavior of the system, particularly at faster intrinsic timescales.

### Supplementary figures

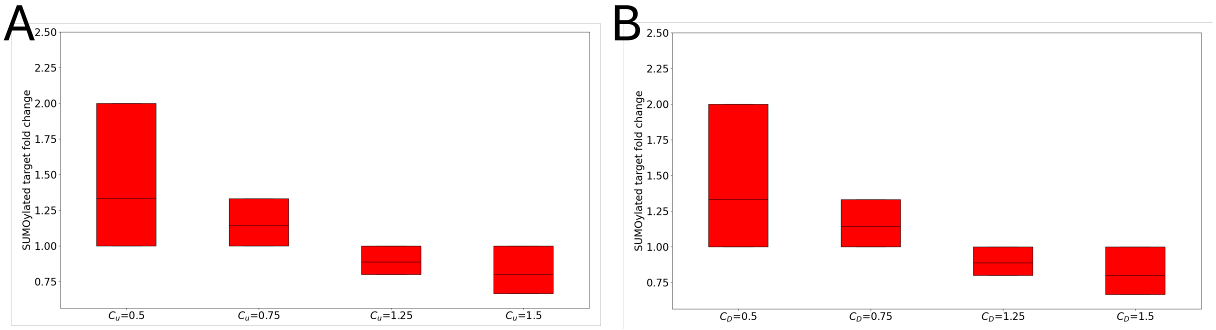


**Supplementary Figure S1. Effect of ULP or DeSI fold changes on SUMOylated target.** Predicted steady-state fold changes of SUMOylated target given variation only in (**A**) ULP, with $C_u \in \{0.5, 0.75, 1.25, 1.5\}$ and $C_s = C_1 = C_D = 1$, and (**B**) DeSI, with $C_D \in \{0.5, 0.75, 1.25, 1.5\}$ and $C_s = C_u = C_1 = 1$ (see $\mathrm{Eq.}\ (29)$). In all simulations, $K_7$ and $K_8$ were varied over the range $[1/100, 100]$ using 10,000 parameter sets sampled from log space as described in section 2.3. Results in (**A**) are consistent with the analytical bounds in $\mathrm{Eq.}\ (30)$.

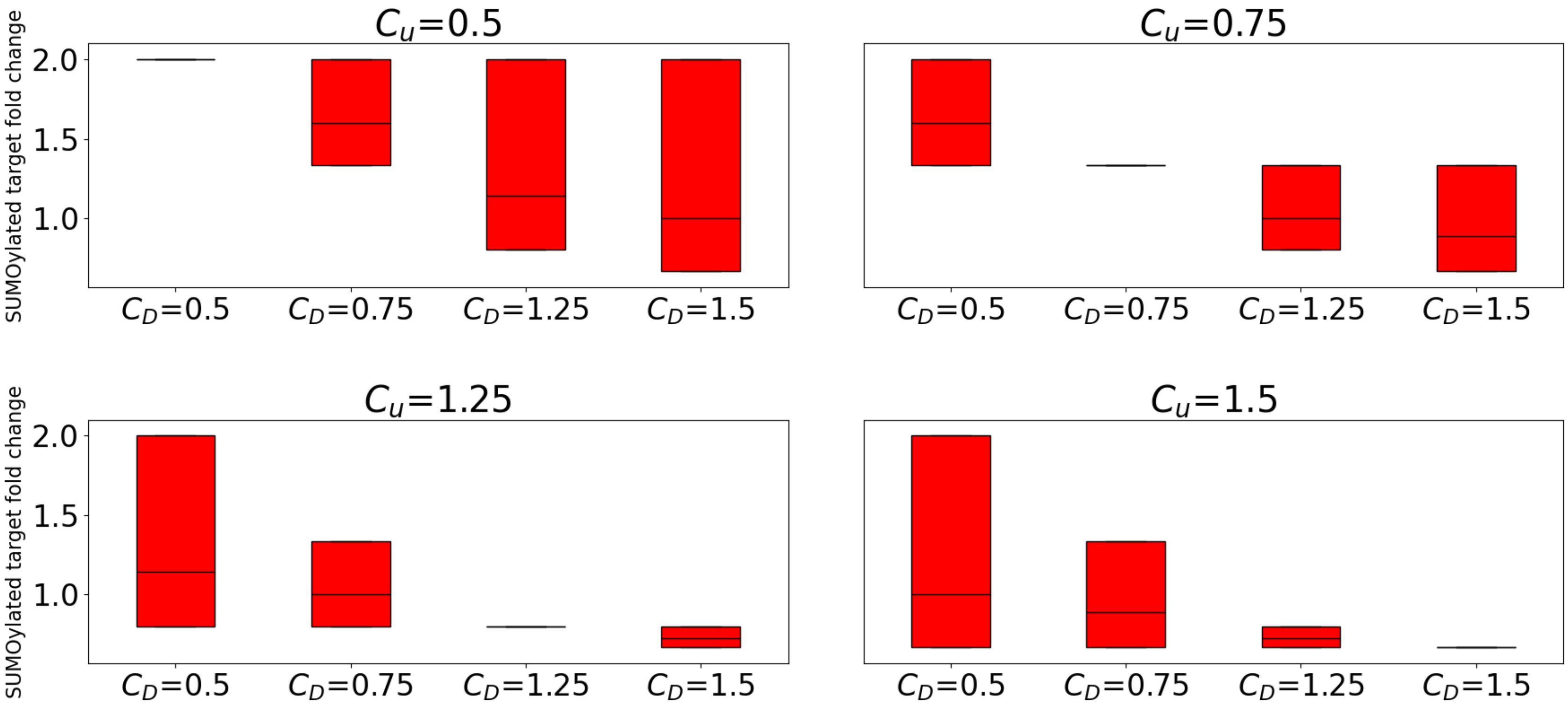


**Supplementary Figure S2. Combined effect of ULP and DeSI fold changes.** Predicted steady-state fold changes of SUMOylated target given variation in both ULP and DeSI (see $\mathrm{Eq.}\ (31)$) for $C_u, C_D \in \{0.5, 0.75, 1.25, 1.5\}$ and $C_s, C_1 = 1$. In all simulations, $K_7$ and $K_8$ were varied over the range $[1/100, 100]$ using 10,000 parameter sets sampled from log space as described in section 2.3. Results agree with the analytical bounds in $\mathrm{Eq.}\ (32)$.

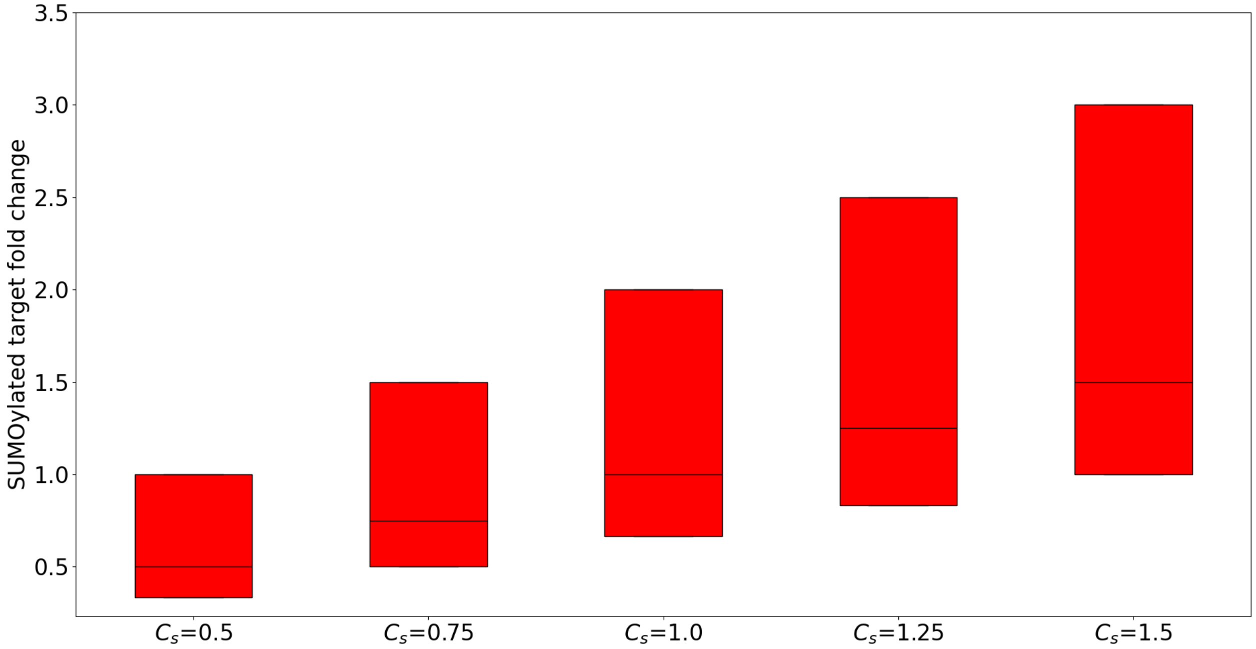


**Supplementary Figure S3. Effect of varying SUMO with fixed ULP and DeSI.** Predicted steady-state fold changes of SUMOylated target for $C_s \in \{0.5, 0.75, 1.25, 1.5\}$ with $C_u = 0.5$, $C_1 = 1$, and $C_D = 1.5$ (see $\mathrm{Eq.}\ (14)$). In all simulations, $K_7$ and $K_8$ were varied over the range $[1/100, 100]$ using 10,000 parameter sets sampled from log space as described in section 2.3. Results agree with $\mathrm{Eq.}\ (34)$. We note that varying the value of $C_1$ for the same values of $C_u$ and $C_D$ produces the same output.

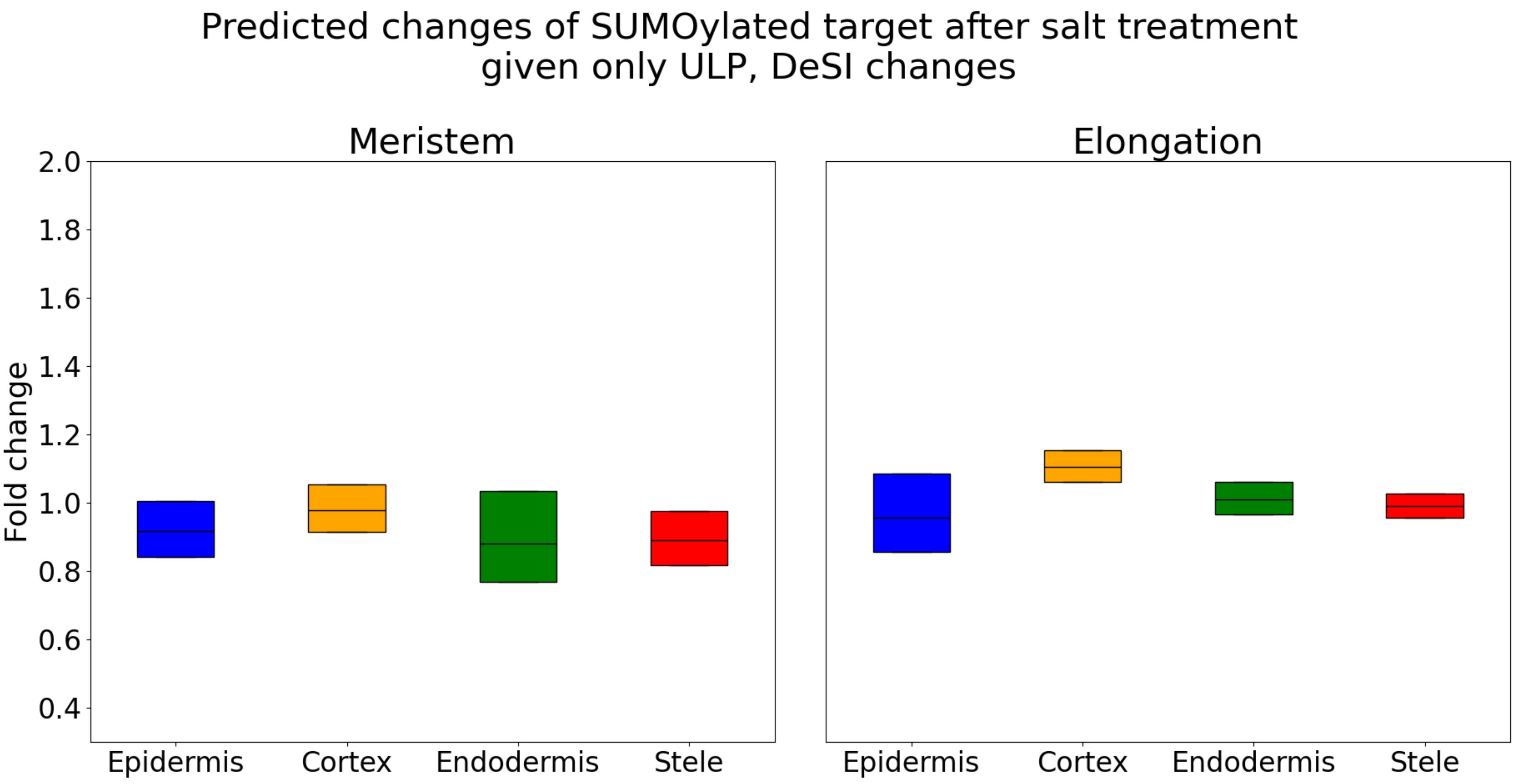


**Supplementary Figure S4. Predicted SUMOylated target steady-state fold changes after salt stress given only ULP and DeSI changes.** Model-predicted fold changes of SUMOylated target across the parameter space (see $\mathrm{Eq.}\ (14)$) using the amalgamated salt stress data (Fig. 2**B**) and considering only ULP and DeSI fold changes. In all simulations, $K_7$ and $K_8$ were varied over the range $[1/100, 100]$ using 10,000 parameter sets sampled from log space as described in section 2.3. Results agree with $\mathrm{Eq.}\ (34)$.

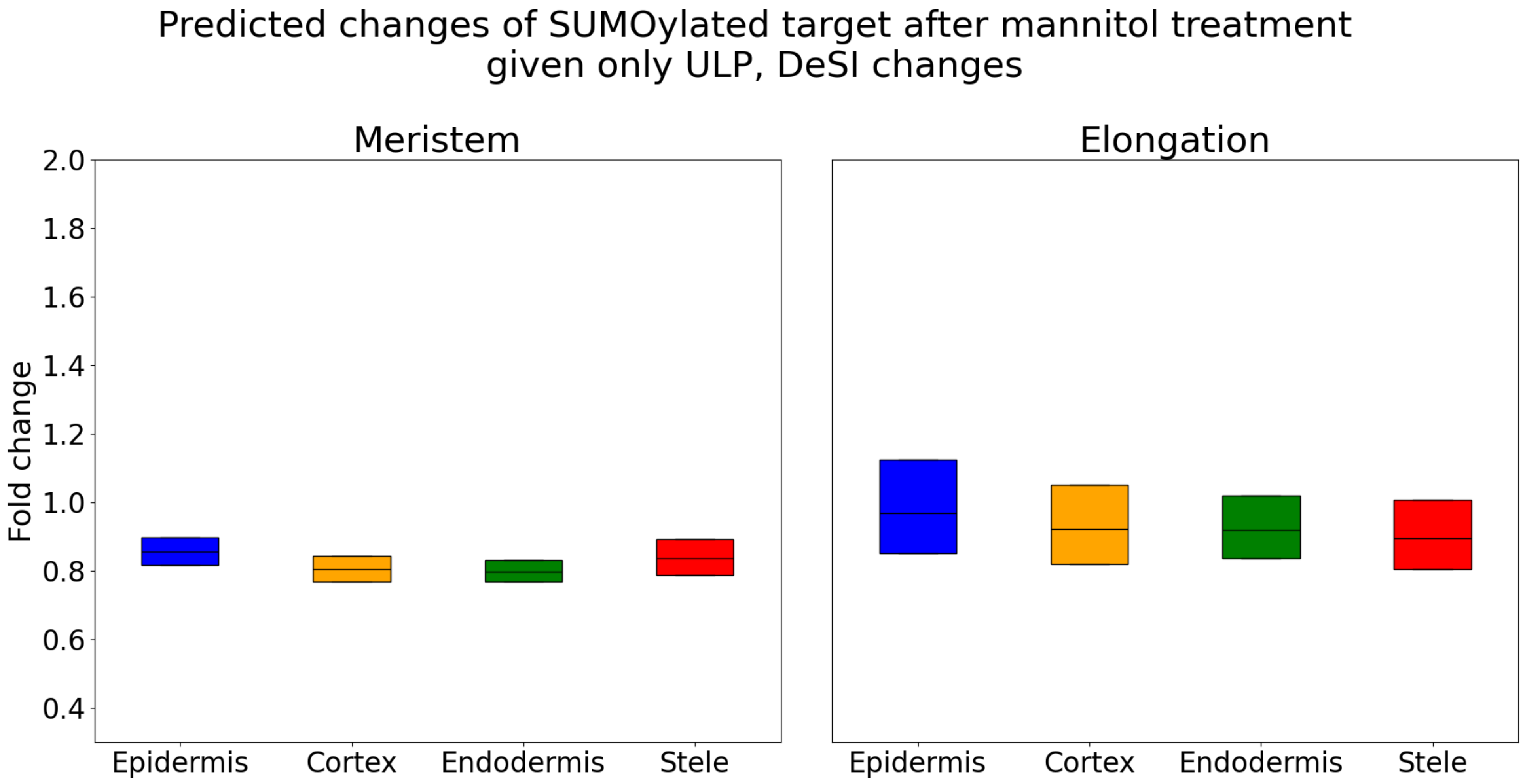


**Supplementary Figure S5. Predicted steady-state fold changes after osmotic stress given only ULP and DeSI changes.** Model-predicted fold changes of SUMOylated target across the parameter space (see $\mathrm{Eq.}\ (14)$) using the amalgamated mannitol stress data (Fig. 2**C**) and considering only ULP and DeSI fold changes. In all simulations, $K_7$ and $K_8$ were varied over the range $[1/100, 100]$ using 10,000 parameter sets sampled from log space as described in section 2.3. Results agree with $\mathrm{Eq.}\ (34)$.

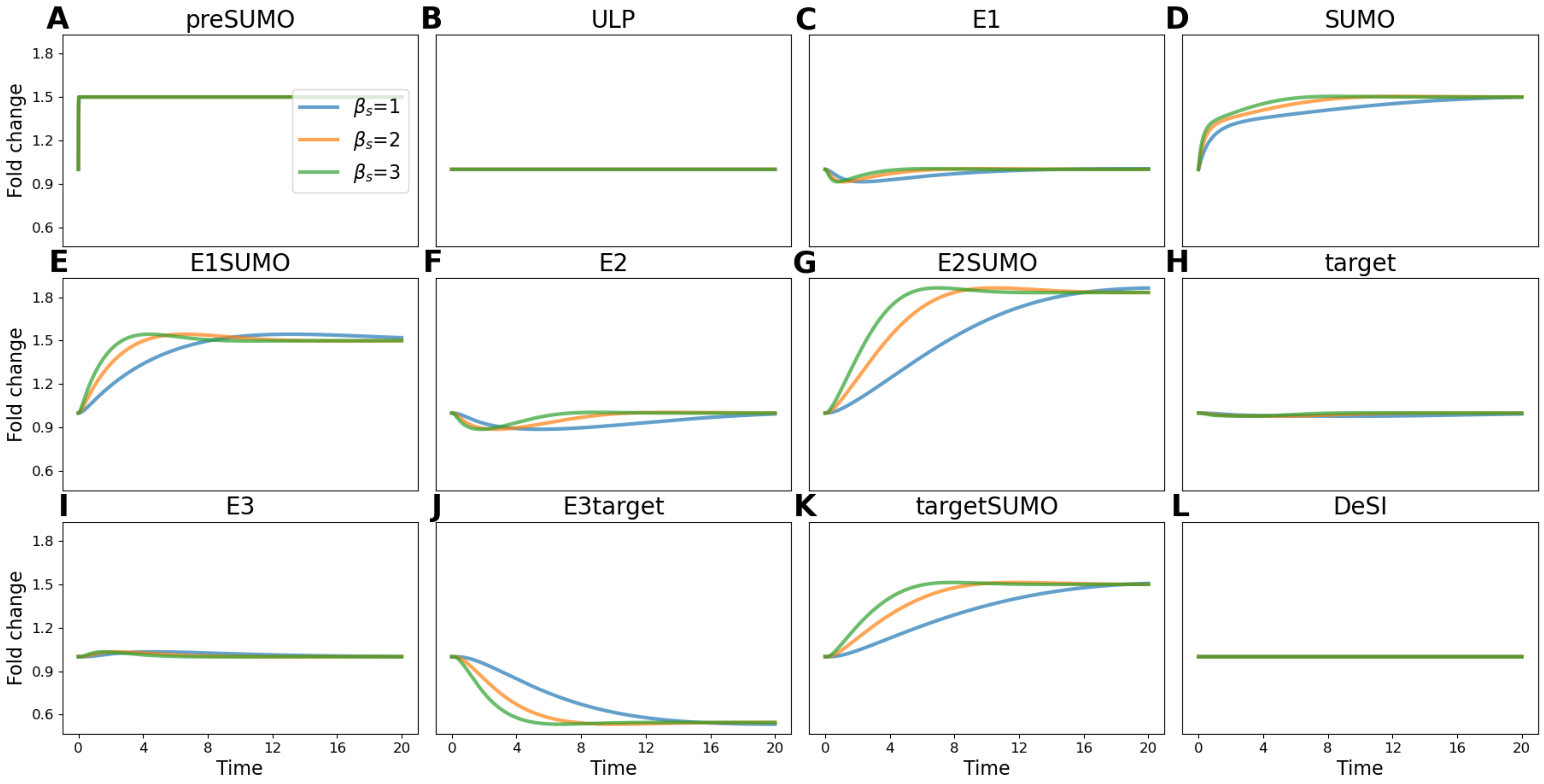


**Supplementary Figure S6. Predicted dynamics following a 50% increase in preSUMO.** System $\mathrm{Eq.}\ (9)$ is initialized at the unstressed steady state, after which $C_p$ is increased by 50% under parameter sets $\mu_{1,p}$, $\mu_{2,p}$ and $\mu_{3,p}$. Shown are the resulting model-predicted trajectories of all species.

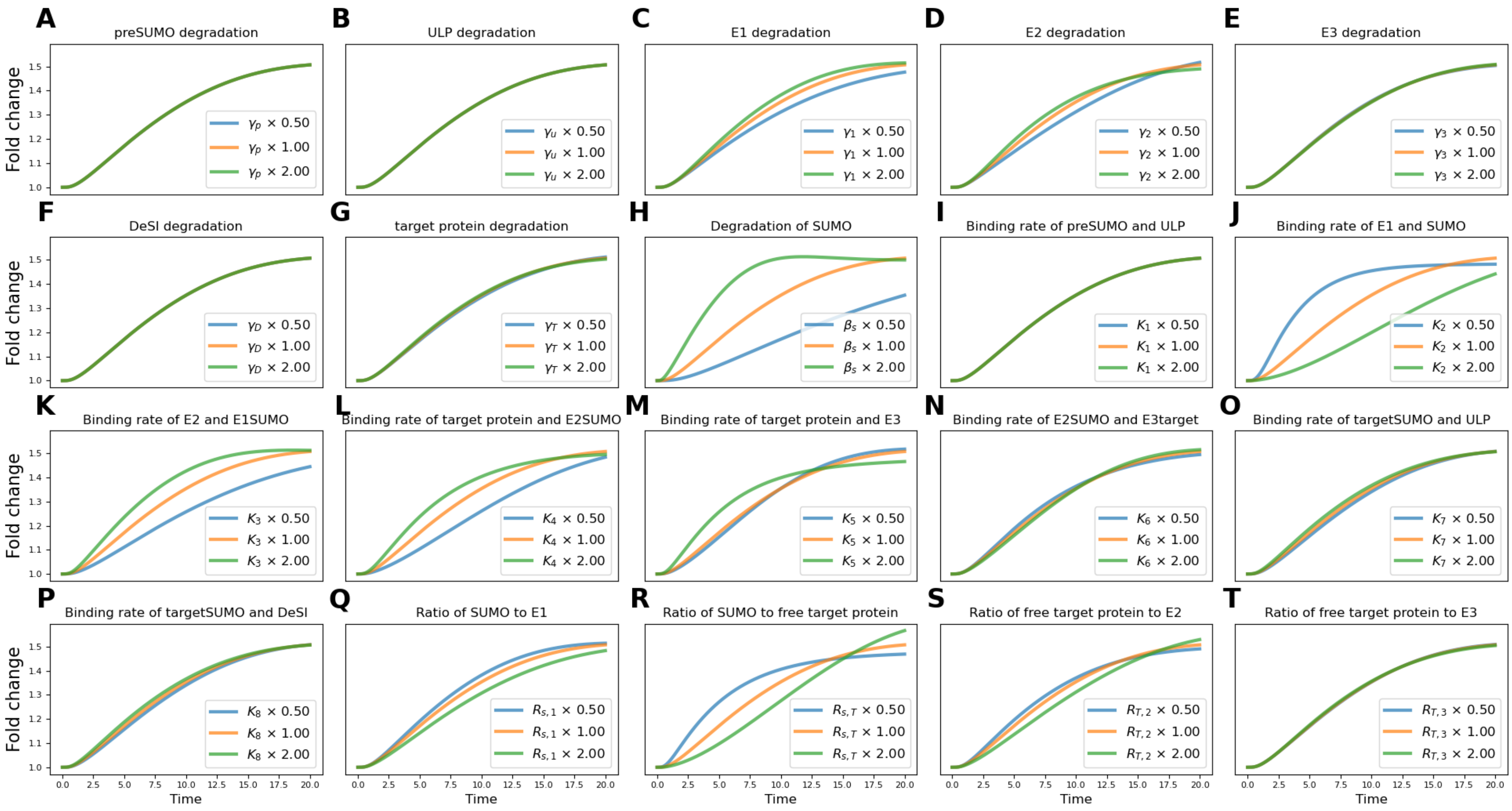


**Supplementary Figure S7. Sensitivity of SUMOylated target dynamics to parameter variation after a preSUMO increase given $\beta_s = 1$.** System Eq. (9) is initialized at the unstressed steady state and subjected to a 50% increase in preSUMO. Using the parameter set $\mu_{1,p}$ as a baseline, each non-dimensional parameter Eq. (8) and $\beta_s$ is varied independently by doubling and halving the parameter. Each subfigure corresponds to variation of the parameter indicated in its title.

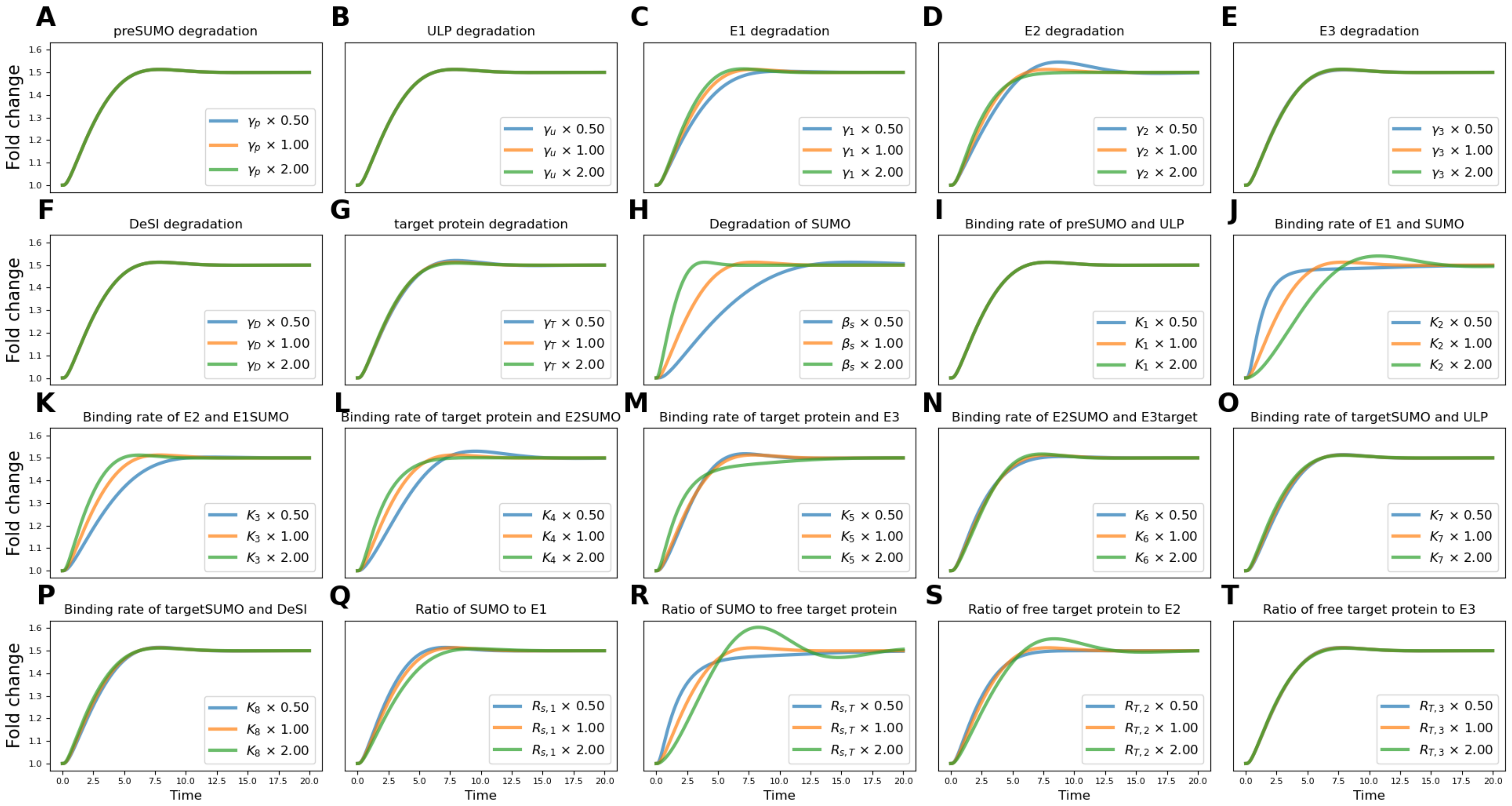


**Supplementary Figure S8. Sensitivity of SUMOylated target dynamics to parameter variation after a preSUMO increase given $\beta_s = 3$.** System $\mathrm{Eq.}~(9)$ is initialized at the unstressed steady state and subjected to a 50% increase in preSUMO. Starting from parameter set $\mu_{3,p}$, each non-dimensional parameter $\mathrm{Eq.}~(8)$ and $\beta_s$ is varied independently by doubling and halving the parameter. Each subfigure corresponds to variation of the parameter indicated in its title.

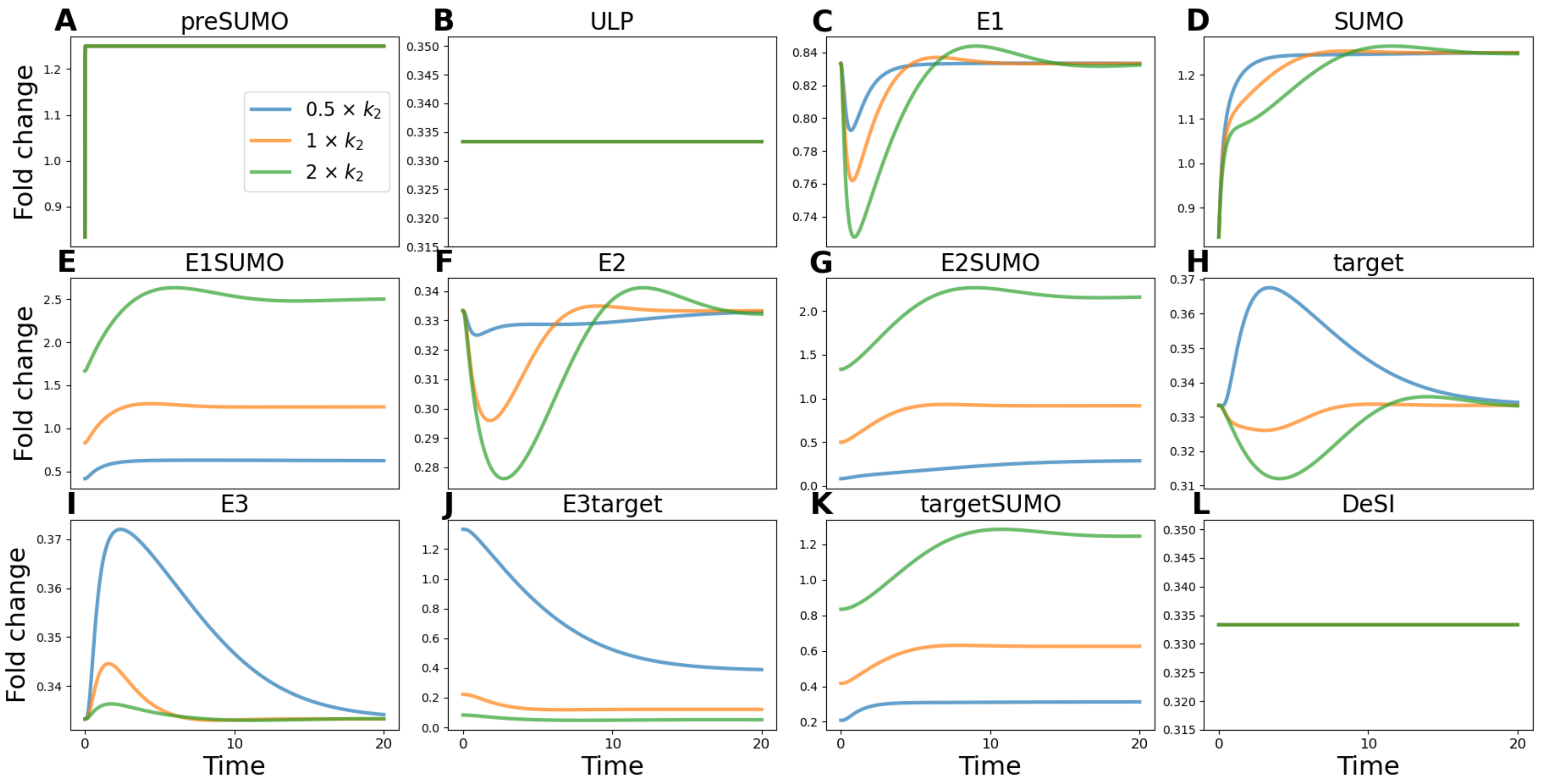


**Supplementary Figure S9. Predicted dimensional dynamics following a 50% increase in preSUMO and variation in $k_2$.** The dimensional system is initialized at the unstressed steady state, after which $C_p$ is increased by 50%. We redimensionalise using the set $\mu_{3,p}$ and setting $\alpha_u$, $\alpha_T$, and $\alpha_D$ equal to 1. We then plot the dimensional system under this set, alongside the set with $k_2$ halved and $k_2$ doubled. Shown are the resulting model-predicted trajectories of all species.

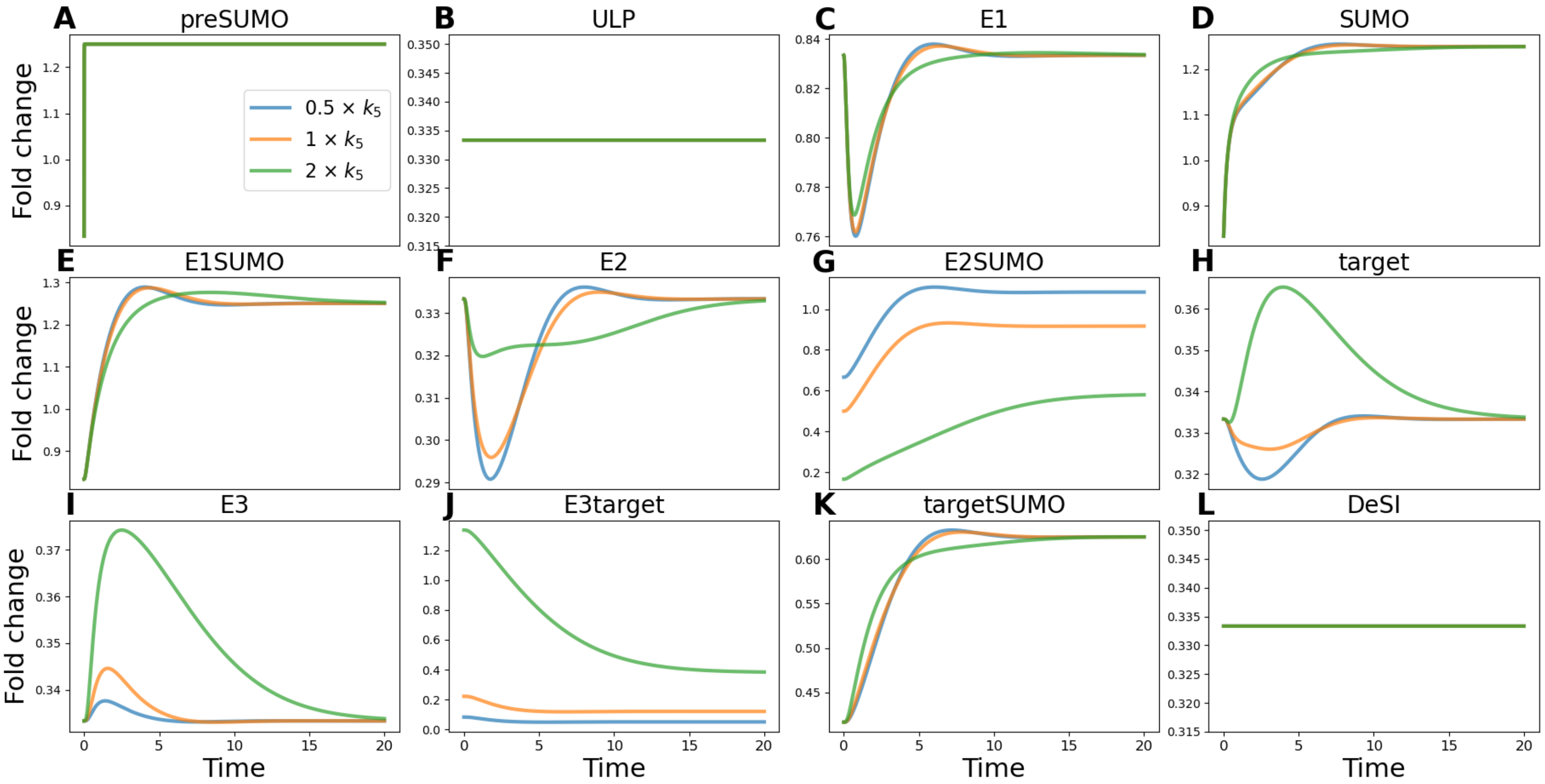


**Supplementary Figure S10. Predicted dimensional dynamics following a 50% increase in preSUMO and variation in $k_5$.** The dimensional system is initialized at the unstressed steady state, after which $C_p$ is increased by 50%. We redimensionalize using the set $\mu_{3,p}$ and setting $\alpha_u$, $\alpha_T$, and $\alpha_D$ equal to 1. We then plot the dimensional system under this set, alongside the set with $k_5$ halved and $k_5$ doubled. Shown are the resulting model-predicted trajectories of all species. See section 3.1 for details of redimensionalization.

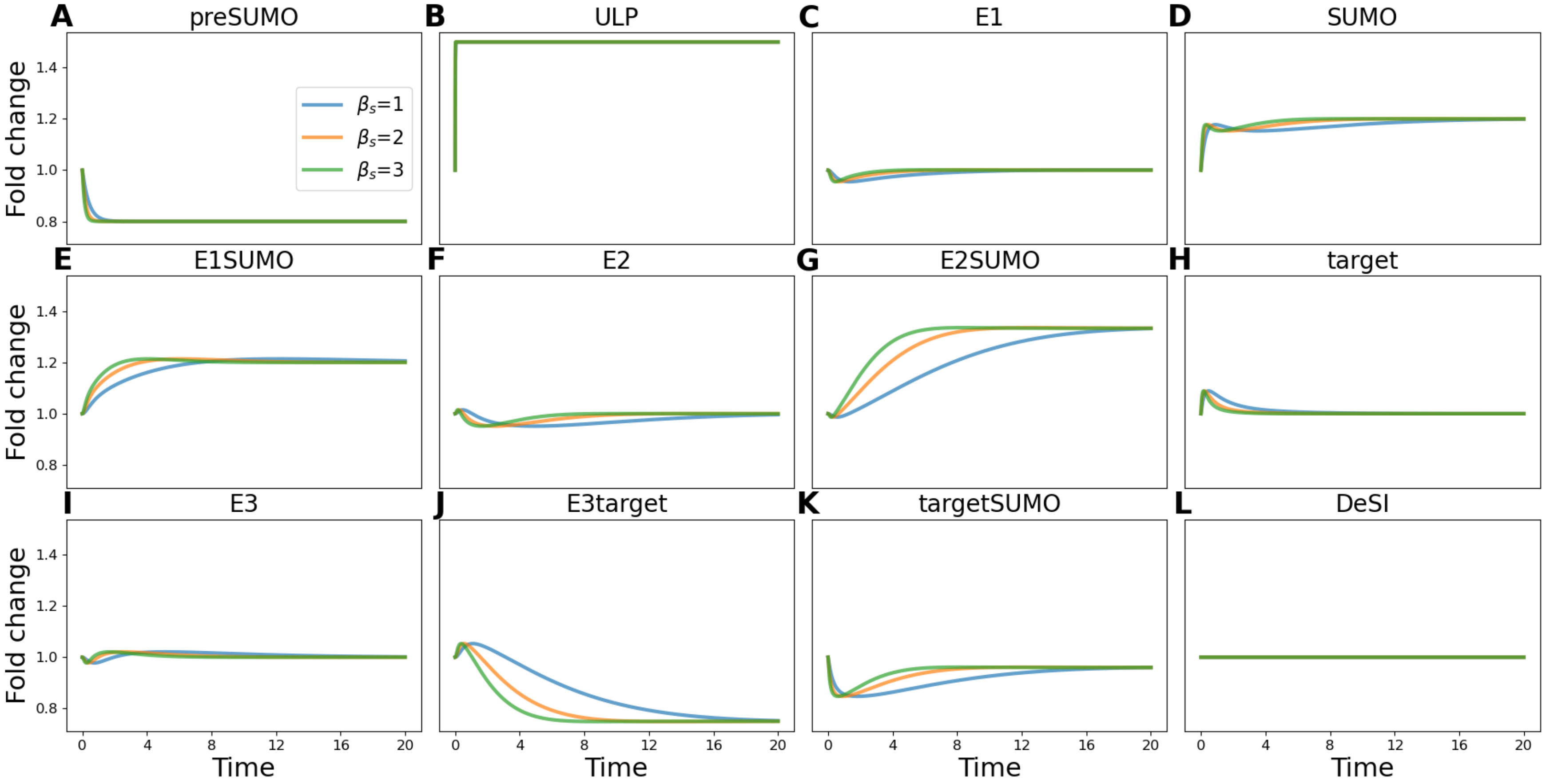


**Supplementary Figure S11. Predicted dynamics following a 50% increase in ULP.** System $\mathrm{Eq.}\ (9)$ is initialized at the unstressed steady state, after which $C_u$ is increased by 50% under parameter sets $\mu_{1,u}$, $\mu_{2,u}$ and $\mu_{3,u}$. Shown are the resulting model-predicted trajectories of all species.

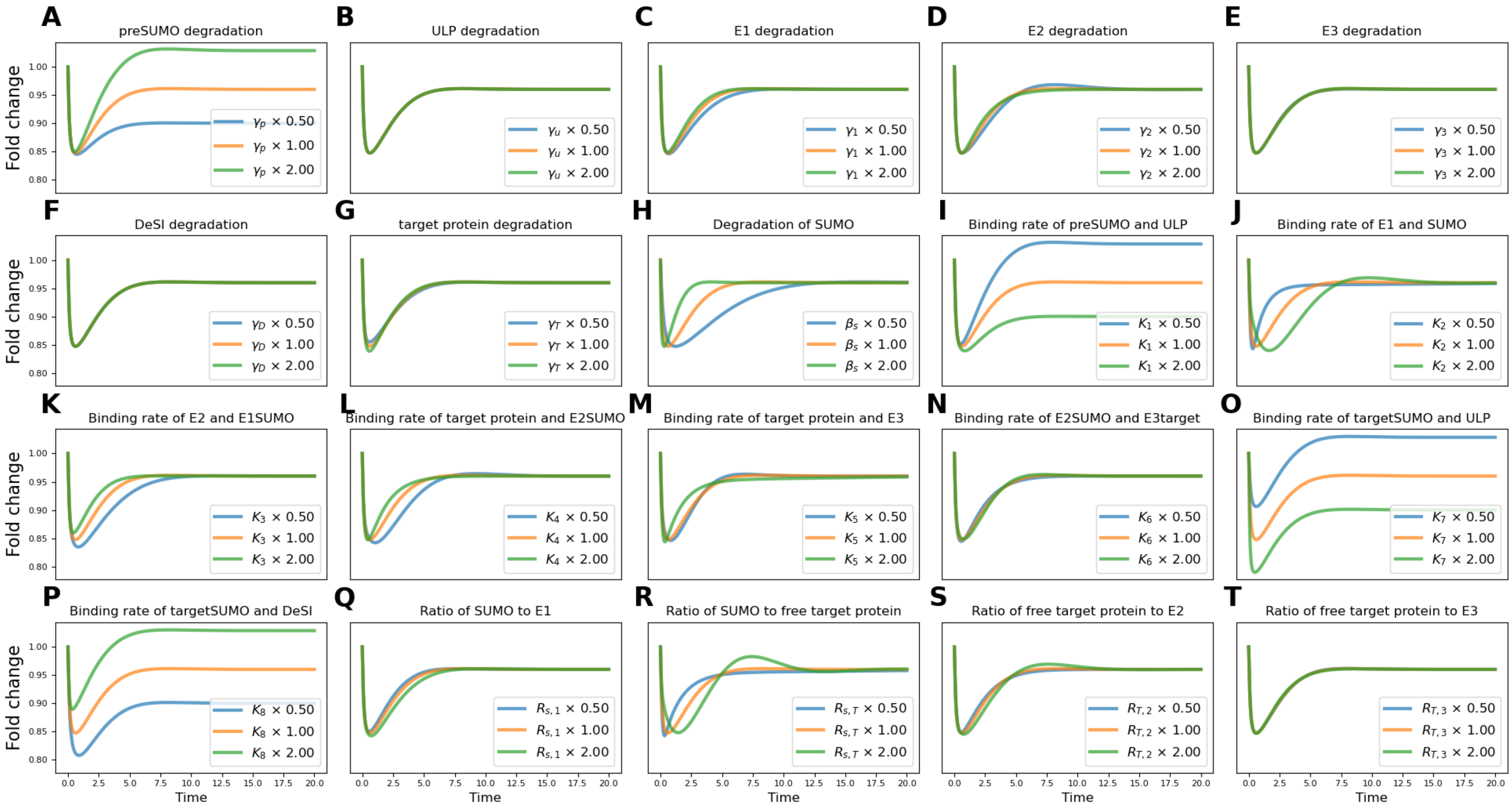


**Supplementary Figure S12. Sensitivity of SUMOylated target dynamics to parameter variation after a ULP increase given $\beta_s = 3$.** System Eq. (9) is initialized at the unstressed steady state and subjected to a 50% increase in ULP. Using the parameter set $\mu_{3,u}$ as a baseline, each non-dimensional parameter Eq. (8) and $\beta_s$ is varied independently by doubling and halving the parameter. Each subfigure corresponds to variation of the parameter indicated in its title.

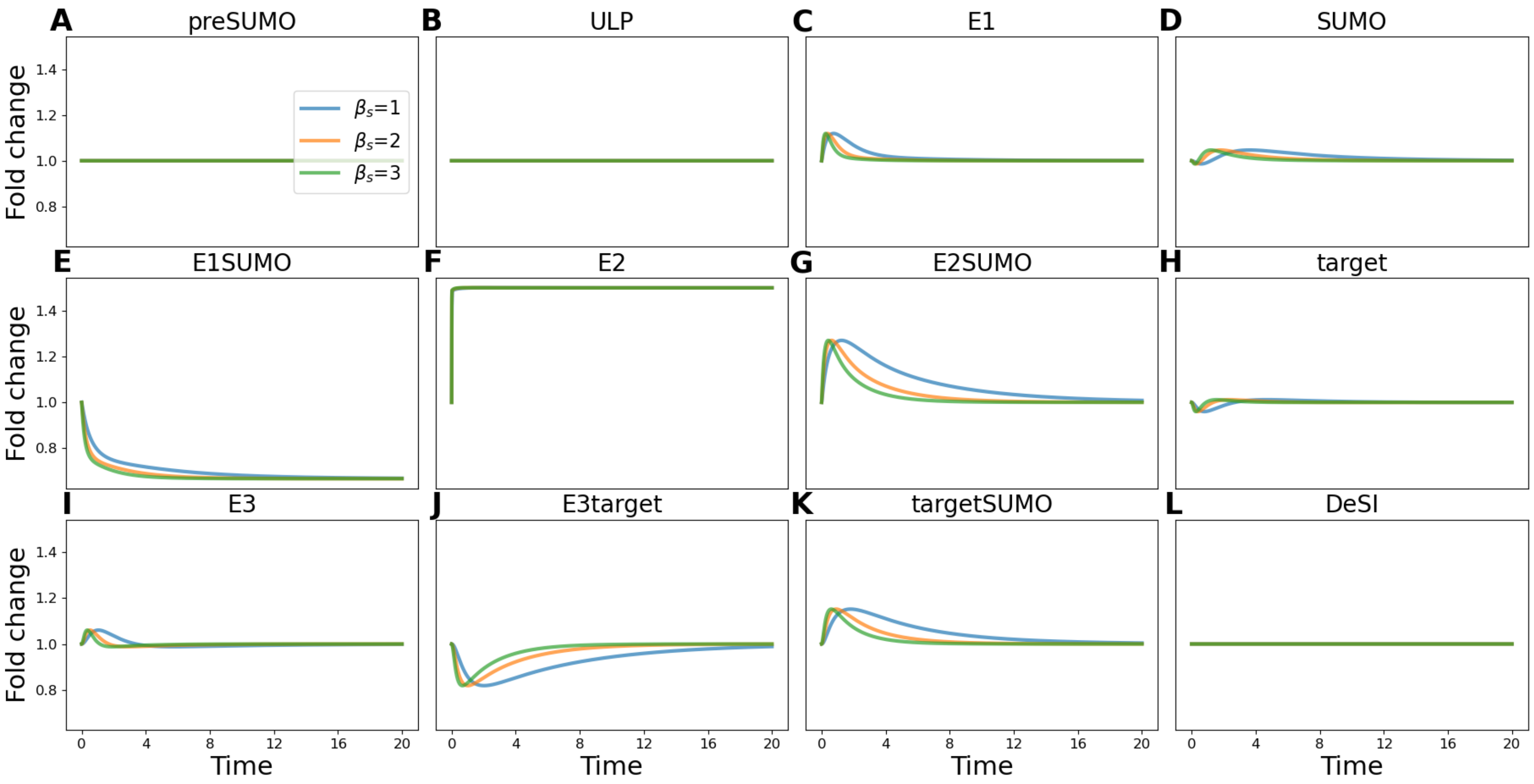


**Supplementary Figure S13. Predicted dynamics following a 50% increase in E2.** System Eq. (9) is initialized at the unstressed steady state, after which $C_2$ is increased by 50% under parameter set $\mu_{1,2}$, $\mu_{2,2}$ and $\mu_{3,2}$. Shown are the resulting model-predicted trajectories of all species.

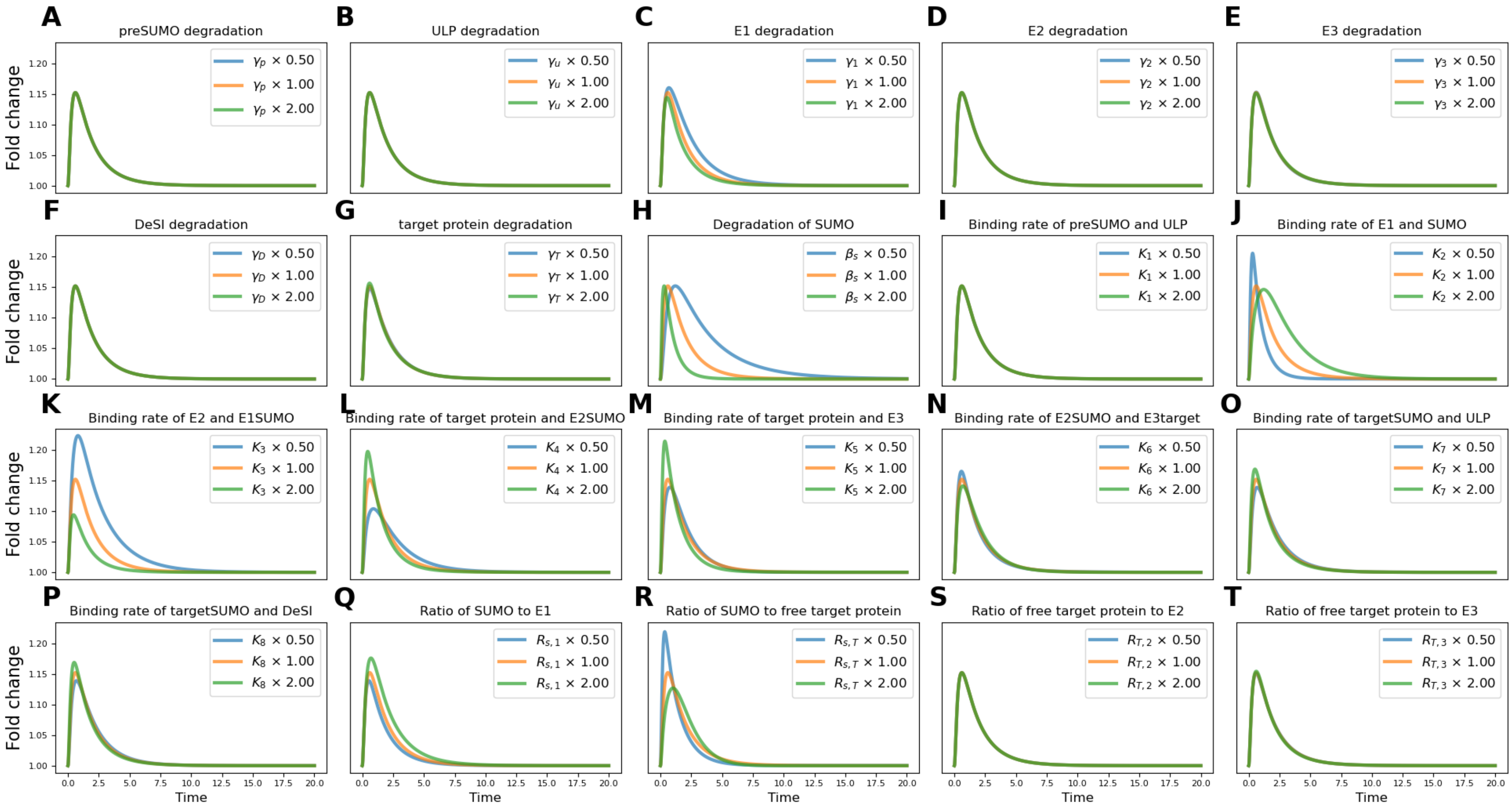


**Supplementary Figure S14. Sensitivity of SUMOylated target dynamics to parameter variation after an E2 increase given $\beta_s = 3$.** System $\mathrm{Eq.}\ (9)$ is initialized at the unstressed steady state and subjected to a 50% increase in E2. Using the parameter set $\mu_{3,2}$ as a baseline, each non-dimensional parameter $\mathrm{Eq.}\ (8)$ and $\beta_s$ is varied independently by doubling and halving the parameter. Each subfigure corresponds to variation of the parameter indicated in its title.

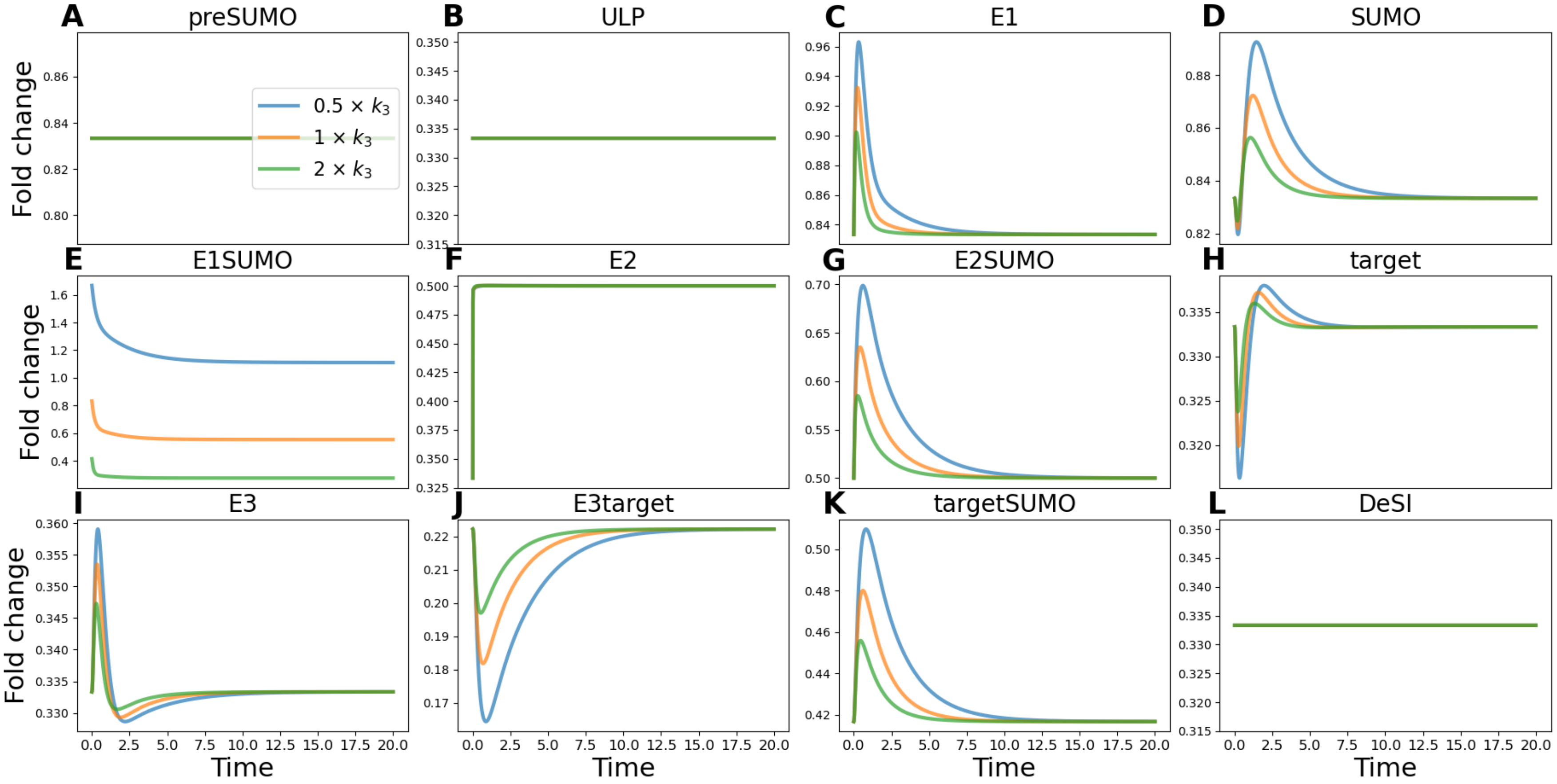


**Supplementary Figure S15. Predicted dimensional dynamics following a 50% increase in E2 and variation in $k_3$.** The dimensional system is initialized at the unstressed steady state, after which $C_2$ is increased by 50%. We redimensionalise using the set $\mu_{3,2}$ and setting $\alpha_u$, $\alpha_T$, and $\alpha_D$ equal to 1. We then plot the dimensional system under this set, alongside the set with $k_3$ halved and $k_3$ doubled. Shown are the resulting model-predicted trajectories of all species.

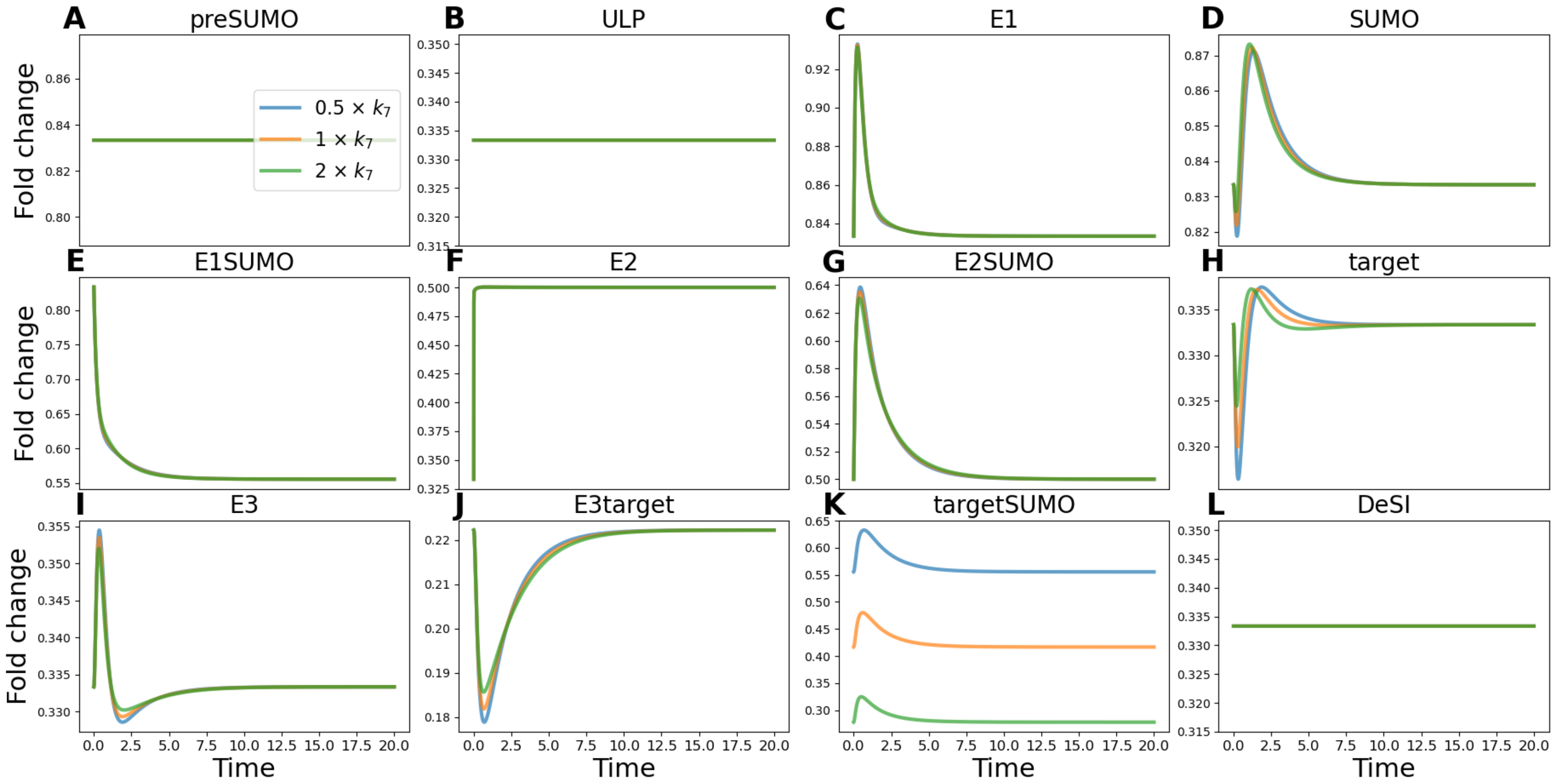


**Supplementary Figure S16. Predicted dimensional dynamics following a 50% increase in E2 and variation in $k_7$.** The dimensional system is initialized at the unstressed steady state, after which $C_2$ is increased by 50%. We redimensionalise using the set $\mu_{3,2}$ and setting $\alpha_u$, $\alpha_T$, and $\alpha_D$ equal to 1. We then plot the dimensional system under this set, alongside the set with $k_7$ halved and $k_7$ doubled. Shown are the resulting model-predicted trajectories of all species. Variation in $k_8$ produces identical qualitative behavior.

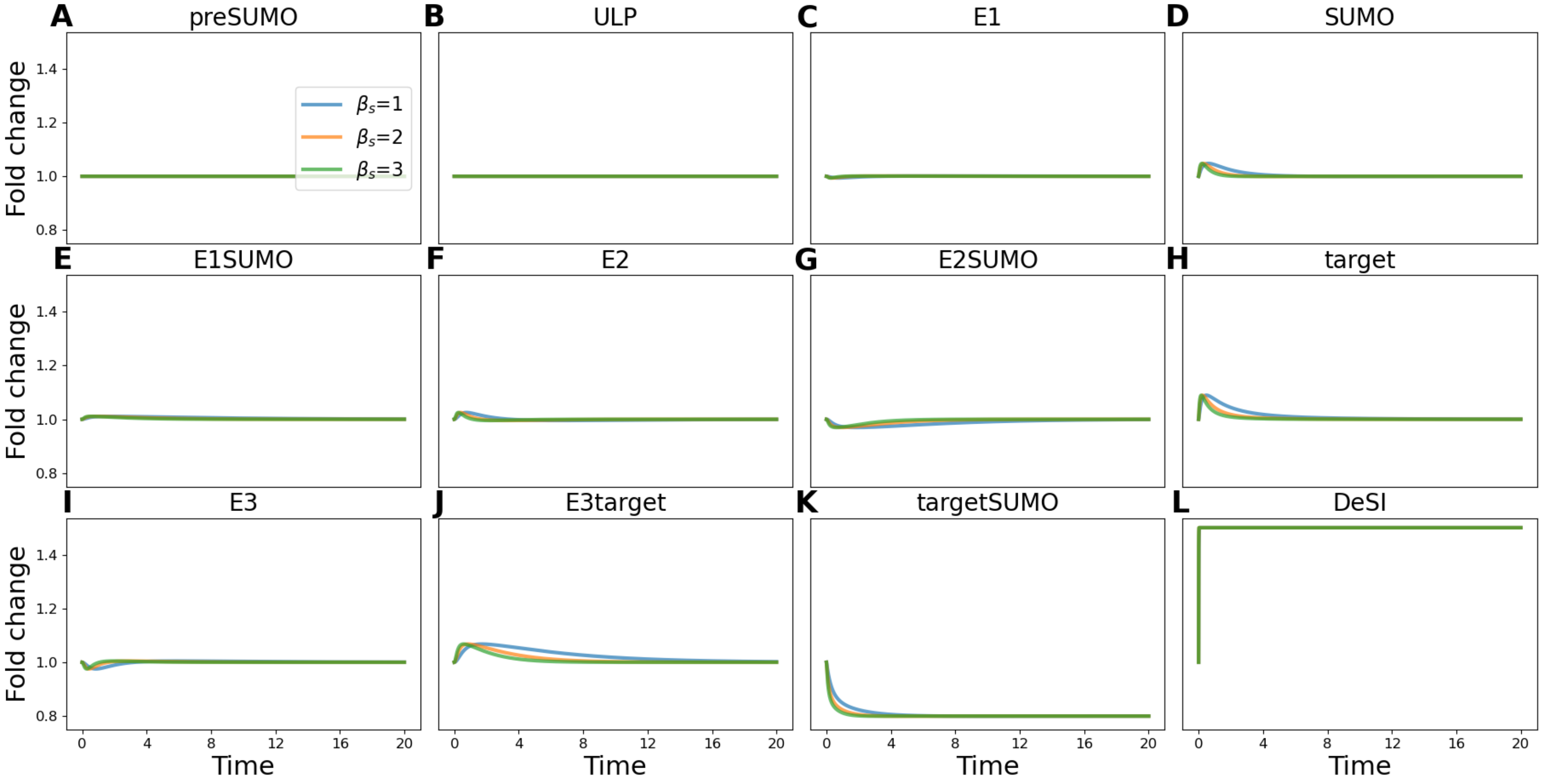


**Supplementary Figure S17. Predicted dynamics following a 50% increase in DeSI.** System $\mathrm{Eq.}~(9)$ is initialized at the unstressed steady state, after which $C_D$ is increased by 50% under parameter set $\mu_{1,D}$, $\mu_{2,D}$ and $\mu_{3,D}$. Shown are the resulting model-predicted trajectories of all species.

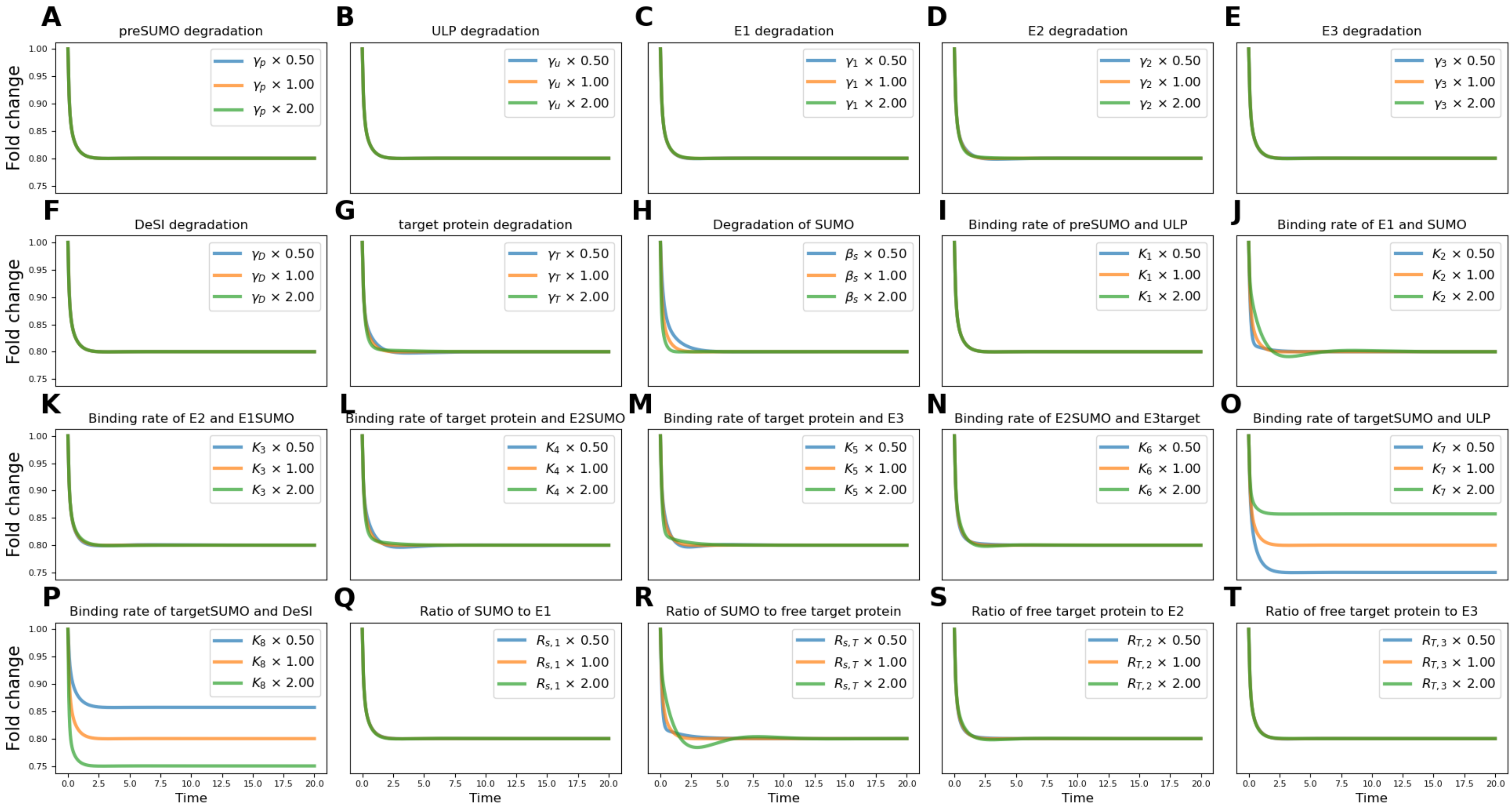


**Supplementary Figure S18. Sensitivity of SUMOylated target dynamics to parameter variation after a DeSI increase given $\beta_s = 3$.** System $\mathrm{Eq.}~(9)$ is initialized at the unstressed steady state and subjected to a 50% increase in DeSI. Using the parameter set $\mu_{3,D}$ as a baseline, each non-dimensional parameter $\mathrm{Eq.}~(8)$ and $\beta_s$ is varied independently by doubling and halving the parameter. Each subfigure corresponds to variation of the parameter indicated in its title.

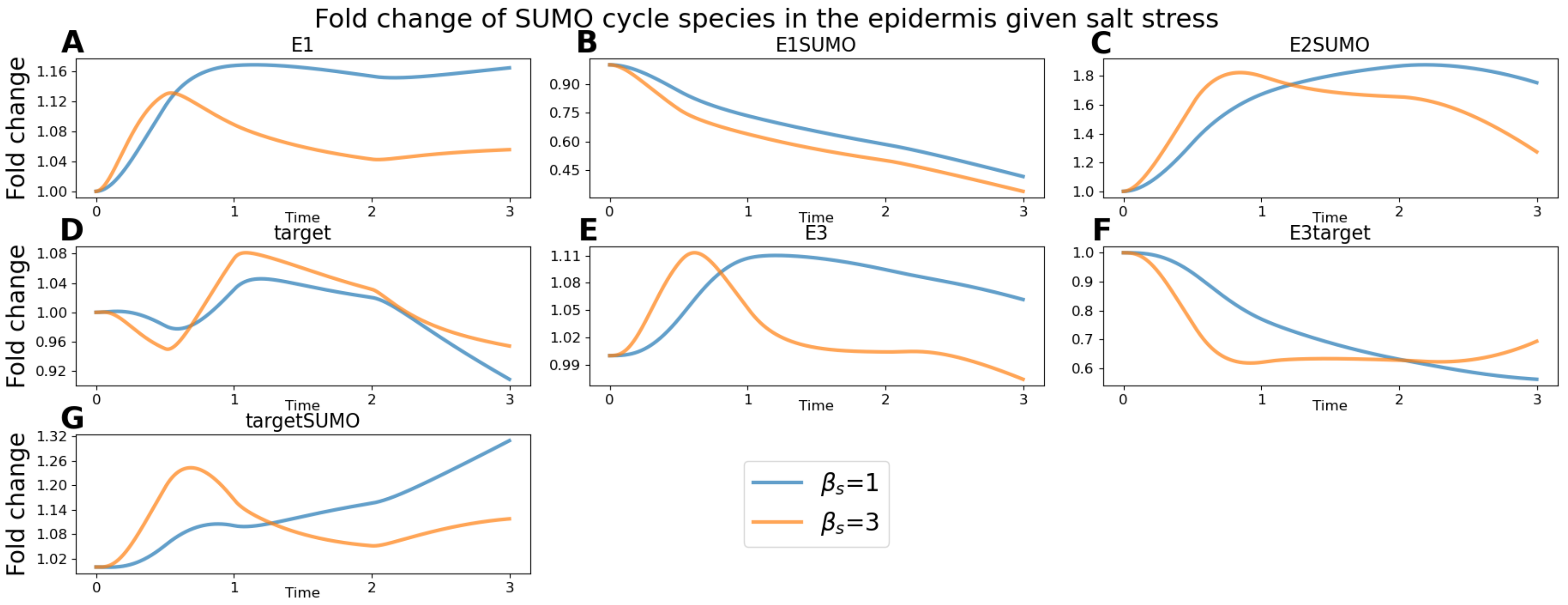


**Supplementary Figure S19. Predicted dynamics of the remaining SUMO cycle species after salt stress in the epidermis.** System Eq. (38) is initialized at the unstressed steady state under parameter sets $\chi_1$ and $\chi_3$. The means of the experimentally derived time courses for SUMO, ULP, E2, and DeSI given salt stress in the epidermis (Fig. 3) are imposed as input functions. In this figure, we see the predicted dynamics of the remaining species.

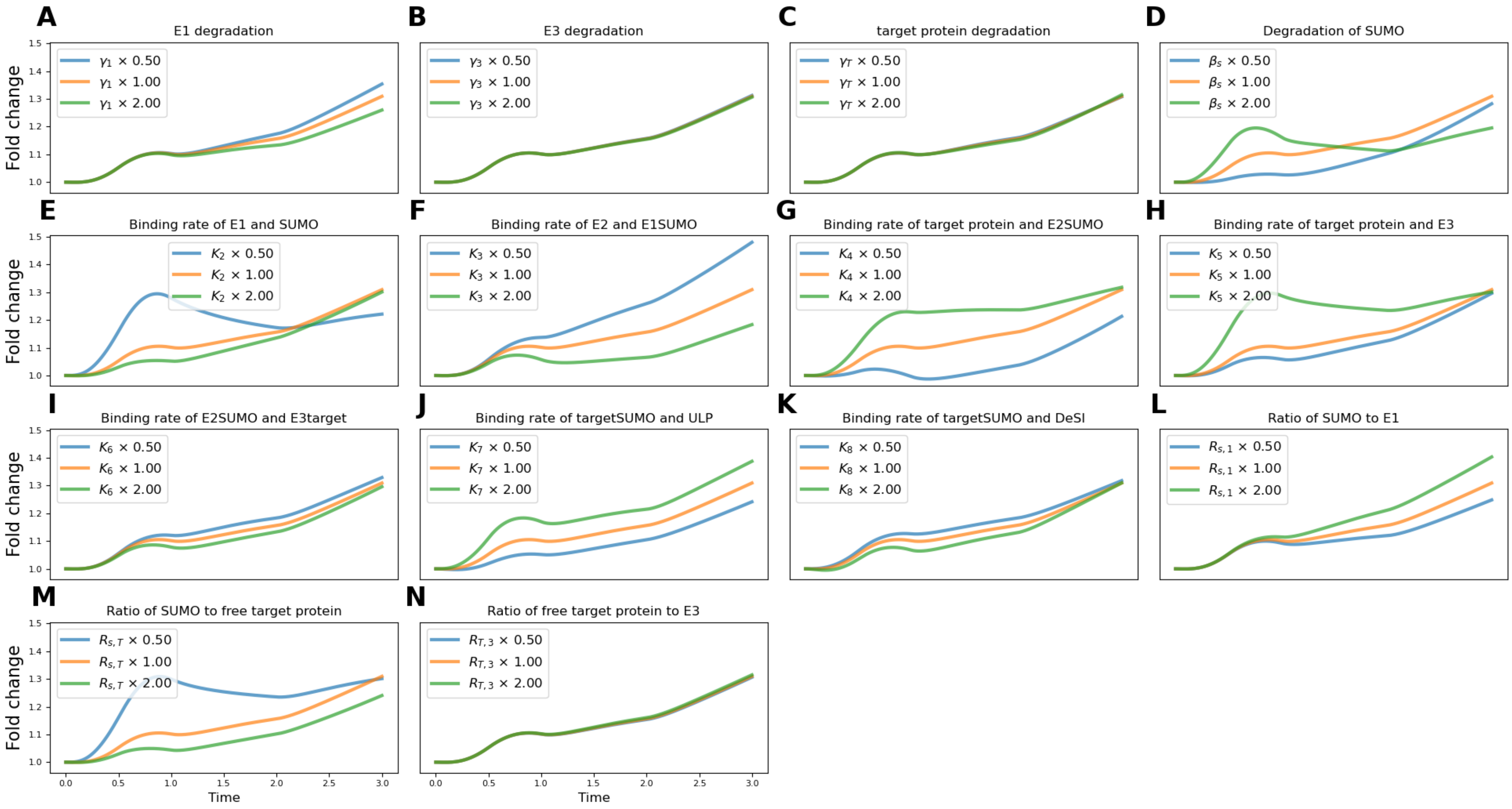


**Supplementary Figure S20. Sensitivity of SUMOylated target dynamics to parameter variation following salt stress in the epidermis given $\beta_s = 1$.** System Eq. (38) is initialized at the unstressed steady state under parameter set $\chi_1$. The means of the experimentally derived dynamics of SUMO, ULP, E2, and DeSI given salt stress in the epidermis (Fig. 3) are imposed as input functions. All parameters in $\chi$ are varied independently by doubling and halving the parameter. Each subfigure shows the resulting dynamics of SUMOylated target for the parameter indicated in the panel title.

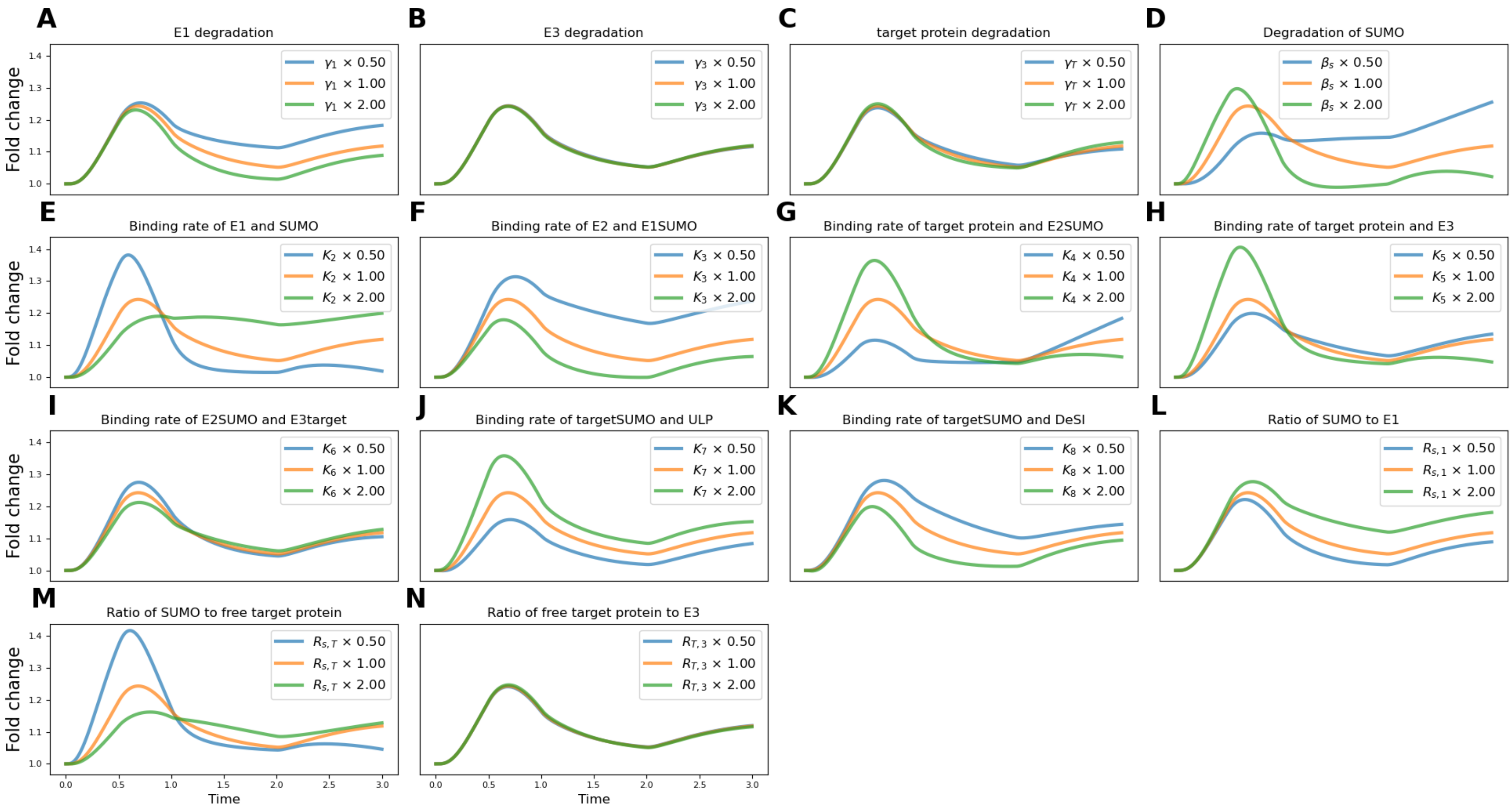


**Supplementary Figure S21. Sensitivity of SUMOylated target dynamics to parameter variation following salt stress in the epidermis given** $\beta_s = 3$**.** System $\mathrm{Eq.}\ (38)$ is initialized at the unstressed steady state under parameter set $\chi_3$. The means of the experimentally derived dynamics of SUMO, ULP, E2, and DeSI given salt stress in the epidermis (Fig. 3) are imposed as input functions. All parameters in $\chi$ are varied independently by doubling and halving the parameter. Each subfigure shows the resulting dynamics of SUMOylated target for the parameter indicated in the panel title.

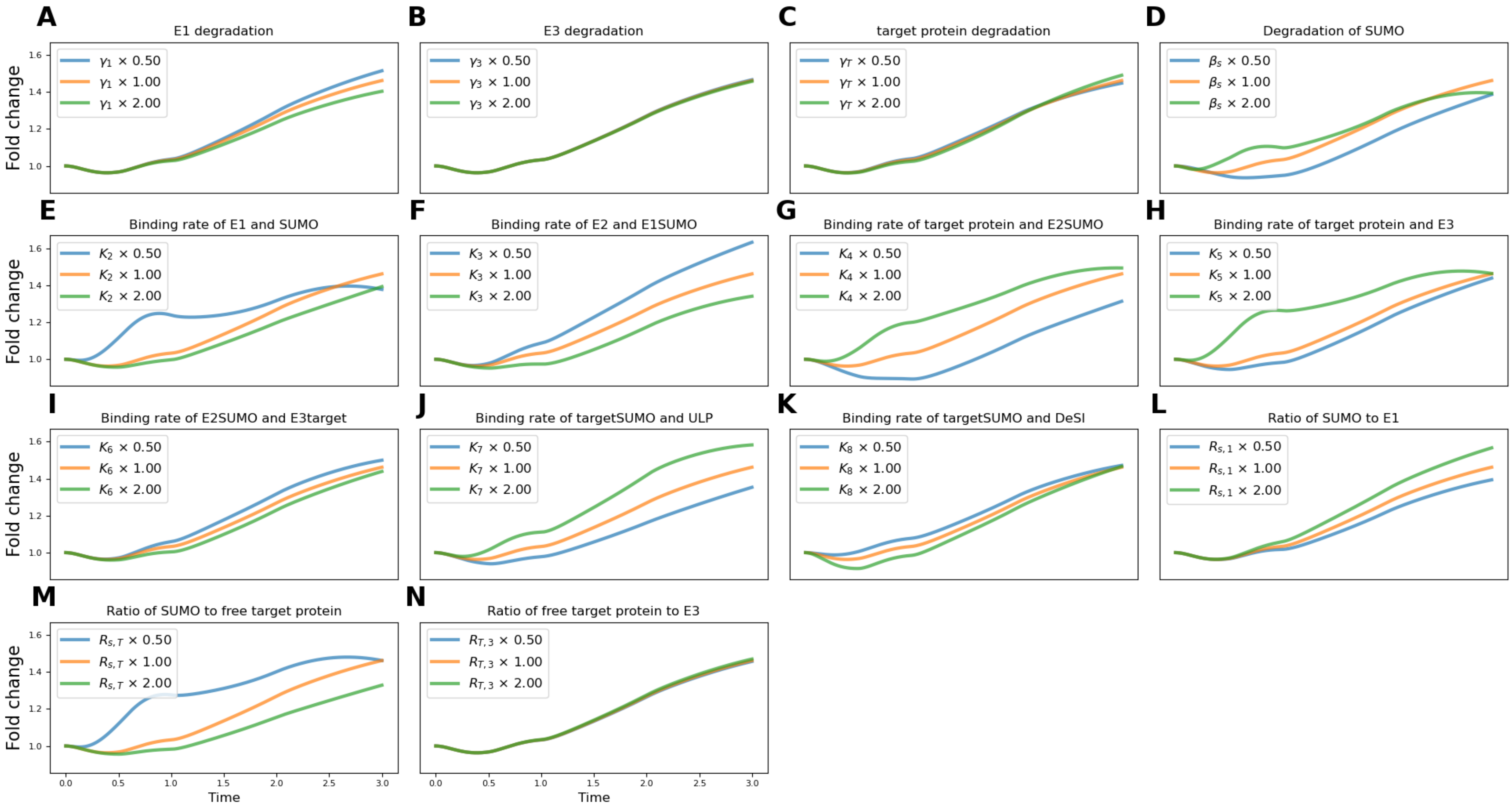


**Supplementary Figure S22. Sensitivity of SUMOylated target dynamics to parameter variation following salt stress in the stele given $\beta_s = 1$.** System Eq. (38) is initialized at the unstressed steady state under parameter set $\chi_1$. The means of the experimentally derived dynamics of SUMO, ULP, E2, and DeSI given salt stress in the stele (Fig. 3) are imposed as input functions. All parameters in $\chi$ are varied independently by doubling and halving the parameter. Each subfigure shows the resulting dynamics of SUMOylated target for the parameter indicated in the panel title.

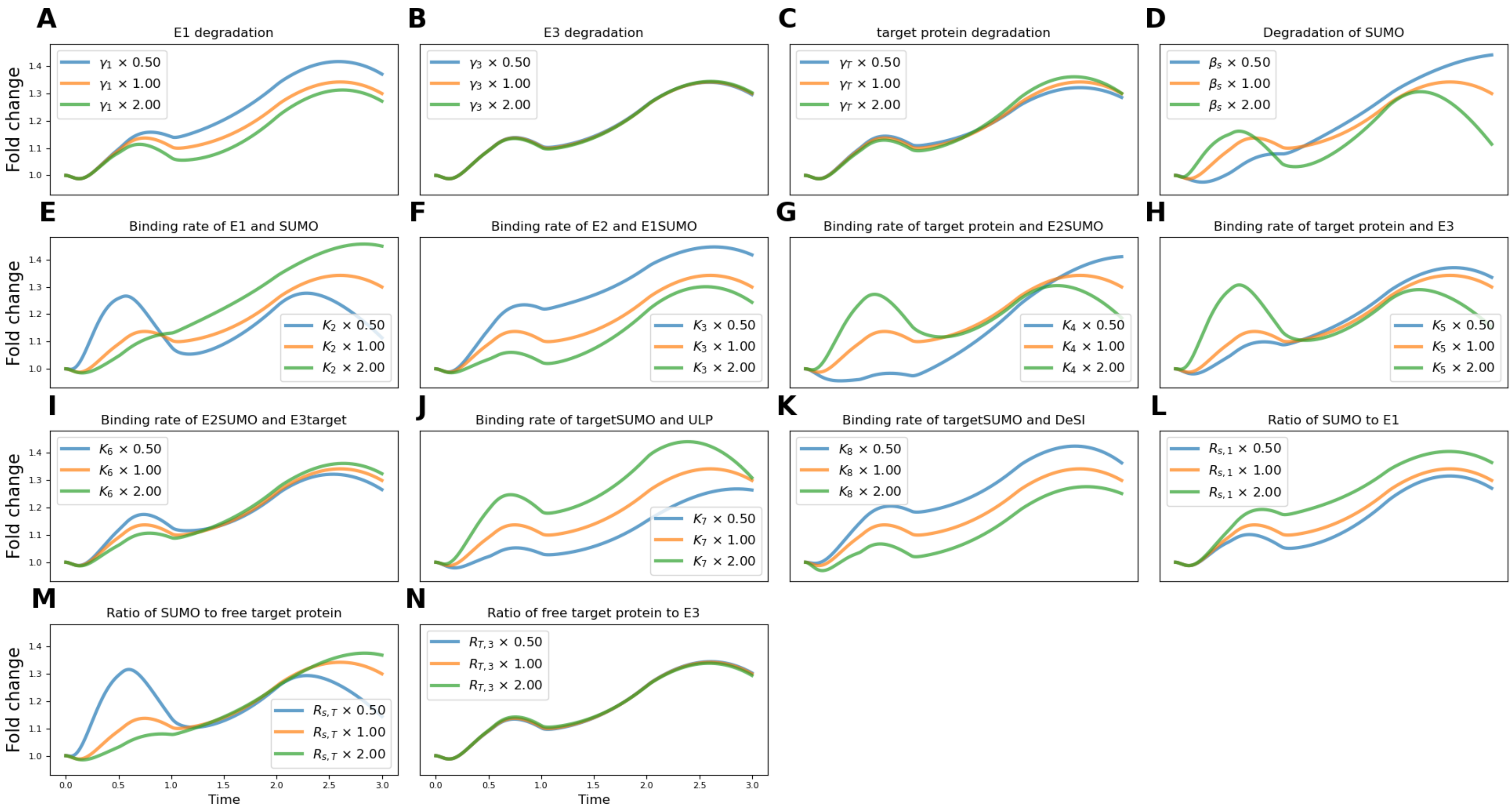


**Supplementary Figure S23. Sensitivity of SUMOylated target dynamics to parameter variation following salt stress in the stele given $\beta_s = 3$.** System Eq. (38) is initialized at the unstressed steady state under parameter set $\chi_3$. The means of the experimentally derived dynamics of SUMO, ULP, E2, and DeSI given salt stress in the stele (Fig. 3) are imposed as input functions. All parameters in $\chi$ are varied independently by doubling and halving the parameter. Each subfigure shows the resulting dynamics of SUMOylated target for the parameter indicated in the panel title.

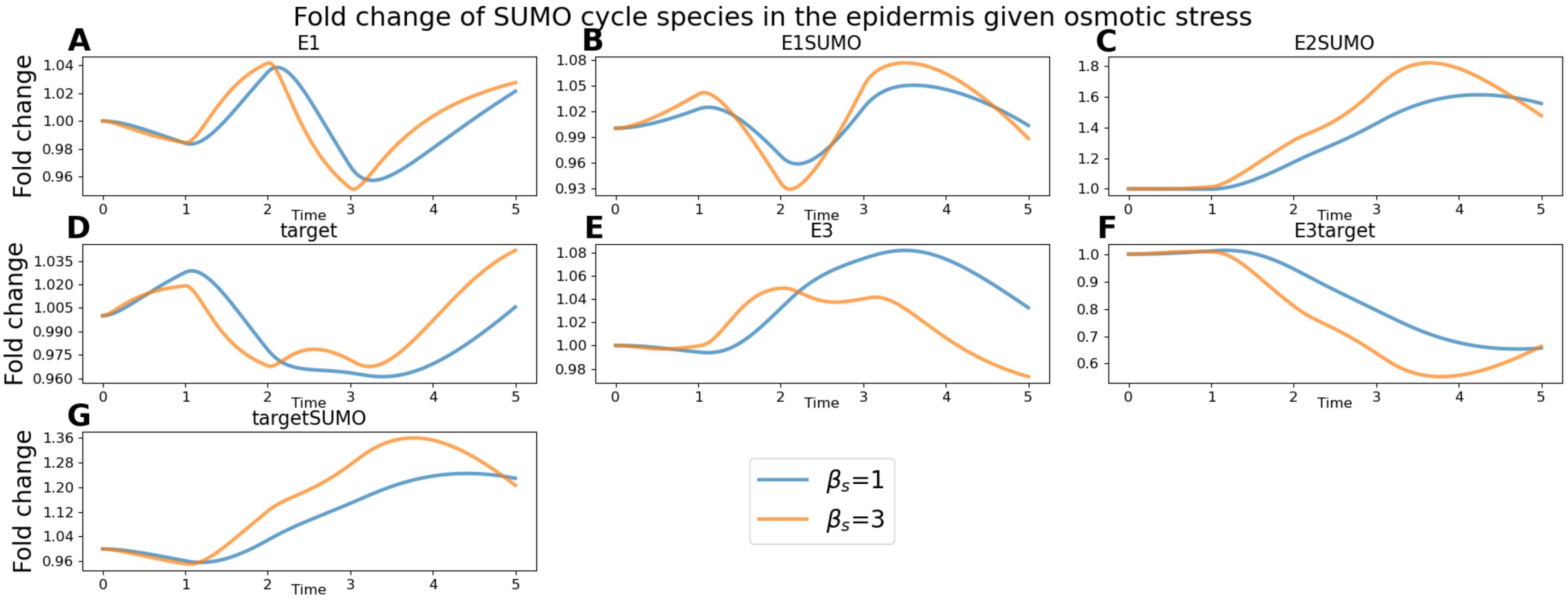


**Supplementary Figure S24. Predicted dynamics of the remaining SUMO cycle species after osmotic stress in the epidermis.** System Eq. (38) is initialized at the unstressed steady state under parameter sets $\chi_1$ and $\chi_3$. The means of the experimentally derived time courses for SUMO, ULP, E2, and DeSI given osmotic stress in the epidermis (Fig. 3) are imposed as input functions. In this figure, we see the predicted dynamics of the remaining SUMO cycle species.

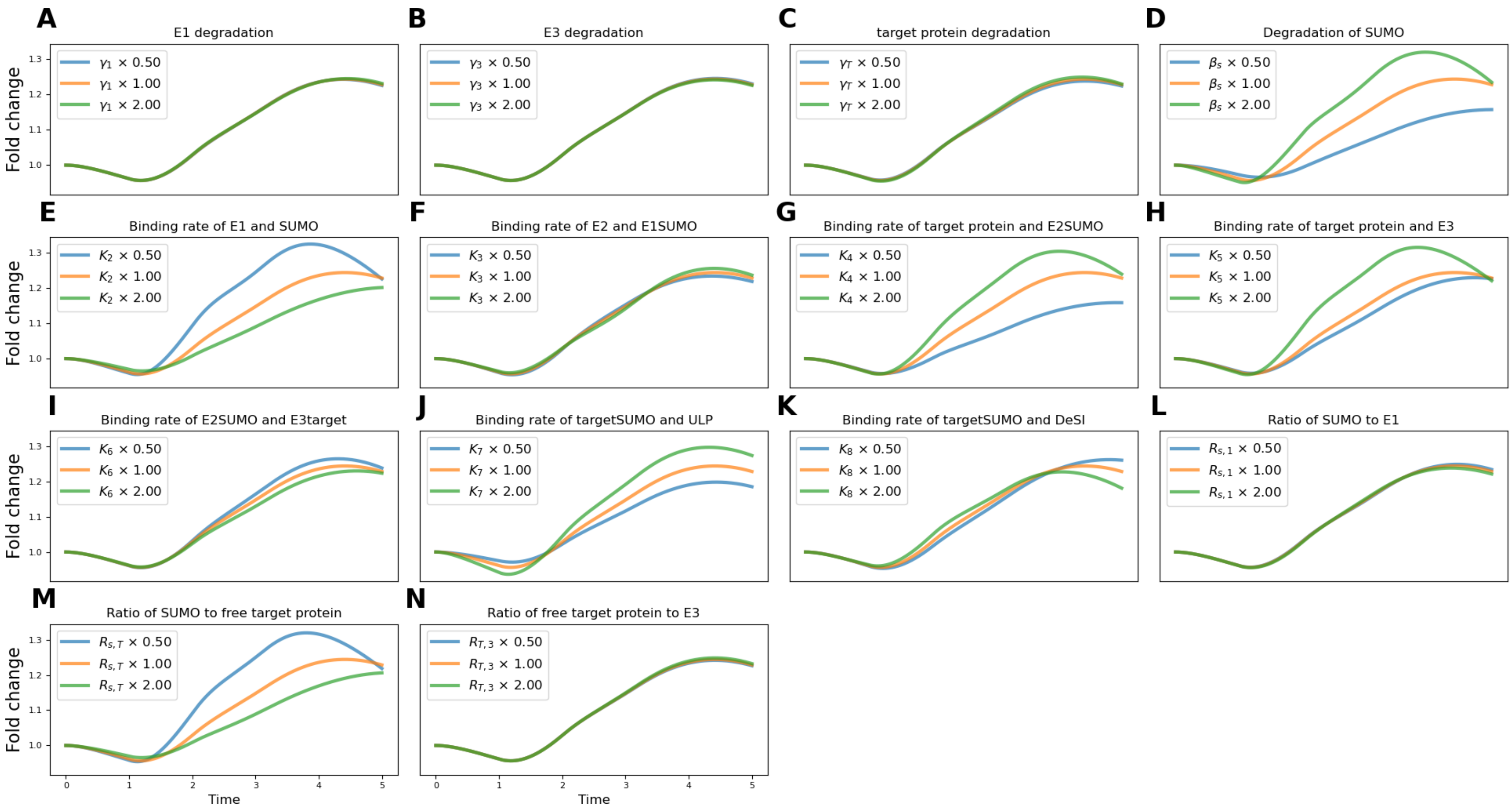


**Supplementary Figure S25. Sensitivity of SUMOylated target dynamics to parameter variation following osmotic stress in the epidermis given $\beta_s = 1$.** System Eq. (38) is initialized at the unstressed steady state under parameter set $\chi_1$. The means of the experimentally derived dynamics of SUMO, ULP, E2, and DeSI given osmotic stress in the epidermis (Fig. 3) are imposed as input functions. All parameters in $\chi$ are varied independently by doubling and halving the parameter. Each subfigure shows the resulting dynamics of SUMOylated target for the parameter indicated in the panel title.

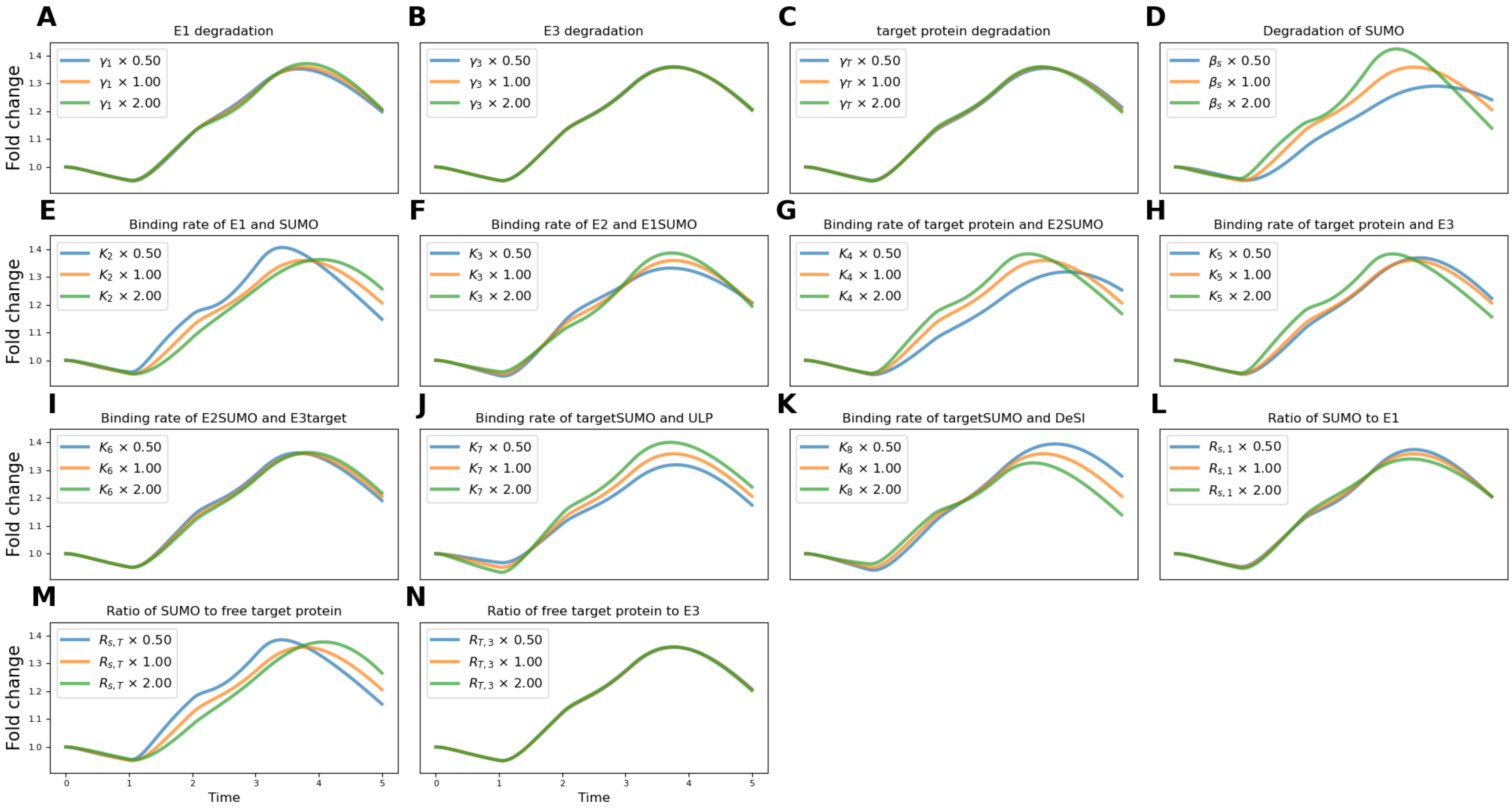


**Supplementary Figure S26. Sensitivity of SUMOylated target dynamics to parameter variation following osmotic stress in the epidermis given $\beta_s = 3$.** System $\mathrm{Eq.}\ (38)$ is initialized at the unstressed steady state under parameter set $\chi_3$. The means of the experimentally derived dynamics of SUMO, ULP, E2, and DeSI given osmotic stress in the epidermis (Fig. 3) are imposed as input functions. All parameters in $\chi$ are varied independently by doubling and halving the parameter. Each subfigure shows the resulting dynamics of SUMOylated target for the parameter indicated in the panel title.

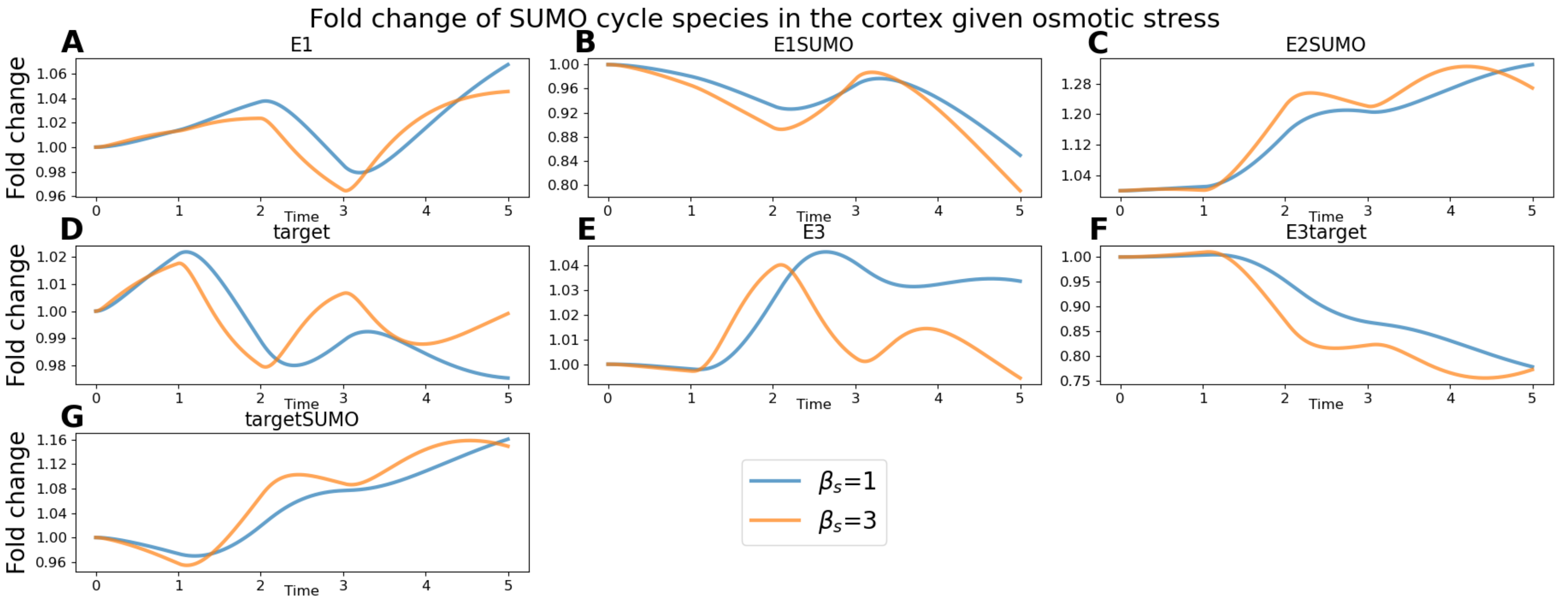


**Supplementary Figure S27. Predicted dynamics of the remaining SUMO cycle species after osmotic stress in the cortex.** System $\mathrm{Eq.}\ (38)$ is initialized at the unstressed steady state under parameter sets $\chi_1$ and $\chi_3$. The means of the experimentally derived time courses for SUMO, ULP, E2, and DeSI given osmotic stress in the cortex (Fig. 3) are imposed as input functions. In this figure, we see the predicted dynamics of the remaining SUMO cycle species.

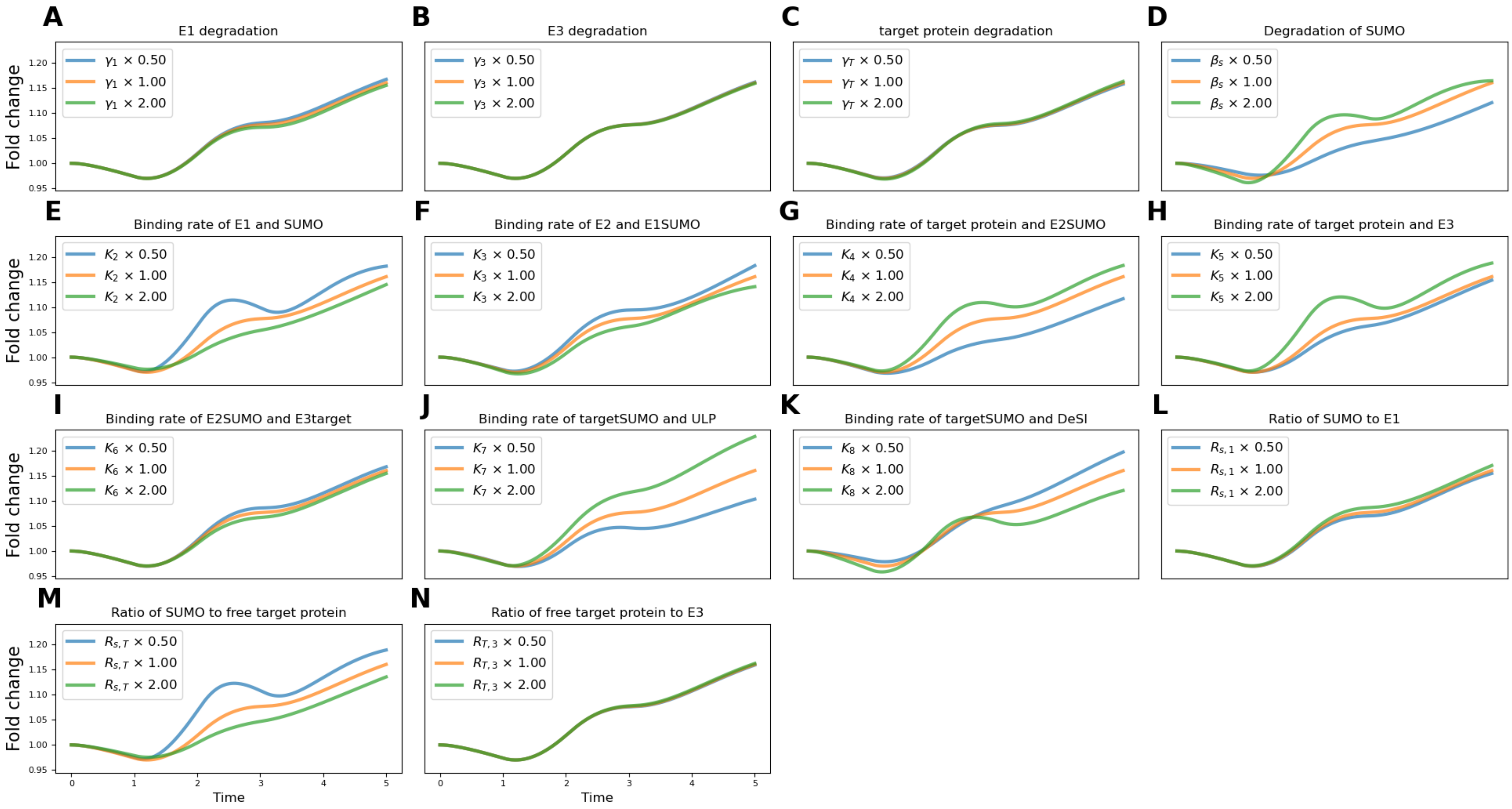


**Supplementary Figure S28. Sensitivity of SUMOylated target dynamics to parameter variation following osmotic stress in the cortex given $\beta_s = 1$.** System Eq. (38) is initialized at the unstressed steady state under parameter set $\chi_1$. The means of the experimentally derived dynamics of SUMO, ULP, E2, and DeSI given osmotic stress in the cortex (Fig. 3) are imposed as input functions. All parameters in $\chi$ are varied independently by doubling and halving the parameter. Each subfigure shows the resulting dynamics of SUMOylated target for the parameter indicated in the panel title.

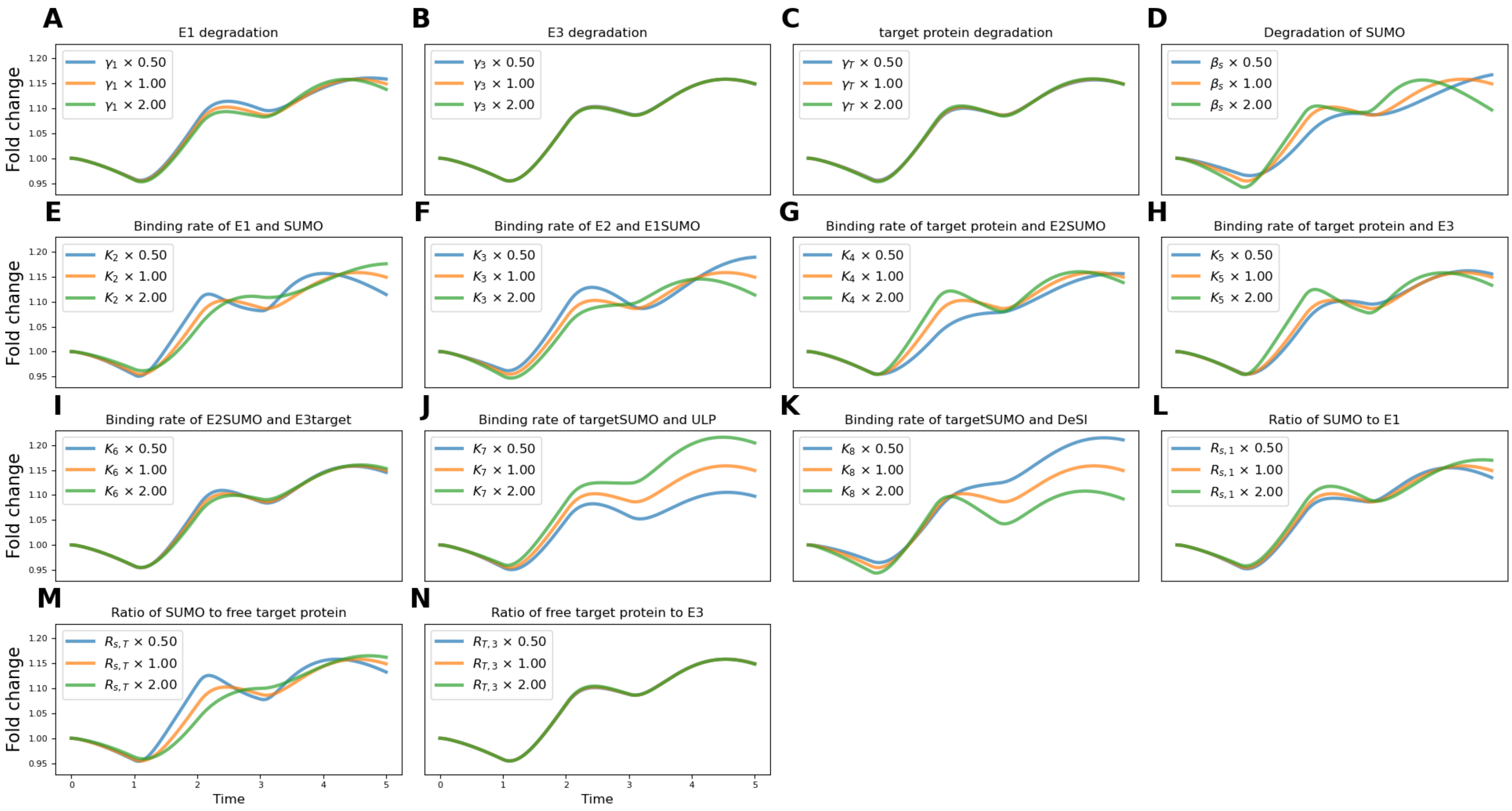


**Supplementary Figure S29. Sensitivity of SUMOylated target dynamics to parameter variation following osmotic stress in the cortex given $\beta_s = 3$.** System Eq. (38) is initialized at the unstressed steady state under parameter set $\chi_3$. The means of the experimentally derived dynamics of SUMO, ULP, E2, and DeSI given osmotic stress in the cortex (Fig. 3) are imposed as input functions. All parameters in $\chi$ are varied independently by doubling and halving the parameter. Each subfigure shows the resulting dynamics of SUMOylated target for the parameter indicated in the panel title.

### Supplementary tables

**Supplementary Table S1.** The SUMO Cell Atlas data (1) showing $\log_2$ fold changes of protein abundance of SUMO pathway species after salt stress across root tissues (rounded to 3 d.p.). We note that values close to zero correspond to little to no change in species levels after stress

| Family | Protein | Meristem | | | | Elongation Zone | | | |
|---|---|---|---|---|---|---|---|---|---|
| | | Epidermis | Cortex | Endodermis | Stele | Epidermis | Cortex | Endodermis | Stele |
| **SUMO** | SUMO2 | −0.192 | −0.412 | −0.280 | −0.099 | −0.519 | −0.596 | −0.518 | −0.570 |
| **E1** | SAE1a | 0.659 | 0.782 | 0.820 | 0.997 | 0.077 | 0.142 | 0.165 | 0.420 |
| | SAE1b | 0.186 | 0.554 | 0.505 | 0.340 | −0.210 | −0.210 | −0.109 | 0.099 |
| | SAE2 | −0.080 | −0.020 | −0.191 | 0.128 | −0.514 | −0.495 | −0.488 | −0.102 |
| **E2** | SCE1 | 0.679 | 0.427 | 0.074 | 0.229 | 0.996 | 0.818 | 1.080 | 1.296 |
| **E3** | SIZ1 | −1.099 | −1.200 | −1.206 | −1.497 | −0.817 | −0.569 | −0.846 | −0.780 |
| | HPY2 | 0.048 | 0.129 | 0.011 | 0.078 | NA | NA | NA | −0.214 |
| **E4** | PIAL1 | 0.103 | 0.088 | 0.272 | 0.600 | NA | NA | NA | NA |
| | PIAL2 | 0.532 | 0.456 | 0.640 | 0.386 | 0.205 | −0.298 | −0.449 | −0.059 |
| **ULP** | OTS1 | 0.081 | −0.152 | −0.094 | −0.103 | −0.048 | −0.214 | −0.376 | −0.197 |
| | OTS2 | 0.059 | −0.454 | 0.001 | 0.090 | −0.383 | −0.491 | −0.491 | −0.407 |
| | ESD4 | −0.230 | −0.277 | −0.363 | −0.269 | 0.132 | −0.222 | −0.037 | −0.177 |
| | ELS1 | 0.282 | 0.189 | 0.351 | 0.293 | 0.408 | 0.229 | 0.257 | 0.106 |
| | ELS2 | NA | NA | NA | NA | NA | NA | NA | NA |
| | FUG1 | NA | NA | NA | 0.008 | NA | NA | NA | 0.083 |
| | SPF1 | 0.743 | 0.660 | 1.181 | 0.958 | 0.550 | 0.010 | 0.439 | 0.475 |
| | SPF2 | −0.890 | −0.699 | −0.664 | −0.503 | −0.446 | −0.805 | −0.749 | −0.268 |
| **DeSI** | DeSI1 | −0.391 | −0.494 | −0.356 | −0.467 | −0.218 | −0.318 | −0.256 | −0.318 |
| | DeSI2a | 0.169 | −0.033 | 0.113 | 0.194 | −0.044 | −0.133 | 0.081 | 0.175 |
| | DeSI2b | 0.140 | 0.132 | 0.083 | 0.274 | −0.076 | −0.344 | 0.021 | −0.083 |
| | DeSI3a | 0.091 | 0.127 | −0.049 | −0.002 | 0.449 | 0.451 | 0.568 | 0.467 |
| | DeSI3b | 0.231 | 0.186 | 0.221 | 0.354 | 0.006 | 0.317 | 0.295 | 0.377 |
| | DeSI3c | 0.296 | 0.425 | NA | 0.242 | 0.044 | −0.131 | 0.462 | 0.606 |
| | DeSI4a | −0.118 | −0.240 | −0.209 | −0.057 | −0.647 | −1.266 | −0.978 | −0.609 |
| | DeSI4b | −0.014 | 0.061 | 0.023 | 0.136 | −0.050 | 0.048 | −0.341 | −0.173 |

**Supplementary Table S2.** The SUMO Cell Atlas data (1) showing $\log_2$ fold changes of protein abundance of SUMO pathway species after osmotic stress across root tissues (rounded to 3 d.p.). We note that values close to zero correspond to little to no change in species levels after stress.

| Family | Protein | Meristem | | | | Elongation Zone | | | |
|---|---|---|---|---|---|---|---|---|---|
| | | Epidermis | Cortex | Endodermis | Stele | Epidermis | Cortex | Endodermis | Stele |
| **SUMO** | SUMO2 | 0.236 | 0.284 | 0.180 | 0.228 | 0.515 | 0.215 | 0.229 | 0.385 |
| **E1** | SAE1a | 0.260 | 0.221 | 0.080 | 0.179 | 0.279 | 0.236 | 0.490 | 0.328 |
| | SAE1b | −0.355 | −0.278 | −0.236 | −0.089 | 0.055 | 0.011 | −0.061 | 0.086 |
| | SAE2 | −0.064 | 0.024 | 0.013 | 0.063 | 0.059 | −0.120 | −0.004 | −0.119 |
| **E2** | SCE1 | 0.223 | 0.241 | 0.180 | 0.413 | 0.525 | 0.529 | 0.713 | 0.800 |
| **E3** | SIZ1 | 0.210 | 0.503 | 0.608 | 0.437 | 0.046 | 0.039 | −0.012 | −0.072 |
| | HPY2 | 0.108 | 0.151 | 0.125 | 0.147 | NA | NA | NA | −0.270 |
| **E4** | PIAL1 | −0.059 | 0.217 | 0.185 | 0.190 | NA | NA | NA | NA |
| | PIAL2 | 0.437 | 0.323 | 0.365 | 0.393 | −0.086 | −0.078 | 0.022 | 0.171 |
| **ULP** | OTS1 | 0.178 | 0.196 | 0.150 | 0.128 | −0.066 | −0.196 | −0.074 | −0.164 |
| | OTS2 | 0.528 | 0.596 | 0.630 | 0.545 | −0.006 | −0.146 | 0.064 | 0.161 |
| | ESD4 | 0.169 | 0.119 | 0.252 | 0.278 | −0.041 | 0.070 | −0.060 | 0.031 |
| | ELS1 | 0.069 | 0.434 | 0.276 | 0.107 | −0.333 | 0.045 | −0.123 | −0.017 |
| | ELS2 | NA | NA | NA | NA | NA | NA | NA | NA |
| | FUG1 | NA | NA | NA | 0.097 | NA | NA | NA | −0.375 |
| | SPF1 | 0.152 | 0.288 | 0.402 | 0.126 | −0.387 | −0.160 | 0.063 | 0.122 |
| | SPF2 | 0.127 | 0.079 | 0.109 | 0.071 | −0.036 | 0.076 | 0.182 | −0.020 |
| **DeSI** | DeSI1 | 0.143 | 0.417 | 0.506 | 0.422 | 0.139 | 0.273 | 0.108 | 0.310 |
| | DeSI2a | 0.591 | 0.529 | 0.551 | 0.556 | 0.611 | 0.526 | 0.514 | 0.614 |
| | DeSI2b | 0.450 | 0.639 | 0.479 | 0.469 | 0.303 | 0.493 | 0.637 | 0.618 |
| | DeSI3a | −0.003 | 0.057 | −0.054 | 0.127 | −0.474 | −0.313 | −0.298 | −0.252 |
| | DeSI3b | 0.124 | 0.282 | 0.253 | 0.041 | −0.095 | 0.090 | 0.050 | −0.128 |
| | DeSI3c | 0.403 | NA | NA | 0.257 | 0.366 | NA | NA | −0.071 |
| | DeSI4a | 0.138 | 0.177 | 0.151 | 0.054 | 0.138 | 0.113 | 0.076 | −0.160 |
| | DeSI4b | 0.069 | 0.047 | 0.129 | 0.066 | −0.058 | 0.048 | −0.005 | 0.163 |

**Supplementary Table S3.** Fluorescence intensities of SUMO pathway isoforms and their proportional contribution to total fluorescence in the meristem under baseline conditions.

| Family | Protein | Fluorescence | Proportion of total |
|---|---|---|---|
| **SUMO** | SUMO2 | 3 746 376 | 1.000 |
| **SAE1** | SAE1a | 13 262 550 | 0.518 |
| | SAE1b | 12 348 281 | 0.482 |
| **SAE2** | SAE2 | 8 442 102 | 1.000 |
| **E2** | SCE1 | 15 295 035 | 1.000 |
| **E3** | SIZ1 | 2 735 313 | 0.597 |
| | HPY2 | 1 847 973 | 0.403 |
| **E4** | PIAL1 | 346 042 | 0.103 |
| | PIAL2 | 3 021 594 | 0.897 |
| **ULP** | OTS1 | 3 085 234 | 0.279 |
| | OTS2 | 365 879 | 0.033 |
| | ESD4 | 2 194 027 | 0.199 |
| | ELS1 | 1 625 151 | 0.147 |
| | ELS2 | 309 341 | 0.028 |
| | FUG1 | 338 925 | 0.031 |
| | SPF1 | 2 863 955 | 0.259 |
| | SPF2 | 269 575 | 0.024 |
| **DeSI** | DeSI1 | 9 254 297 | 0.259 |
| | DeSI2a | 7 537 367 | 0.211 |
| | DeSI2b | 5 477 298 | 0.153 |
| | DeSI3a | 1 422 308 | 0.040 |
| | DeSI3b | 2 333 479 | 0.065 |
| | DeSI3c | 1 486 345 | 0.042 |
| | DeSI4a | 4 069 479 | 0.114 |
| | DeSI4b | 4 202 004 | 0.117 |

### References

1. Jason Banda, Shraboni Ghosh, Dipan Roy, Kishor D. Ingole, Lisa Clark, Eshan Sharma, Sumesh Kakkunath, Kawinnat Sue-Ob, Rahul Bhosale, Leah Band, Srayan Ghosh, Darren Wells, Jonathan

Atkinson, Nicholas J. Provart, Malcolm J. Bennett, Kathryn S. Lilley, Andrew Jones, Miguel De Lucas, Anthony Bishopp, and Ari Sadanandom. Elucidating tissue and subcellular specificity of the entire SUMO network reveals how stress responses are fine-tuned in a eukaryote. *Science Advances*, 11(35): eadw9153, 2025. . URL https://www.science.org/doi/abs/10.1126/sciadv.adw9153.

2. Lucio Conti, Gillian Price, Elizabeth O'Donnell, Benjamin Schwessinger, Peter Dominy, and Ari Sadanandom. Small ubiquitin-like modifier proteases OVERLY TOLERANT to SALT1 and -2 regulate salt stress responses in Arabidopsis. *The Plant Cell*, 20(10):2894–2908, 10 2008. ISSN 1040-4651. . URL https://doi.org/10.1105/tpc.108.058669.
3. Marcus J. Miller, Gregory A. Barrett-Wilt, Zhihua Hua, and Richard D. Vierstra. Proteomic analyses identify a diverse array of nuclear processes affected by small ubiquitin-like modifier conjugation in Arabidopsis. *Proceedings of the National Academy of Sciences*, 107(38):16512–16517, 2010. . URL https://www.pnas.org/doi/abs/10.1073/pnas.1004181107.
4. Hyeong Cheol Park, Wonkyun Choi, Hee Jin Park, Mi Sun Cheong, Yoon Duck Koo, Gilok Shin, Woo Sik Chung, Woe-Yeon Kim, Min Gab Kim, Ray A. Bressan, Hans J. Bohnert, Sang Yeol Lee, and Dae-Jin Yun. Identification and molecular properties of SUMO-binding proteins in Arabidopsis. *Molecules and Cells*, 32(2):143–152, 2011. ISSN 1016-8478. . URL https://www.sciencedirect.com/science/article/pii/S1016847823109381.
5. Rebecca Morrell and Ari Sadanandom. Dealing with stress: A review of plant SUMO proteases. *Frontiers in Plant Science*, 10, 2019. ISSN 1664-462X. . URL https://www.frontiersin.org/journals/plant-science/articles/10.3389/fpls.2019.01122.
6. Pauli Virtanen, Ralf Gommers, Travis E. Oliphant, Matt Haberland, Tyler Reddy, David Cournapeau, Evgeni Burovski, Pearu Peterson, Warren Weckesser, Jonathan Bright, Stéfan J. van der Walt, Matthew Brett, Joshua Wilson, K. Jarrod Millman, Nikolay Mayorov, Andrew R. J. Nelson, Eric Jones, Robert Kern, Eric Larson, C J Carey, İlhan Polat, Yu Feng, Eric W. Moore, Jake VanderPlas, Denis Laxalde, Josef Perktold, Robert Cimrman, Ian Henriksen, E. A. Quintero, Charles R. Harris, Anne M. Archibald, Antônio H. Ribeiro, Fabian Pedregosa, Paul van Mulbregt, and SciPy 1.0 Contributors. SciPy 1.0: Fundamental Algorithms for Scientific Computing in Python. *Nature Methods*, 17:261–272, 2020. .
